# Gene genealogies in haploid populations evolving according to sweepstakes reproduction

**DOI:** 10.64898/2026.01.08.698389

**Authors:** Bjarki Eldon

## Abstract

Recruitment dynamics, or the distribution of the number of offspring among individuals, is fundamental to ecology and evolution. We take sweepstakes reproduction to mean a skewed (heavy right-tailed) offspring number distribution without natural selection being involved. Sweepstakes may be generated by chance matching of reproduction with favorable environmental conditions. Gene genealogies generated by sweepstakes reproduction are in the domain of attraction of multiple-merger coalescents where a random number of lineages merges at such times. We consider population genetic models of sweepstakes reproduction for haploid panmictic populations of both constant (*N*), and varying population size, and evolving in a random environment. We construct our models so that we can recover the observed number of new mutations in a given sample without requiring strong assumptions regarding the population size or the mutation rate. Our main results are *(i)* continuous-time coalescents that are either the Kingman coalescent or specific families of Beta- or Poisson-Dirichlet coalescents; when combining the results the parameter *α* of the Beta-coalescent ranges from 0 to 2, and the Beta-coalescents may be incomplete due to an upper bound on the number of potential offspring an arbitrary individual may produce; *(ii)* in large populations we measure time in units proportional to either *N/* log *N* or *N* generations; *(iii)* incorporating fluctuations in population size leads to time-changed multiple-merger coalescents where the time-change does not depend on *α*; *(iv)* using simulations we show that in some cases approximations of functionals of a given coalescent do not match the ones of the ancestral process in the domain of attraction of the given coalescent; *(v)* approximations of functionals obtained by conditioning on the population ancestry (the ancestral relations of all gene copies at all times) are broadly similar (for the models considered here) to the approximations obtained without conditioning on the population ancestry.

## 1 Introduction

Inferring evolutionary histories of natural populations is one of the main aims of population genetics. Inheritance, or the transfer of genetic information from a parent to an offspring, is the characteristic of organisms that makes inference possible. Inheritance of ancestral gene copies leaves a (unknown) ‘trail’ of ancestral relations; in other words gene copies are connected by a gene tree, or gene genealogy. The probabilistic modeling of the random ancestral relations of sampled gene copies is one way to approximate the trail going backwards in time to learn about the past events that shaped observed genetic variation. The mathematical study of random gene genealogies generates a framework for constructing inference methods [Donnelly and Tavaré, 1995, Wakeley, 2009, Berestycki, 2009]. Absent natural selection and complex demography the shape or structure of gene genealogies, and so predictions about genetic variation, will primarily be influenced by the offspring number distribution. The importance of the offspring number distribution for understanding evolution and ecology of natural populations is clear.

Gene genealogies in populations where one can ignore the potential occurrence of large (proportional to the population size) number of offspring per individual in a sufficiently large population, are in the domain of attraction of the Kingman coalescent [Kingman, 1982b,a]. In population genetics a coalescent is a probabilistic description of the random gene genealogies (random ancestral relations) of sampled gene copies. In the Kingman coalescent at most two gene copies ancestral to the sampled ones reach a common ancestor at the same time assuming the number of sampled gene copies is ‘small enough’ [Wakeley and Takahashi, 2003, Bhaskar et al., 2014, Melfi and Viswanath, 2018b,a]. The classical Wright-Fisher model (corresponding to offspring choosing a parent independently and uniformly at random and with replacement; Fisher [1923], Wright [1931]), the standard reference model in population genetics, is one such population model. However, it may not be a good choice for highly fecund populations characterised by sweepstakes reproduction [Eldon, 2020, Hedgecock and Pudovkin, 2011].

For understanding the ecology and evolution of highly fecund populations the question of sweepstakes reproduction is central [Hedgecock and Pudovkin, 2011, Árnason et al., 2023]. We say that a population evolves according to sweepstakes reproduction [Hedgecock et al., 1982, Hedgecock and Pudovkin, 2011, Hedgecock, 1994, Beckenbach, 1994] when natural selection is not involved in mechanisms generating a skewed offspring number distribution. The chance matching of reproduction with favorable environmental conditions in populations characterised by Type III survivorship may be one such mechanism. Sweepstakes reproduction has also been referred to as random sweepstakes [Árnason et al., 2023]. In highly fecund populations, in particular ones characterised by Type III survivorship, individuals may have the capacity to produce numbers of offspring proportional to the population size, even in a large population. Such events may affect the evolution of the population should they occur often enough. Processes tracking type frequencies in populations characterised by sweepstakes reproduction are in the domain of attraction of Fleming-Viot measure-valued jump diffusions [Fleming and Viot, 1979, Ethier and Kurtz, 1993, Birkner and Blath, 2009]; and gene genealogies of gene copies in samples from such populations are in the domain of attraction of multiple-merger coalescents, where the number of ancestral lineages involved when mergers occur is random [Sagitov, 2003, 1999, Donnelly and Kurtz, 1999, Pitman, 1999, Schweinsberg, 2000]. Multiple-merger coalescents also result from recurrent strong bottlenecks [Birkner et al., 2009, Eldon and Wakeley, 2006], and strong positive selection resulting in recurrent selective sweeps [Durrett and Schweinsberg, 2005]. Multiple-merger coalescents predict patterns of genetic diversity different from the ones predicted by the King-man coalescent [Blath et al., 2016, Birkner et al., 2013b, Sargsyan and Wakeley, 2008, Eldon and Wakeley, 2006], and may be essential for explaining genetic diversity and evolutionary histories across a broad spectrum of highly fecund natural populations [Hedgecock and Pudovkin, 2011, Beckenbach, 1994, Árnason, 2004, Eldon, 2020, Árnason and Halldórsdóttir, 2015, Barfield et al., 2023, Vendrami et al., 2021, CHRISTIE et al., 2010, Árnason et al., 2023, Hedgecock et al., 2007]. This motivates us, and it is the aim here, to investigate gene genealogies based on extensions of current models [Schweinsberg, 2003, Chetwynd-Diggle and Eldon, 2026] of sweepstakes reproduction.

Population genetic models of sweepstakes reproduction for haploid and diploid populations have been rigorously formulated and studied [Sagitov, 1999, Möhle and Sagitov, 2001, Schweinsberg, 2003, Sargsyan and Wakeley, 2008, Möhle and Sagitov, 2003, Birkner et al., 2018, Huillet and Möhle, 2013]. In particular, Eldon and Wakeley [2006] and Birkner et al. [2013a] consider populations evolving according to sweepstakes reproduction in a random environment. Eldon and Wakeley [2006] consider a haploid Moran and Wright-Fisher-type models, in which, with probability *p*_*N*_ (where *p*_*N*_ → 0 as *N* → ∞ where *N* is the population size of a haploid panmictic population), a single individual contributes a fixed fraction of the total population, and the remaining fraction of the offspring choose a parent from the other *N* − 1 available parents independently and uniformly at random and with replacement; with probability 1 − *p*_*N*_ ordinary Wright-Fisher reproduction occurs. Huillet and Möhle [2013] extend the Moran model to be in the domain of attraction of multiple-merger coalescents. Birkner et al. [2013a] construct ancestral recombination graphs admitting simultaneous mergers based on a diploid version of the model studied by Eldon and Wakeley [2006].

Here we investigate coalescents derived from population genetic models of sweepstakes reproduction in a haploid panmictic population evolving in a random environment. This article contributes *(i)* continuous-time coalescents that are either the Kingman coalescent or specific families of multiple-merger coalescents and with time in arbitrarily large populations measured in units proportional to either *N/* log *N* or *N* generations; *(ii)* numerical comparisons of functionals of coalescents and pre-limiting ancestral processes in the domain of attraction of the given coalescent; *(iii)* comparisons of functionals of ancestral processes when conditioning resp. not conditioning on the population ancestry. Here, ‘population ancestry’ refers to a record of the ancestral relations of the entire population from the present and infinitely far into the past.

To briefly summarize the results, we extend a model of the evolution (absent selection) of a haploid panmictic population evolving in a constant environment studied by Chetwynd-Diggle and Eldon [2026] to a random environment. In the random environment, most of the time (i.e. with high probability) individuals produce ‘small’ numbers of potential offspring, but occasionally the environment turns favorable (there is an increased chance) for producing a significant (at least on the order of the population size) numbers of potential offspring. The probability of producing *k* numbers of potential offspring is assumed to decay roughly like *k*^−*a*^ − (*k* + 1)^−*a*^ for *k* = 1, 2, …, *ζ*(*N*) for some given *a >* 0 and *ζ*(*N*) *>* 0. The quantity *ζ*(*N*) is taken to be an upper bound on the number of potential offspring any given individual can produce. Depending on how *ζ*(*N*)*/N* behaves in a large population (*N* → ∞), we obtain specific examples of continuous-time multiple-merger coalescents (Beta(*γ*; 2 − *α, α*)- and Poisson-Dirichlet(*α*, 0)-coalescents with an atom at zero). In contrast to Beta- and Poisson-Dirichlet-coalescents without an atom at zero [Schweinsberg, 2003], time in our models is measured in units proportional to (at least) *N/* log *N* generations, where *N* is the population size (in a constant-size population). Combining the results gives a Beta(*γ*; 2 − *α, α*)-coalescent where 0 *< α <* 2 and 0 *< γ* ≤ 1 is a truncation parameter (cf. also [Chetwynd-Diggle and Eldon, 2026]). We have extended the range of the *α* parameter of the Beta(2 − *α, α*)-coalescent introduced by Schweinsberg [2003]. Using simulations we show that including an atom at zero has implications for the effect of population growth on gene genealogies, in the sense that the effects do not vanish as the skewness of the offspring distribution increases (*α* decreases), in contrast to what has been observed when applying population growth to gene genealogies governed by Beta-coalescents without an atom at zero [Freund, 2020]. We use simulations to compare coalescent trees with the pre-limiting gene genealogies (using relative branch lengths); the agreement, especially when the probability distribution for the numbers of potential offspring is highly skewed, is poor. We also use simulations to compare gene genealogies obtained by conditioning resp. not conditioning on the population ancestry, a record of the ancestral relations of the gene copies in the population. The idea is that the complete sample gene genealogy of the gene copies in any given sample is, for a given population ancestry and once the identity of the sampled gene copies is known, fixed. The simulation results indicate that the two approaches lead to different predictions about patterns of genetic variation in a sample.

In § 2 we provide a brief background to coalescent processes and to models of sweep-stakes, in § 3 we state our main results given in Theorems 3.5 and 3.6 for constant population size (see § 3.1), in Theorems 3.8 and 3.9 (see § 3.3) concerning varying population size; in § 3.2 we give numerical examples comparing functionals of stated processes, § 4 contains concluding remarks. Proofs of our mathematical results may be found in § 5. A brief discussion of the Beta(2 − *β, β*)-Poisson-Dirichlet(*α*, 0)-coalescent may be found in Appendix A. Appendices B–F contain further numerical examples.

## 2 Background

For ease of reference we state standard notation used throughout.

### Definition 2.1

(Standard notation). *Write* ℕ ≡ {1, 2, …}, ℕ_0_ ≡ ℕ ∪ {0}. *Let N* ∈ ℕ *denote the population size (when constant). Asymptotics refer to the ones in an arbitrarily large population, i.e. taking N* → ∞, *unless otherwise noted. Write* [*n*] ≡ {1, 2, …, *n*} *for any n* ∈ ℕ. *For x* ∈ ℝ *(the set of reals) and m* ∈ ℕ_0_ *recall the falling factorial*

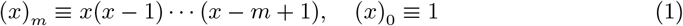

*For positive sequences* (*x*_*n*_)_*n*_ *and* (*y*_*n*_)_*n*_ *we write x*_*n*_ ∈ *o*(*y*_*n*_) *if* lim sup_*n*→∞_ *x*_*n*_*/y*_*n*_ = 0, *and x*_*n*_ ∈ *O*(*y*_*n*_) *if* lim sup_*n*→∞_ *x*_*n*_*/y*_*n*_ *<* ∞ *(so that o*(*y*_*n*_) ⊂ *O*(*y*_*n*_)*), and x*_*n*_ ~ *y*_*n*_ *if* lim_*n*→∞_ *x*_*n*_*/y*_*n*_ = 1 *(assuming y*_*n*_ *>* 0 *for all n). We extend the notation x*_*n*_ ~ *y*_*n*_ *for asymptotic equivalence and write*

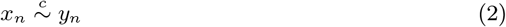

*if* lim_*n*→∞_ *x*_*n*_*/y*_*n*_ = *c for some constant c >* 0 *that may change depending on the context (when we use this notation we simply intend to emphasize the conditions under which* (2) *holds given* (*x*_*n*_) *and* (*y*_*n*_)*); K, C, C*^′^, *c, c*^′^ *will denote unspecified positive constants. Define*

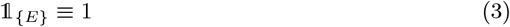

*if a given condition/event E holds, and 1*_{*E*}_ ≡ 0 *otherwise. The abbreviation i.i.d. will stand for independent and identically distributed (random variables)*.

The introduction of the coalescent [Kingman, 1982b,a, Hudson, 1983, Tajima, 1983] advanced the field of population genetics. A coalescent {*ξ*} ≡ {*ξ*(*t*); *t* ≥ 0} is a Markov chain taking values in the partitions of ℕ, such that the restriction {*ξ*^*n*^} ≡ {*ξ*^*n*^(*t*); *t* ≥ 0} to [*n*] for a fixed *n* ∈ ℕ takes values in ℰ_*n*_, the set of partitions of [*n*]. The only transitions are the merging of blocks of the current partition, and the time between transitions is a random exponential with rate given by the generator of the associated semigroup. Each block in a partition represents an ancestor to the elements (the leaves) in each of the block, in the sense that distinct leaves (corresponding to sampled gene copies) *i* and *j* are in the same block at time *t* ≥ 0 if and only if they share a common ancestor at time *t* in the past [Möhle and Sagitov, 2001]. At time zero, *ξ*^*n*^(0) ≡ {{1}, …, {*n*}}, and the time inf{*t* ≥ 0 : *ξ*^*n*^(*t*) = {[*n*]}}, where the partition {[*n*]} contains only the block {[*n*]},is the time of the most recent common ancestor of the *n* sampled gene copies. By {*ξ*_*n,N*_} ≡ {*ξ*^*n,N*^ (*r*); *r* ∈ ℕ_0_} we denote the (pre-limiting) *ancestral process*. The ancestral process for a sample of *n* gene copies is a Markov sequence (a Markov process with a countable state space and evolving in discrete-time) taking values in ℰ_*n*_, where each block represents the ancestor of the leaves (gene copies; arbitrarily labelled) it contains, starting from {{1}, …, {*n*}}, and the only transitions are the merging of blocks of the current partition (we exclude further elements such as recombination and focus on gene genealogies of a single contiguous non-recombining segment of a chromosome in a haploid panmictic population). We will also use the term ‘coalescent’ for the block-counting process associated with a given coalescent. A central quantity in proving convergence of {*ξ*^*n,N*^ (⌊*t/c*_*N*_ ⌋); *t* ≥ 0 is the coalescence probability [Sagitov, 1999].

### Definition 2.2

(The coalescence probability). *Define c*_*N*_ *as the probability that two given gene copies of the same generation derive from the same parent gene copy*.

One obtains a continuous-time coalescent whenever *c*_*N*_ → 0 as *N* → ∞ with time measured in units of ⌊1*/c*_*N*_⌋ generations [Schweinsberg, 2003, Möhle and Sagitov, 2001, Sagitov, 2003]. Let *ν*_*i*_ be the random number of *surviving* offspring of the *i*th (arbitrarily labelled) individual in a haploid panmictic population. When the population is of constant size *N* it holds that *ν*_1_ + · · · + *ν*_*N*_ = *N*. Moreover, since then E [*ν*_1_] = 1, taking V(*ν*_1_) as the variance of *ν*_1_, then by Definition 2.2

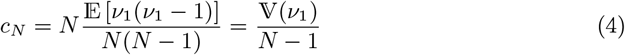

[Möhle and Sagitov, 2001, Equation 4].

For a given population model, one aims to identify the limiting process {*ξ*^*n*^(*t*); *t* ≥ 0} to which {*ξ*^*n,N*^ (⌊*t/c*_*N*_⌋), *t* ≥ 0} converges in finite dimensional distributions as *N*→ ∞. The Kingman coalescent [Kingman, 1982b,a], in which each pair of blocks in the current partition merges at rate 1, and transitions involving more than two blocks are not possible, holds for a large class of population models [Möhle, 1998]. A more general class of coalescents, in which a random number of blocks merges each time, are generally referred to as multiple-merger coalescents [Pitman, 1999, Donnelly and Kurtz, 1999, Sagitov, 1999, Schweinsberg, 2000]. They arise for example from population models of sweepstakes reproduction [Huillet and Möhle, 2013, Schweinsberg, 2003, Sargsyan and Wakeley, 2008, Eldon and Wakeley, 2006, Birkner et al., 2018, Huillet and Möhle, 2011]. Lambda-coalescents (Λ-coalescents) are multiple-merger coalescents where mergers occur asynchronously [Gnedin et al., 2014a]. They are characterised by a finite measure Λ_+_ on (0, 1] [Pitman, 1999]. In a Λ-coalescent, a given group of 2 ≤ *k* ≤ *m* blocks merges at the rate [Sagitov, 1999, Equation 4]

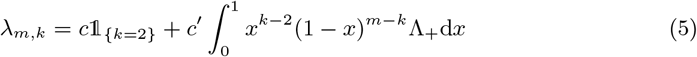

[Sagitov, 1999, Donnelly and Kurtz, 1999, Pitman, 1999]. The Kingman coalescent is thus a special case of a Λ-coalescent where *c* = 1 and Λ_+_ = 0 in (5). Birkner and Blath [2009] discuss the connection between Λ-coalescents and forward-in-time Fleming-Viot measure-valued diffusions [Fleming and Viot, 1979, Ethier and Kurtz, 1993] tracing the frequency of genetic types in populations evolving according to sweepstakes reproduction.

We will verify Case 2 of Theorem 3.5 and Case 2 of Theorem 3.6 given in § 3 by checking the conditions of [Schweinsberg, 2003, Proposition 3] (see also [Sagitov, 1999] and [Möhle and Sagitov, 2001]).

### Proposition 2.3

([Schweinsberg, 2003], Proposition 3; conditions for convergence to a Λ-coalescent). *Suppose, with c*_*N*_ *defined in Definition 2.2 (see* (4))

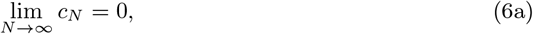

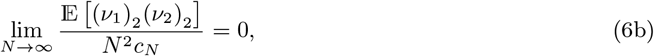

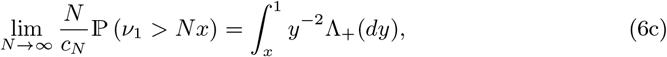

*all hold where in* (6c) Λ_+_ *is a finite measure on (the Borel subsets of)* (0, 1] *and* 0 *< x <* 1 *is fixed. Then* {*ξ*^*n,N*^ (⌊*t/c*_*N*_ ⌋); *t* ≥ 0} *converges (in the sense of convergence of finite-dimensional distributions) to* {*ξ*^*n*^(*t*); *t* ≥ 0} *with transition rates as in* (5) *with* Λ_+_ *>* 0.

On the other hand, suppose [Schweinsberg, 2003, Proposition 2], [Möhle, 2000, Section 4, Equation 14]

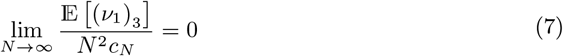

Then *c*_*N*_ → 0, and {*ξ*^*n,N*^ (⌊*t/c*_*N*_⌋); *t* ≥ 0} converges in the sense of convergence of finite-dimensional distributions (in our case equivalent to weak convergence in the *J*_1_-Skorokhod topology) to the Kingman coalescent with transition rates (5) where *c* = 1 and Λ_+_ = 0 [Möhle, 2000, 1998].

One particular family of Λ-coalescents has been much investigated (e.g. [Birkner et al., 2005, Dahmer et al., 2014, Berestycki et al., 2008, 2007, Gnedin et al., 2014b, Birkner et al., 2024]).

### Definition 2.4

(The Beta(*γ*, 2 − *α, α*)-coalescent). *The Beta*(*γ*, 2 − *α, α*)*-coalescent is a* Λ*-coalescent with transition rates as in* (5) *with c* = 0, *and* Λ_+_, *for* 0 *< α <* 2 *and* 0 *< γ* ≤ 1, *is*

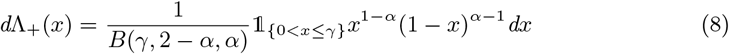

*In* (8) 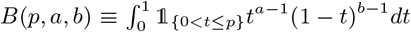 *for some given a, b >* 0 *and* 0 *< p* ≤ 1 *all fixed is the (lower incomplete when p <* 1*) beta function*.

In Definition 2.4 we take 0 *< α <* 2. The family of Beta(*γ*, 2 − *α, α*)-coalescents with *γ* = 1 and 1 ≤ *α <* 2 can be shown to follow from a particular model of sweepstakes reproduction [Schweinsberg, 2003, Equation 11]. We will study extensions of [Chetwynd-Diggle and Eldon, 2026, Equation 3.2], in turn an extension of [Schweinsberg, 2003, Equation 11], such that by combining the results we can take 0 *< α <* 2. Moreover, we will consider a *incomplete* Beta(*γ*, 2 − *α, α*)-coalescent (recall Definition 2.4) where 0 *< γ <* 1.

Xi-coalescents (Ξ-coalescents) [Schweinsberg, 2000, Sagitov, 2003, Möhle and Sagitov, 2001] extend Λ-coalescents to simultaneous mergers where ancestral lineages may merge in two or more groups simultaneously. Xi-coalescents arise, for example, from models of diploid populations evolving according to sweepstakes [Birkner et al., 2018, 2013a, Möhle and Sagitov, 2003] (see also [Sargsyan and Wakeley, 2008]), recurrent strong bottlenecks [Birkner et al., 2009], and strong positive selection resulting in recurrent selective sweeps [Durrett and Schweinsberg, 2005, Schweinsberg and Durrett, 2005]. Write

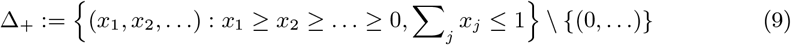

Let Ξ_+_ denote a finite measure on Δ_+_. Then, with *n* ≥ 2 blocks in a given partition, *k*_1_, …, *k*_*r*_ ≥ 2, *r* ∈ ℕ, and 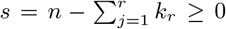, there exists a coalescent {*ξ*^*n*^(*t*); *t* ≥ 0} restricted to the partitions of [*n*] where the rate at which *k*_1_ + · · · + *k*_*r*_ ∈ {2, …, *n*} blocks merge in *r* groups with group *j* of size *k*_*j*_ is (*s* = *n* − *k*_1_ − · · · − *k*_*r*_)

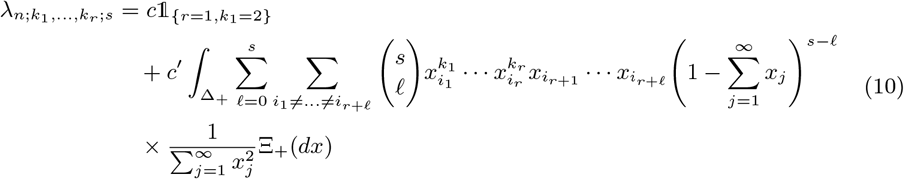

[Schweinsberg, 2000, Möhle and Sagitov, 2001]. Birkner et al [Birkner et al., 2018, Theorem A.5] summarise equivalent conditions for convergence, in the sense of finite-dimensional distributions, to a Ξ-coalescent. One condition is the existence of the limits (*k*_1_, …, *k*_*r*_ ∈ ℕ and 2 ≤ min {*k*_1_, …, *k*_*r*_} for all *r* ∈ ℕ)

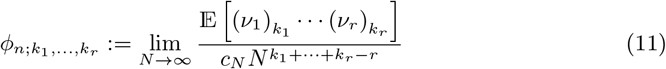

[Möhle and Sagitov, 2001].

### Proposition 2.5

([Schweinsberg, 2003] Proposition 1; convergence to a Ξ-coalescent). *Let r* ∈ ℕ *and k*_1_, …, *k*_*r*_ ≥ 2. *Suppose that the limits in* (11) *exist. Then* {*ξ*^*n,N*^ (⌊*t/c*_*N*_⌋)} *converge (in the sense of convergence of finite-dimensional distributions) to* {*ξ*^*n*^} *with transition rates as in* (10).

A discrete-time (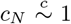 as *N* → ∞) Ξ-coalescent associated with the Poisson-Dirichlet distribution with parameter (*α*, 0) for 0 *< α <* 1 is obtained from the same model [Schweinsberg, 2003, Equation 11] of sweepstakes reproduction as gives rise to the Beta(2 − *α, α*)-coalescent [Schweinsberg, 2003, Theorem 4d]. The Poisson-Dirichlet distribution [Kingman, 1975] has found wide applicability, including in population genetics (cf. e.g. [Feng, 2010, Bertoin, 2006]). We will focus on the two-parameter Poisson-Dirichlet(*α, θ*) distribution, for 0 *< α <* 1 and *θ >* −*α* restricted to *θ* = 0 [Schweinsberg, 2003]. One can sample from the Poisson-Dirichlet(*α, θ*) distribution in the following way. Let (*W*_*k*_)_*k*_∈_ ℕ_ be a sequence of independent random variables where *W*_*k*_ has the beta distribution with parameters 1 – *α* and *θ* + *kα*. Define

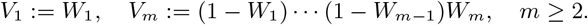

Then 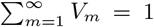 almost surely by construction, i.e. the length *V*_*m*_ of every random fragment of a stick of length 1 is added back to the sum. The distribution of (*V*_1_, *V*_2_, …) goes by the name of the two-paremeter GEM distribution, and will be denoted GEM(*α, θ*) (cf. [Feng, 2010, Ewens, 1979, Engen, 1978, McCloskey, 1965]). One refers to the law of (*V*_(1)_, *V*_(2)_, …), where *V*_(1)_ ≥ *V*_(2)_ ≥ … is (*V*_*j*_)_*j*_∈_ℕ_ ordered in descending order, as the two-parameter Poisson-Dirichlet distribution [Bertoin, 2006]. An alternative construction of PD(*α, θ*) for the case *θ* = 0 and 0 *< α <* 1 uses the ranked points of a Poisson point process on (0, ∞) with characteristic measure of the form *ν*_*α*_((*x*, ∞)) = 1_{*x>*0}_*Cx*^−*α*^ [Schweinsberg, 2003, Section 1.3].

### Definition 2.6

([Schweinsberg, 2003]; The Poisson-Dirichlet(*α*, 0)-coalescent). *Let* 0 *< α <* 1 *be fixed, write x* := (*x*_1_, *x*_2_, …) *for x* ∈ Δ_+_ *(recall* (9)*), and* 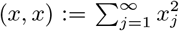. *Let F*_*α*_ *be a probability measure on* Δ_+_ *associated with the Poisson-Dirichlet*(*α*, 0)*-distribution, and* Ξ_*α*_ *a measure on* Δ_+_ *given by*

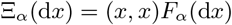

*A Poisson-Dirichlet*(*α*, 0)*-coalescent is a discrete-time* Ξ*-coalescent with* Ξ*-measure* Ξ_*α*_ *and no atom at zero. The transition probability of merging blocks in r groups of size k*_1_, …, *k*_*r*_ ≥ 2 *with current number of blocks b* ≥ *k*_1_ + · · · + *k*_*r*_ *and s* = *b* − *k*_1_ −· · · − *k*_*r*_ *is (recall* (1) *in Definition *2*.1)*,

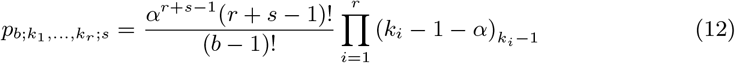

### Remark 2.7

(The transition probability of the Poisson-Dirichlet coalescent). *According to [Schweinsberg, 2003] the transition probability of the Poisson-Dirichlet*(*α*, 0)*-coalescent for an r-merger of k*_*i*_ ≥ 2 *blocks in merger i with k*_1_ + · · · + *k*_*r*_ ≤ *b and s* = *b* − *k*_1_ − · · · − *k*_*r*_, *where b is the current number of blocks, is*

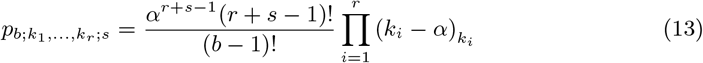

*[Schweinsberg, 2003, Equation 13]. For r* = 1 *and k*_1_ = *b* (13) *becomes*

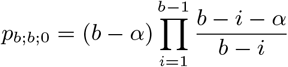

*In § 5.1 we verify* (12).

The Poisson-Dirichlet(*α*, 0)-coalescent, and the Beta(1, 2 − *α, α*)-coalescent with 1 ≤ *α <* 2 (also referred to as the Beta(2 − *α, α*)-coalescent) result from the following model of the evolution of a haploid population when the distribution of the number of potential offspring is as in (14) [Schweinsberg, 2003].

### Definition 2.8

(Evolution of the population). *Consider a haploid, panmictic population evolving in discrete (non-overlapping) generations. In any given generation each of the current individuals independently produces a random number of potential offspring according to some given law. If the total number of potential offspring produced in this way is at least some given number M, then M of them sampled uniformly at random without replacement survive to maturity and replace the current individuals; the remaining potential offspring perish. Otherwise we will assume an unchanged population over the generation (all the potential offspring perish before reaching maturity)*.

The details of what happens when the total number of potential offspring is less than the desired number can be shown to be irrelevant in the limit by tuning the model such that the mean number of potential offspring produced by any given individual is greater than 1 [Schweinsberg, 2003, Lemma 5].

Suppose a population evolves according to Definition 2.8. Let *X* denote the random number of potential offspring produced by an arbitrary individual, *α, C >* 0 fixed, and

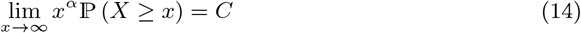

[Schweinsberg, 2003, Equation 11]. Then {*ξ*^*n,N*^ (⌊*t/c*_*N*_⌋); *t* ≥ 0} converges (in the sense of convergence of finite-dimensional distributions) to the Kingman coalescent when *α* ≥ 2, to the Beta(*γ*, 2 − *α, α*)-coalescent as in Definition 2.4 with *γ* = 1 when 1 ≤ *α <* 2, and to the Poisson-Dirichlet(*α*, 0)-coalescent as in Definition 2.6 when 0 *< α <* 1 but with time measured in generations 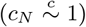 [Schweinsberg, 2003].

### Theorem 2.9

([Schweinsberg, 2003]; Theorem 4, Lemmas 6, 13, and 16). *Recall Definition 2.1, and c*_*N*_ *from Definition 2.2. Suppose a haploid population of size N evolves according to Definition 2.8 with law on the random number of potential offspring as given in* (14).

*Then* {*ξ*^*n,N*^ (⌊*t/c*_*N*_ ⌋); *t* ≥ 0} *convergences in the sense of convergence of finite-dimensional distributions to* {*ξ*^*n*^} ≡ {*ξ*^*n*^(*t*); *t* ≥ 0} *as given in each case:*

1. *1. if α* ≥ 2 *then* {*ξ*^*n*^} *is the Kingman coalescent;*
2. *if* 1 ≤ *α <* 2 *then* {*ξ*^*n*^} *is the Beta*(*γ*, 2 − *α, α*)*-coalescent as in Definition 2.4 with γ* = 1
3. *if* 0 *< α <* 1 *then* {*ξ*^*n*^} *is the Poisson-Dirichlet*(*α*, 0)*-coalescent as in Definition 2.6. By Theorem 4d, and Lemmas 13 and 16 in [Schweinsberg, 2003], as N* → ∞,

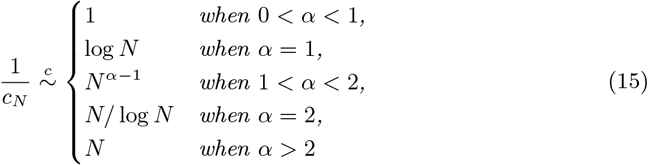

*Let E* = {*given groups of k*_1_, …, *k*_*r*_ ≥ 2 *blocks merge*}. *By Lemma 6 in [Schweinsberg, 2003], where X*_1_, …, *X*_*N*_ *denotes the i.i.d. random number of potential offspring produced by the current N individuals according to the law in* (14), *with S*_*N*_ = *X*_1_ + · · · + *X*_*N*_ *and k*_1_, …, *k*_*r*_ ≥ 2,

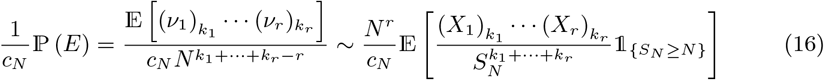

*Moreover, [Schweinsberg, 2003, Lemma 6; Equation 17]*

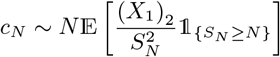

The law (14) requires that *X* can be arbitrarily large. We will consider extensions of (14) that relax this requirement. To state our results we require the following example of a Λ-coalescent with rate as in (5) with *c, c*^′^ *>* 0.

### Definition 2.10

(The *δ*_0_-Beta(*γ*, 2 − *α, α*)-coalescent). *The δ*_0_*-Beta*(*γ*, 2 − *α, α*)*-coalescent is a* Λ*-coalescent with* Λ*-measure* Λ = *cδ*_0_ + *c*^′^Λ_+_ *taking values in the partitions of* [*n*] *with rate as in* (5) *with c, c*^′^ *>* 0 *and measure* Λ_+_ *as in* (8) *in Definition 2.4*.

We also require the following extension of the Poisson-Dirichlet(*α*, 0)-coalescent (recall Definition 2.6).

### Definition 2.11

(The *δ*_0_-Poisson-Dirichlet(*α*, 0)-coalescent). *The δ*_0_*-Poisson-Dirichlet*(*α*, 0)*-coalescent is a continuous-time* Ξ*-coalescent with* Ξ*-measure* Ξ = *δ*_0_ + Ξ_+_ *taking values in the partitions of* [*n*], *where* Ξ_+_ = Ξ_*α*_ *is as in Definition 2.6. Let n* ≥ 2 *denote the current number of blocks in a partition, k*_1_, …, *k*_*r*_ ≥ 2 *with* 2 ≤ *k*_1_ + · · ·+ *k*_*r*_ ≤ *n for some r* ∈ ℕ *denoting the merger sizes of r (simultaneous) mergers, and s* = *n* − *k*_1_ − · · · −*k*_*r*_. *Then the rate at which such mergers occur is*

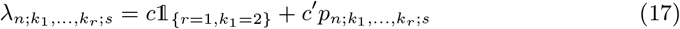

*where* 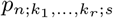 *is as in* (12).

## 3 Results

In this section we collect the results. In § 3.1 we state the main mathematical results in Theorems 3.5, 3.6, 3.8, and 3.9. In § 3.2 we give numerical examples comparing functionals (mean relative branch lengths) of the {*ξ*^*n*^} and {*ξ*^*n,N*^}, and comparing functionals of {*ξ*^*n,N*^} to the ones obtained by conditioning on the population ancestry (see § 3.2).

First we state the population model underlying the mathematical results. We will adapt the formulation of [Chetwynd-Diggle and Eldon, 2026]. Suppose a population is and evolves as in Definition 2.8. Let *X* be the random number of potential offspring produced by an arbitrary individual. Let *ζ* : ℕ → ℕ be a deterministic function and *a >* 0 be fixed. For all *k* ∈ [*ζ*(*N*)] = {1, 2, …, *ζ*(*N*)} let the probability mass function of the law of *X* be bounded by

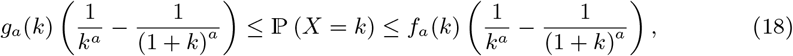

where *f*_*a*_ and *g*_*a*_ are positive functions on ℕ. The quantity *ζ*(*N*) is an upper bound on the random number of potential offspring produced by any one individual such that ℙ (*X* ≤ *ζ*(*N*)) = 1. We assign any mass outside [*ζ*(*N*)] to {*X* = 0}. The model in (18) is an extension of the one in (14) since, if *X* has law (14) then 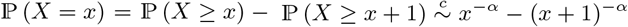 for *x* large. We will identify conditions on *g* and *f* required for convergence of the (time-rescaled) ancestral process to a non-trivial limit. Write

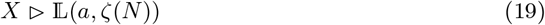

when the law of *X* is according to (18) with *a* and *ζ*(*N*) as given each time.

We define

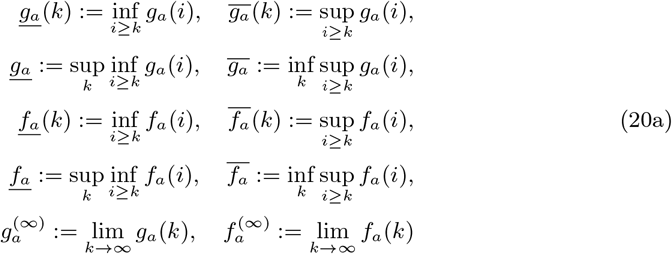

We assume 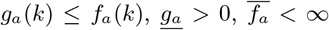, and that 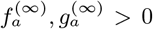 exist. Then, for all *k* ∈ [*ζ*(*N*)],

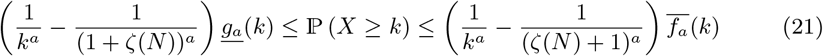

### Assumption 3.1

(E [*X*] *>* 1). *Suppose X* ▷ L(*a, ζ*(*N*)) *(recall* (19)*). The functions f*_*a*_ *and g*_*a*_ *in* (18) *are such that* E [*X*] *>* 1 *for all N* ∈ ℕ.

Assumption 3.1 results in ℙ (*X*_1_ + · · · + *X*_*N*_ *< N*) decreasing exponentially in *N*, where *X*_1_, …, *X*_*N*_ are independent copies of *X* [Schweinsberg, 2003, Lemma 5].

For *X* ▷ L(*a, ζ*(*N*)), using the lower bound in (50) and (51) in Lemma 5.3 with *a*^′^ ≡ *a*1_{0*<a<*1}_ + 1_{*a*≥1}_

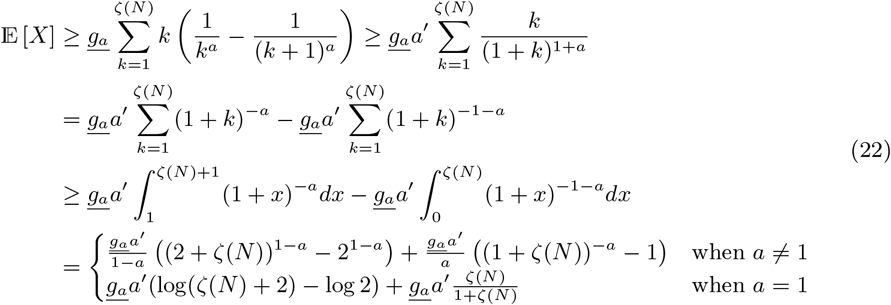

For ease of reference we state notation used from here on.

### Definition 3.2

(Notation). *Let X*_1_, …, *X*_*N*_ *denote the independent random number of potential offspring (recall Definition *2*.8) produced in an arbitrary generation by the current N individuals. Write*

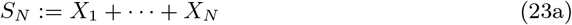

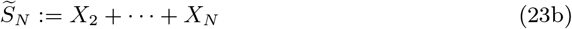

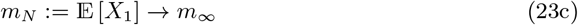

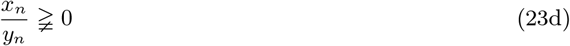

*where m*_∞_ *in* (23c) *is the limit as N* → ∞, *and* (23d) *for positive sequences* (*x*_*n*_) *and* (*y*_*n*_) *will mean that either x*_*n*_*/y*_*n*_ *converges to some ℓ >* 0, *or x*_*n*_*/y*_*n*_ *diverges as n* → ∞.

### 3.1 Beta- and Poisson-Dirichlet-coalescents

Before we state the results we define random environments (Definitions 3.3 and 3.4); the population then evolves in the given environment as in Definition 2.8. The environments are modelled as simple mixture distributions on the law of the number of potential offspring (recall (18)). Depending on the specific scenario each time, we obtain a continuous-time Λ- or Ξ-coalescent with an atom at zero, and with time measured in units proportional to either *N/* log *N* or *N* generations.

#### Definition 3.3

(Type *A* random environment). *Suppose a population evolves according to Definition 2.8. Fix* 0 *< α <* 2 *and* 2 ≤ *κ. Recall* (19) *and the X*_1_, …, *X*_*N*_ *from Definition 3.2. Write E for the event when X*_*i*_ ▷ L(*α, ζ*(*N*)) *for all i* ∈ [*N*], *and E*^c^ *for the event when κ replaces α in E (X*_*i*_ ▷ L(*κ, ζ*(*N*)) *for all i* ∈ [*N*]*); ζ*(*N*) *is fixed for each N* ∈ ℕ. *Let* (*ε*_*N*_)_*N* ∈ℕ_ *be a positive sequence with* 0 *< ε*_*N*_ *<* 1 *for all N. It may hold that ε*_*N*_ → 0 *as N* → ∞. *Suppose*

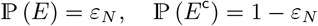

Definition 3.3 says that the population evolves in an environment where changes to the environment affect all individuals equally so that at any given time the *X*_1_, …, *X*_*N*_ are i.i.d. The bound *ζ*(*N*) stays the same between *E* and *E*^c^. We will identify conditions on *ε*_*N*_ (see (53) in Lemma 5.7) so that the ancestral process {*ξ*^*n,N*^ (⌊*t/c*_*N*_⌋); *t* ≥ 0} converges (in the sense of convergence of finite-dimensional distributions) to a non-trivial limit as *N*→ ∞. Moreover, the *X*_1_, …, *X*_*N*_ in Definition 3.3 are i.i.d. copies of *X* where (recall (3) in Definition 2.1)

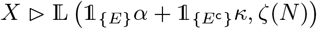

We will also consider a scenario where the *X*_1_, …, *X*_*N*_ stay independent but may not always be identically distributed.

#### Definition 3.4

(Type *B* random environment). *Suppose a population evolves according to Definition 2.8. Fix* 0 *< α* ≤ 1 *and κ* ≥ 2. *Recall* (19) *and write E*_1_ *for the event (*[*N*] = {1, 2, …, *N*} *by Definition *2*.1) when there exists exactly one i* ∈ [*N*] *such that X*_*i*_ ▷ L(*α, ζ*(*N*)), *and X*_*j*_ ▷ L(*κ, ζ*(*N*)) *for all j* ∈ [*N*] \ {*i*}. *When E*_1_ *occurs, the ‘lucky’ individual (the index i) is picked uniformly at random. When event* 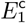 *is in force κ replaces α in E*_1_ *(X*_*i*_ ▷ L(*κ, ζ*(*N*)) *for all i* ∈ [*N*]*) from Definition 3.3. Let* 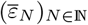 *be a sequence with* 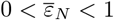 *for all N* ∈ ℕ. *It may hold that* 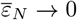 *as N* → ∞. *Suppose* 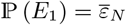, *and* 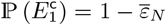.

Definition 3.4 says that when the environment turns favorable for producing potential offspring through *α* (event *E*_1_ occurs), exactly one individual sampled uniformly at random will do so, i.e. with probability 1*/N* it holds that

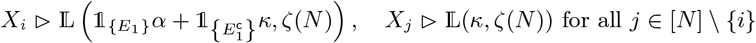

for all *i* ∈ [*N*]. Since the ‘lucky’ individual is sampled at random when event *E*_1_ occurs the *X*_1_, …, *X*_*N*_ are exchangeable and we can use the results of [Möhle and Sagitov, 2001] for proving convergence of {*ξ*^*n,N*^}. As in Definition 3.3 the bound *ζ*(*N*) stays the same between *E*_1_ and 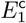. We will identify conditions on 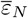 (see (68) in Lemma 5.16) such that{*ξ*_*n,N*_ (⌊*t/c*_*N*_ ⌋); *t* ≥ 0} converges to a nontrivial limit. In Theorems 3.5 and 3.6 the law for the number of potential offspring is as in (18). Write (for *κ* ≥ 2)

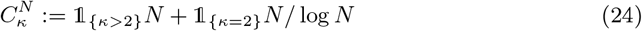

Section 5.2 contains a proof of Theorem 3.5.

#### Theorem 3.5

(Evolution in a type *A* random environment (Definition 3.3)). *Suppose a population evolves as defined in Definitions 2.8 and 3.3 (type A random environment) with the law for the X*_1_, …, *X*_*N*_ *given by* (18), *that Assumption 3.1 holds*, 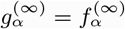, *and* 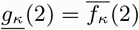 *(recall* (20a)*). Then* {*ξ*^*n,N*^ (⌊*t/c*_*N*_⌋); *t* ≥ 0} *converges in the sense of convergence of finite-dimensional distributions to* {*ξ*^*n*^} ≡ {*ξ*^*n*^(*t*); *t* ≥ 0} *where* {*ξ*^*n*^} *is as specified in each case*.

1. *Suppose ζ*(*N*)*/N* → 0 *and* 1 ≤ *α <* 2. *Then* {*ξ*^*n*^} *is the Kingman coalescent*.
2. *Suppose* 1 ≤ *α <* 2 *and ζ*(*N*)*/N* ≩ 0 *(recall* (23d) *in Definition *3*.2). Take* (*ε*_*N*_)_*N*_ *from Definition 3.3 with ε*_*N*_ = *cN* ^*α*−2^1_{*κ>*2}_ + 1_{*κ*=2}_ log *N for some fixed c >* 0. *Then* {*ξ*^*n*^} *is the δ*_0_*-Beta*(*γ*, 2 − *α, α*)*-coalescent defined in Definition 2.10 with* 1 *< m*_∞_ *<* ∞ *(recall* (23c)*). The transition rate for a k-merger of a given group of k* ∈ {2, 3, …, *n*} *blocks is*

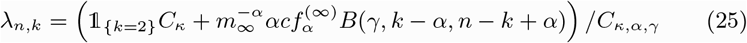

*with* 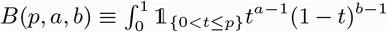 *(a, b >* 0, 0 *< p* ≤ 1*) and (recalling* 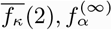 *from* (20a))

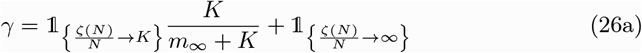

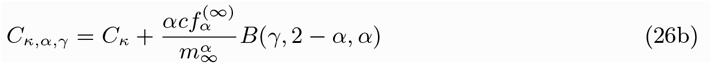

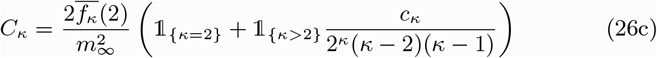

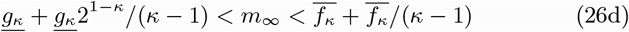

*where in* (26c) *we have κ* + 2 *< c*_*κ*_ *< κ*^2^ *for κ >* 2.

*3. Suppose* 0 *< α <* 1 *and ζ*(*N*)*/N* ^1*/α*^ → ∞ *and* 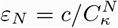 *with c >* 0 *fixed. Then* {*ξ*^*n*^} *is the δ*_0_*-Poisson-Dirichlet*(*α*, 0) *coalescent defined in Definition 2.11 with transition rates (recall* (17))

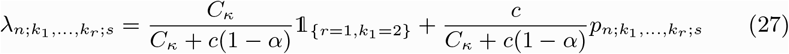

*with C*_*κ*_ *as in* (26c) *and p*_2;2;0_ = 1 − *α so that λ*_2;2;0_ = 1.

*In all cases (recall* (2) *in Definition 2.1 and c*_*N*_ *from Definition 2.2 and* 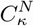 *from* (24)*) as N* → ∞

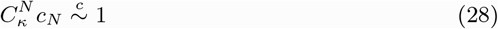

In Theorem 3.5 the population evolves according to Definition 3.3, where everyone produces potential offspring through *α* (recall *α < κ*) when the environment turns favorable. In Theorem 3.6 the population evolves according to Definition 3.4 where exactly one randomly picked individual produces potential offspring through *α* when the conditions are favorable; thus the *X*_1_, …, *X*_*N*_ may not always be identically distributed. Suppose 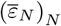 from Definition 3.4 takes the form as in (68) in Lemma 5.16,

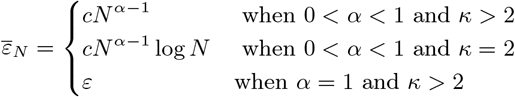

with *c >* 0 and 0 *< ε <* 1 both fixed.

Section 5.3 contains a proof of Theorem 3.6.

#### Theorem 3.6

(Evolution in a type *B* random environment (Definition 3.4)). *Suppose a population evolves as defined in Definitions 2.8 and 3.4 (type B random environment), Assumption 3.1 holds, and* 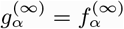 *(recall* (20a)*). Take* 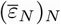 *from Definition 3.4 as in* (68) *in Lemma 5.16. Then* {*ξ*^*n,N*^ (⌊*t/c*_*N*_⌋); *t* ≥ 0} *converges in the sense of convergence of finite-dimensional distributions to* {*ξ*^*n*^} ≡ {*ξ*^*n*^(*t*); *t* ≥ 0} *where* {*ξ*^*n*^} *is as specified in each case*.

1. *Suppose ζ*(*N*)*/N* → 0. *Then* {*ξ*^*n*^} *is the Kingman coalescent*.
2. *Suppose* 0 *< α* ≤ 1 *and ζ*(*N*)*/N* ≩ 0 *(recall* (23d) *in Definition *3*.2). Then* {*ξ*^*n*^} *is the δ*_0_*-Beta*(*γ*, 2 − *α, α*)*-coalescent defined in Definition 2.10 with* 1 *< m*_∞_ *<* ∞ *(recall* (23c)*). The transition rates are as in* (25) *with c*_*α*_ *replacing c where c*_*α*_ = 1_{*α*=1}_*ε* + 1_{0*<α<*1}_*c where* 0 *< ε <* 1 *and c >* 0 *as in* (68).

*Furthermore*, (28) *holds in both cases*.

#### Remark 3.7

(The behaviour of *ζ*(*N*)). *When ζ*(*N*)*/N* → ∞ *in Theorems 3.5 and 3.6 (or ζ*(*N*)*/N* ^1*/α*^ → ∞ *as in Case 3 of Theorem *3*.5) it is with the understanding that* 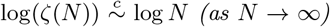, *so that ζ*(*N*) *is at most of the form N* ^1+*η*^ *for* 0 *< η* ≪ 1 *(see Lemma *5*.14). One could also think of* 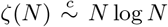 *(or N* ^1*/α*^ log *N in Case 3 of Theorem *3*.5). We do not see ζ*(*N*) *increasing exponentially (or faster) with N*.

#### 3.1.1 A general random environment

This section is a response to the question

> *‘If I take 2 (or m, integer) different reproduction models (Cannings models) that have known coalescent limits and mix them from generation to generation, do they have a coalescent limit (with which time scaling), what is the limit and is it e.g. a Lambda coalescent with Lambda a sum of the Lambdas of the underlying different reproduction models (assuming their limits are Lambda-coalescents)?’*

We have considered random environments each split into two possibilities. When warranted by the biology of the study system at hand, one may want to consider a random environment split into several (say *m* ≥ 2) events, where each event entails reproduction according to a given Cannings model with known coalescent limit. For example, one may extend the type *A* random environment (Definition 3.3) to one where, with 0 *< α <* 1 and 1 ≤ *β <* 2 both fixed,

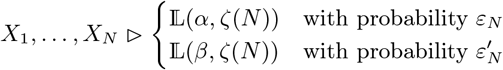

such that 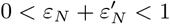 for all *N* ∈ ℕ, and with *κ* ≥ 2 fixed,

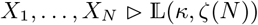

with probability 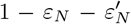. Choosing 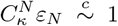, and 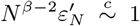 (recall (53) in Lemma 5.7), our calculations show we would obtain a continuous-time Ξ-coalescent with an atom at zero and with the transition rate being a linear combination of the rates given in (25) and (27) in Theorem 3.5 with timescale proportional to 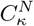 (24). In contrast, the continuous-time coalescent considered in Appendix A runs on a timescale proportional to *N* ^*β*−1^ generations (1 *< β <* 2) and so runs into the problem of recovering mutations observed in a given sample of DNA gene copies, especially for estimates of *β* close to 1. Barring any additional complications such as separation of timescales due to e.g. selfing [Möhle, 1998] or diploidy [Möhle and Sagitov, 2003, Birkner et al., 2013a, 2018], a central condition required to hold to prove convergence to a continuous-time Ξ-coalescent is the existence of the limits in (11) [Möhle and Sagitov, 2001, Equation 16] (see [Birkner et al., 2018, Equation 6] for the diploid version). Consider a random environment consisting of *m* events *E*^(*i*)^, where event *E*^(*i*)^ occurs with probability 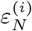 such that when *E*^(*i*)^ occurs the (haploid panmictic) population reproduces according to some given Cannings reproduction scheme with timescale 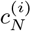 such that 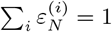 and for at least one index *i* it holds 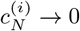. Moreover, suppose each Cannings model has a known coalescent limit with transition rates 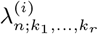 (10) driven by a measure Ξ^(*i*)^ determined by the Cannings model specified in event *E*^(*i*)^. Then

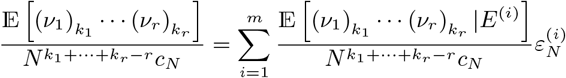

and Condition (11) becomes equivalent to checking that

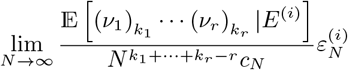

exists for all *i* ∈ [*m*] and *r* ∈ ℕ and *k*_1_, …, *k*_*r*_ ≥ 2 (and with the 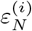 suitably tuned; recall that *ν*_1_, …, *ν*_*N*_ are the random numbers of offspring of the current *N* individuals in a haploid panmictic population of constant size *N*). Convergence to a continuous-time Ξ-coalescent with transition rates as a linear combination of the individual rates 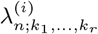 then follows by [Möhle and Sagitov, 2001, Theorem 2.1]. However, even though extending the simple random environments behind Theorems 3.5 and 3.6 to more general settings is straightforward, the biology of the study system at hand (and a general principle of parsimony to avoid overfitting) should dictate the modeling.

### 3.2 Comparing processes

In this section we use simulations to compare functionals of coalescents {*ξ*^*n*^} and ancestral processes {*ξ*_*n,N*_} in the domain of attraction of {*ξ*^*n*^} as given in Theorems 3.5 and 3.6. We are interested in comparing {*ξ*^*n*^} and {*ξ*^*n,N*^}.

Let |*A*| be the number of elements in a given finite set *A*, write *τ* ^*N*^ (*n*) := inf *j* ∈ ℕ_0_ : |*ξ*^*n,N*^ (*j*)| = 1, and *τ* (*n*) := inf {*t* ≥ 0 : |*ξ*^*n*^(*t*)| = 1}. For any given ancestral process{*ξ*^*n,N*^} and coalescent {*ξ*^*n*^} consider the functionals

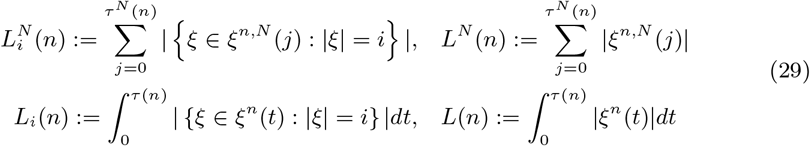

[Birkner et al., 2018]. Viewing {*ξ*^*n,N*^} and {*ξ*^*n*^} as gene genealogies (trees) the random variables 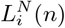 resp. *L*_*i*_(*n*) can be interpreted as the random length of branches supporting *i* ∈ [*n* −1] leaves when the gene genealogy with *n* leaves is governed by a given ancestral process {*ξ*_*n,N*_} resp. coalescent {*ξ*^*n*^}.

Given the example gene genealogy for *n* = 10 in Figure 1, *L*_1_(*n*) (resp. 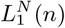) would be the total length of the blue branches, *L*_2_(*n*) the green branches, *L*_3_(*n*) the black branch, and *L*_7_(*n*) the red branch.

**Figure 1.**
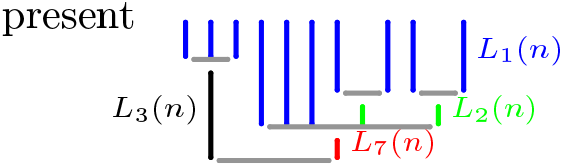
The functionals 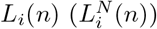 for an example gene genealogy.

It holds that 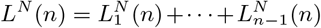, and *L*(*n*) = *L*_1_(*n*) +· · · + *L*_*n*−1_(*n*). Define, for all *i* ∈ [*n* − 1],

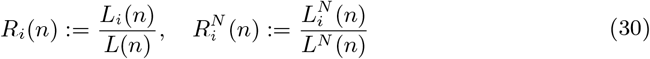

Both 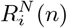 and *R*_*i*_(*n*) are well defined since *L*(*n*) *>* 0 and *L*^*N*^ (*n*) ≥ *n* both almost surely. We will compare approximations of E [*R*_*i*_(*n*)] to approximations of 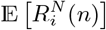 for corresponding processes. Interpreting {*ξ*^*n*^} and {*ξ*^*n,N*^} as ‘trees’ one may view *R*_*i*_(*n*) and 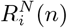 as random ‘relative branch lengths’. Under the infinitely-many sites mutation model [Kimura, 1969, Watterson, 1975] the site-frequency spectrum, or the count of mutations of given frequencies in a sample, corresponds directly to branch lengths [Eldon et al., 2015]. We will use the notation (recall (30))

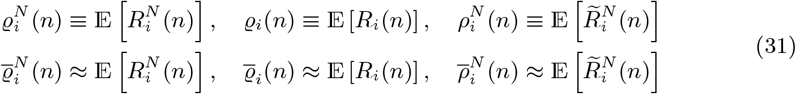

Where 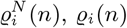, and 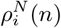 are functionals we are interested in, and 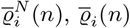, and 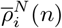 are the corresponding approximations. The quantities 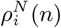 are relative branch lengths read off fixed complete sample trees (conditioning on the population ancestry and averaging over the ancestries; see § 3.2.5).

Various evolutionary histories may yield similar site-frequency spectra, thus giving rise to non-identifiability issues [Myers et al., 2008, Bhaskar and Song, 2014, Rosen et al., 2018, Freund et al., 2023]. The site-frequency spectrum may, nevertheless, be used in conjunction with other statistics in inference [Freund and Siri-Jégousse, 2021].

Whenever {*ξ*^*n,N*^} converges to {*ξ*^*n*^} it should follow that 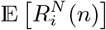 converges to E [*R*_*i*_ (*n*)] as *N* → ∞ (and with *n* fixed). Moreover, one would hope that the functionals would be in good agreement for all ‘not too small’ *N*.

In Appendix B we consider the quantity (recall *L*(*n*) *>* 0 almost surely)

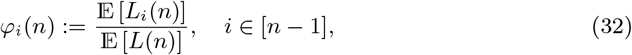

for the *δ*_0_-Beta(*γ*, 2 − *α, α*)-coalescent (see Figure B1). Values of *φ*_*i*_(*n*) can be computed exactly using a recursion derived for Λ-coalescents [Birkner et al., 2013b].

For sampling {*ξ*^*n,N*^} we will use the following example of (18). Let (*p*_*k*_(*a*))_*k*∈ℕ_ be probability weights with

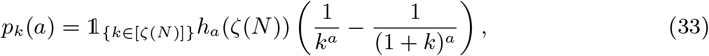

where *h*_*a*_(*ζ*(*N*)) is such that 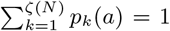 (i.e. *h*_*a*_(*ζ*(*N*)) = (1 (1 + *ζ*(*N*))^−*a*^)^−1^ and *h*_*a*_(*ζ*(*N*)) → 1 as *ζ*(*N*) → ∞). If *ζ*(*N*) *<* ∞ for *N* fixed (33) is a special case of (18) with *f*_*a*_(*k*) = *g*_*a*_(*k*) = *h*_*a*_(*ζ*(*N*)) for all *k* (and of (14) when *ζ*(*N*) = for all *N*). For the numerical experiments we will assume that the sample comes from a population evolving according to Definition 2.8, and to either Definition 3.3 or 3.4 as specified each time. Briefly, Definition 2.8 describes the evolution of a haploid panmictic population evolving in non-overlapping generations through the generation of potential offspring, and Definitions 3.3 (type *A* random environment) and 3.4 (type *B* random environment) describe the random environments used, the difference being that when Definition 3.4 is in force the numbers of potential offspring (always independent) may not always be identically distributed. The probability weights of the random number of potential offspring are as in (33) with *a* and *ζ*(*N*) as given in each case. Case 2 of Theorem 3.5 and Case 2 of Theorem 3.6 then show that choosing (*ε*_*N*_)_*N*_, 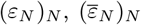 suitably the (time-rescaled) ancestral process converges to a *δ*_0_-Beta(*γ*, 2 − *α, α*)-coalescent (recall Definition 2.10) with transition rates as in (25) with 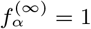. With *κ* = 2 the transition rate for a *k*-merger of *k* ∈ {2, 3, …, *n*} blocks is

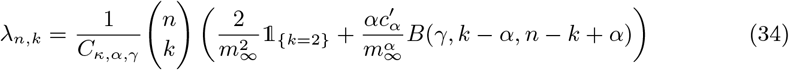

with 0 *< α <* 2, *m*_∞_ from (23c) and *γ* from (26a) and *B*(*γ, k* − *α, n* − *k*+*α*) as in Definition 2.4 and

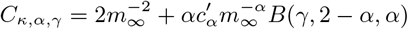

so that *λ*_2,2_ = 1. When *α* = 1 we need to decide, which of the random environments (Definitions 3.3 or 3.4) holds; let 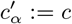 if Definition 3.3 holds, and 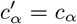 if Definition 3.4 holds where *c*_*α*_ = 1_{0*<α<*1}_*c* + 1_{*α*=1}_*ε* and 0 *< ε <* 1 is fixed.

We see, with *κ* ≥ 2,

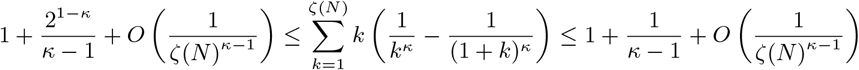

Then, when a population evolves according to Definition 2.8 and Definition 3.3 with 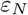 as in (53) when Definition 3.3 holds, and 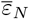 as in (68) when Definition 3.4 holds we see that we can approximate *m*_∞_ = lim_*N* →∞_ E [*X*_1_] (recall (23c)) with m where *κ* ≥ 2 and

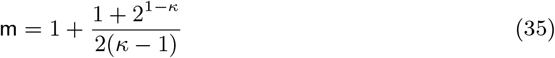

Thus, the *δ*_0_-Beta(*γ*, 2 − *α, α*)-coalescent when *κ* = 2 involves the parameters, *c*^′^_*α*_, *α, γ*, in addition to *m*_∞_ (here approximated by m as in (35)).

We also compare 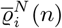 to 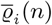 as predicted by the *δ*_0_-Poisson-Dirichlet(*α*, 0)-coalescent (recall Case 3 of Theorem 3.5), where the distribution of the number of potential offspring is as in (35) with *ζ*(*N*) = *N* ^1*/α*^ log *N* (so that *ζ*(*N*)*/N* ^1*/α*^ as *N* → ∞ as *N* → ∞ required). Recall from Case 3 of Theorem 3.5 the transition rates of the *δ*_0_-Poisson-Dirichlet(*α*, 0)-coalescent,

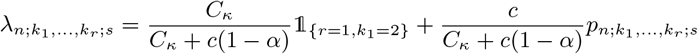

where *C*_*κ*_ is as in (26c) with *m*_∞_ approximated as in (35).

In § 3.2.1 we describe an algorithm for approximating 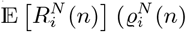, annealed mean relative branch lengths predicted by a given ancestral process, recall (29), (30), (31)), in § 3.2.2 for approximating 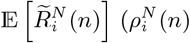, quenched mean relative branch lengths), and in § 3.2.3 for sampling from the *δ*_0_-Poisson-Dirichlet(*α*, 0)-coalescent. We used `R`[R Core Team, 2026] and the shell tool `parallel`[Tange, 2011] for generating the graphs.

#### 3.2.1 Approximating 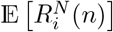

Recall § 3.2, in particular (30). In this section we describe the (non-quenched, annealed) algorithm for sampling branch lengths of a gene genealogy of a sample of size *n* from a finite haploid panmictic population of constant size *N*. Suppose the population evolves according to Definition 2.8. Our algorithm returns a realisation 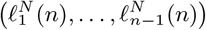 of 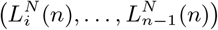. Since we are working with a Markov sequence (a Markov process evolving in discrete time), and we are only interested in the site-frequency spectrum, we only keep track of the current block sizes *b*_1_(*g*), …, *b*_*m*_(*g*), where *b*_*i*_(*g*) is the size (number of leaves the block is ancestral to) of block *i* at time *g*, so that *b*_*i*_(*g*) ∈ [*n*], and *b*_1_(*g*) + · · ·+ *b*_*m*_(*g*) = *n* when there are *m* blocks. Let 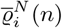 (initialised to zero) denote an estimate of 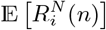 obtained in the following way:

1. 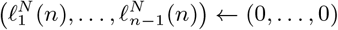
2. set the block sizes *b*_1_(0), …, *b*_*n*_(0) to 1
3. while there are at least two blocks repeat the following steps in order
  a. update the current branch lengths 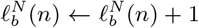 for *b* ∈ {*b*_1_(*g*), …, *b*_*m*_(*g*)} given that there are *m* blocks at time *g*
  b. sample a realisation *x*_1_, …, *x*_*N*_ of the juvenile numbers *X*_1_, …, *X*_*N*_
  c. sample number of blocks per family according to a multivariate hypergeometric with parameters *m* (the current number of blocks) and *x*_1_, …, *x*_*N*_
  d. merge blocks at random according to the numbers sampled in (c)
4. update the estimate 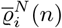 of 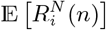:

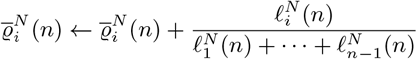
5. after repeating steps (1) to (4) a given number (*M*) of times return 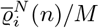

See Figures 2, 3, and Figures C1 and C2 in Appendix C for examples.

**Figure 2.**
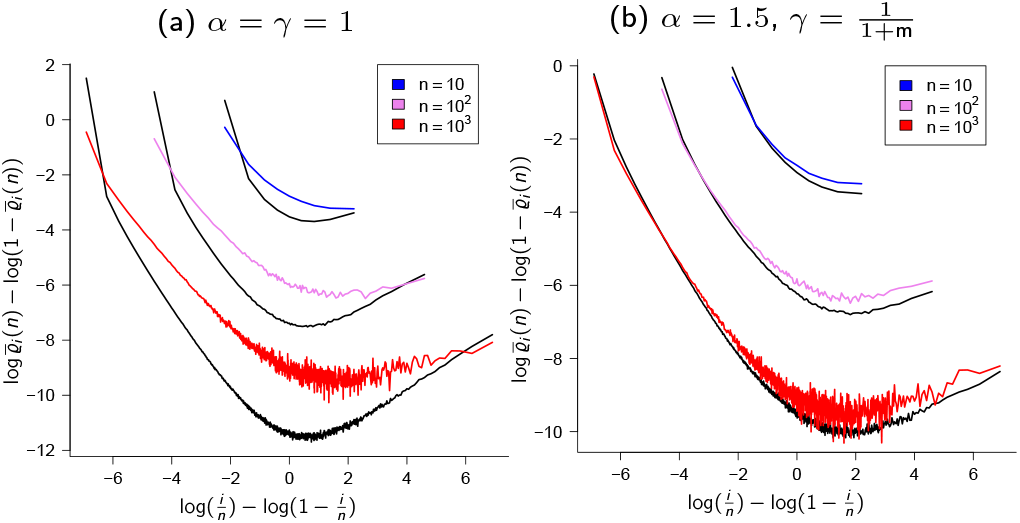
Definition 3.3 (type *A* random environment) and the *δ*_0_-Beta(*γ*, 2 − *α, α*)-coalescent. Comparing 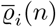 (estimates of mean relative branch lengths predicted by the *δ*_0_-Beta(*γ*, 2− *α, α*)-coalescent, black lines) and 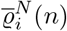 (estimates of mean relative branch lengths predicted by {*ξ*^*n,N*^} in attraction of the *δ*_0_-Beta(*γ*, 2 *α, α*)-coalescent, recall Figure 1 and (31)) when *N* = 10^3^, *κ* = 2, *c* = 1, and *γ* as shown; black lines are 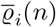 for sample size *n* as shown with rates as in (25) in Theorem 3.5, coloured lines are 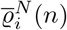 for a sample from a population of finite size *N* evolving according to Definition 2.8 and Definition 3.3 with the potential offspring distributed as in (33) and with *ε*_*N*_ = *cN* ^*α*−2^ log *N* as in (53) in Lemma 5.7; the case *γ* = 1 is compared to *ζ*(*N*) = *N* log *N*, and *γ* = 1*/*(1 + m) to *ζ*(*N*) = *N* with m as in (35) approximating *m*_∞_; 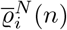 from 10^4^ experiments. Further graphs are in Figure C1 in Appendix C. The scale of the ordinate (vertical axis) may vary between graphs.

**Figure 3.**
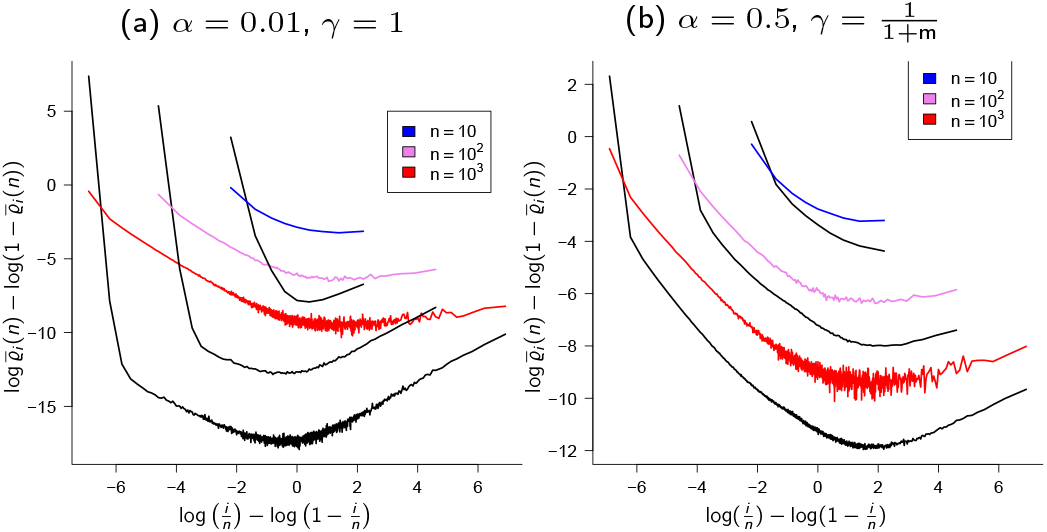
Definition 3.4 (type *B* random environment) and the *δ*_0_-Beta(*γ*, 2 − *α, α*)-coalescent. Comparing 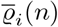 (estimates of mean relative branch lengths predicted by the *δ*_0_-Beta(*γ*, 2−*α, α*)-coalescent, black lines) and 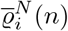 (estimates of mean relative branch lengths predicted by {*ξ*^*n,N*^} in attraction of the *δ*_0_-Beta(*γ*, 2 *α, α*)-coalescent, recall (31)) when *N* = 10^3^, *κ* = 2, *c* = 1, and with *α, γ*, and sampled size *n* as shown; black lines are 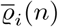 with rates as in (25), coloured lines are estimates of 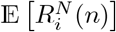 for a sample from a population evolving according to Definition 2.8 and Definition 3.4 with the potential offspring distributed as in (33) and with 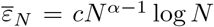 log *N* as in (68) in Lemma 5.16; the case *γ* = 1 is compared to *ζ*(*N*) = *N* log *N*, and the case *γ* = 1*/*(1 +m) compared to *ζ*(*N*) = *N* with m as in (35) approximating *m*_∞_; 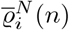 from 10^4^ experiments. Further graphs are in Figure C2 in Appendix C. The scale of the ordinate (vertical axis) may vary between graphs.

#### 3.2.2 Approximating 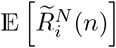

In this section we briefly describe an algorithm for approximating 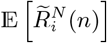 (quenched mean relative branch lengths, 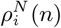, recall (31)). Figure 5 and Figure D1 in Appendix D hold examples. Recall [*n*] and ℕ from Definition 2.1. Let (*A*_*i*_(*g*))_*g*∈ℕ∪{0},*i*∈[*N*]_ denote the ancestry of the population where *A*_*i*_(*g*) ∈ [*N*] is the level of the immediate ancestor of the individual occupying level *i* at time *g*. We set *A*_*i*_(0) = *i* for *i* ∈ [*N*]. If *A*_*i*_(*g*) = *A*_*j*_ (*g*) for *i≠ j* the individuals occupying levels *i* and *j* at time *g* derive from the same immediate ancestor. If the individual on level *i* at time *g* produced *k* surviving offspring then 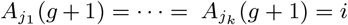. Each individual ‘points’ to its’ immediate ancestor.

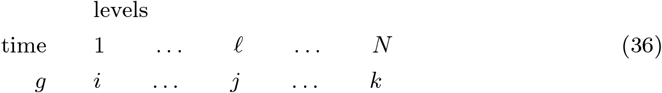

In (36), the individual occupying level 1 resp. *ℓ* resp. *N* at time *g* derives from the immediate ancestor occupying level *i* resp. *j* resp. *k* at time *g* − 1.

A ‘complete’ sample tree is one where the leaves have a common ancestor. Let

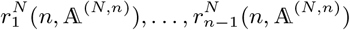

denote the realised relative branch lengths of a complete sample tree whose ancestry is given by A^(*N,n*)^, then we estimate, for *i* ∈ {1, 2, …, *n* − 1},

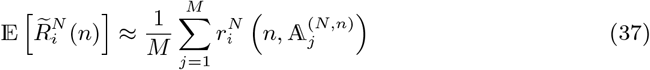

where *M* is the number of experiments, the number of realised ancestries A^(*N,n*)^.

We summarize the algorithm to estimate 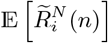.

1. For each experiment:
  a. initialise the ancestry to *A*_*i*_(0) = *i* for *i* ∈ [*N*]
  b. until a complete sample tree is found:
    i. draw a random sample, i.e. sample *n* of *N* levels at the most recent time
    ii. check if the tree of the given sample is complete, if not discard the sample and record the ancestry of a new set of surviving offspring: A. sample numbers of potential offspring *X*_1_, …, *X*_*N*_ B. given *X*_1_, …, *X*_*N*_ potential offspring sample the surviving offspring uniformly at random without replacement and update the ancestry; if the individual on level *i* at time *g* produced *k* surviving offspring then 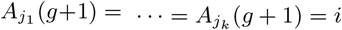.
  c. given a complete tree read the branch lengths off the tree and merge blocks according to the ancestry; suppose two blocks have ancestors on levels *i* and *j* at time *g*, if *A*_*i*_(*g*) = *A*_*j*_ (*g*) the blocks are merged;
  d. given the branch lengths of a complete tree update the estimate of 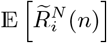
2. from *M* realised ancestries 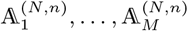 return an estimate (37) of 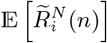

#### 3.2.3 Sampling from the *δ*_0_-Poisson-Dirichlet(*α*, 0)-coalescent

In this section we briefly describe an algorithm for samping from the *δ*_0_-Poisson-Dirichlet(*α*, 0) coalescent (recall Definition 2.11). See Figures 4, and Figures E1 and E2 in Appendix E for examples of approximations 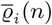 of E [*R*_*i*_(*n*)] obtained using the algorithm.

**Figure 4.**
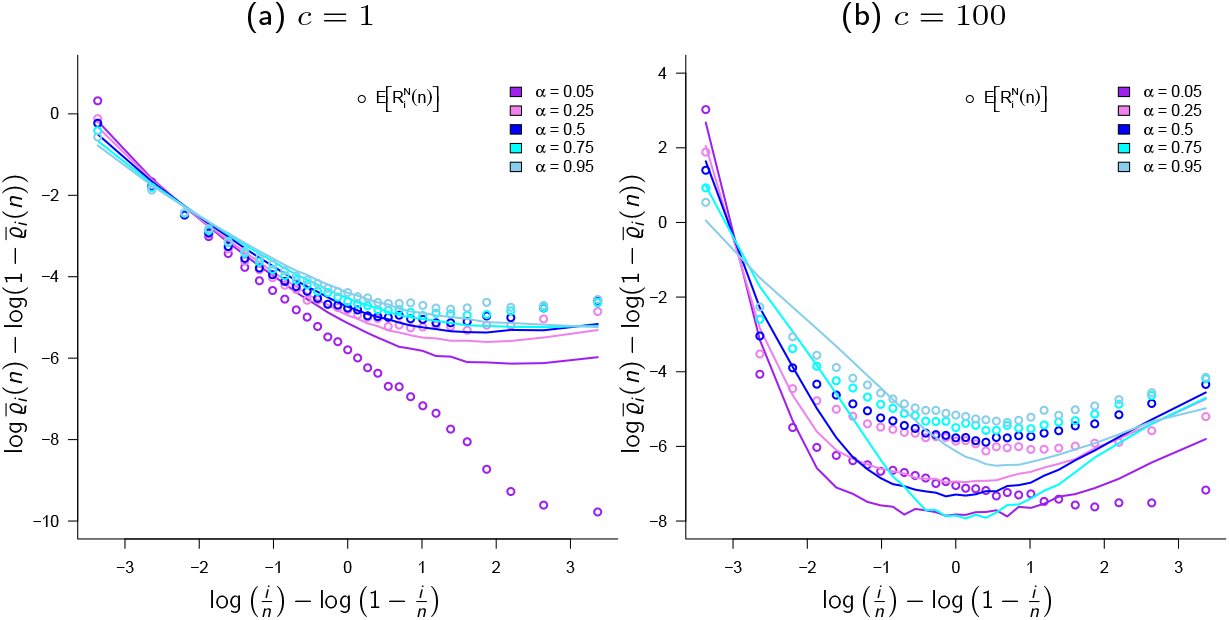
The *δ*_0_-Poisson-Dirichlet(*α*, 0)-coalescent. Approximations 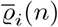 (31) of E [*R*_*i*_(*n*)] (mean relative branch lengths predicted by the *δ*_0_-Poisson-Dirichlet(*α*, 0)-coalescent, lines) predicted by the *δ*_0_-Poisson-Dirichlet(*α*, 0)-coalescent compared to 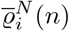 (mean relative branch lengths predicted by {*ξ*^*n,N*^}in the attraction of the *δ*_0_-Poisson-Dirichlet(*α*, 0)-coalescent, circles) when the population evolves according to Definition 3.3 (type *A* random environment) with *ζ*(*N*) = ∞, *ε* = *c*(log *N*)*/N* for *c* as shown, *N* = 3000, *α* as shown and *κ* = 2. The scale of the ordinate (y-axis) may vary between the graphs. See § 3.2.3 for an algorithm for sampling from the *δ*_0_-Poisson-Dirichlet(*α*, 0)-coalescent Further graphs are in Figure E1 in Appendix E. The scale of the ordinate (vertical axis) may vary between graphs.

By Case 3 of Theorem 3.5 the transition rate (where 2 ≤ *k*_1_, …, *k*_*r*_ ≤ *n*, ∑ _*i*_ *k*≤ *n, s* = *n* ™ ∑ _*j*_ *k*_*j*_),

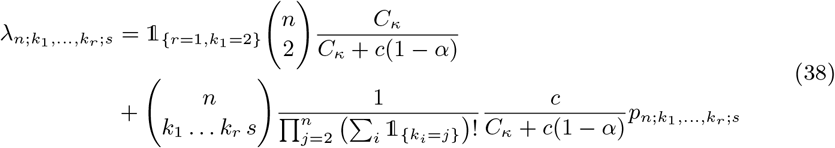

where 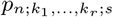 is as in (12). For each *m* ∈ {2, 3, …, *n*} and *r* ∈ [⌊*m/*2⌋] we list all possible ordered merger sizes 2 ≤ *k*_1_ ≤ · · · ≤ *k*_*r*_ ≤ *m* with ∑_*i*_ *k*_*i*_ ≤ *m*; the algorithms for listing mergers borrow from algorithms for listing partitions of integers. The time until a merger when *n* blocks is then an exponential with rate the sum of 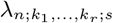 in (38) over the ordered mergers, and the actual merger size(s) can be efficiently sampled given a listing of all the mergers ordered in descending order according to the rate.

#### 3.2.4 Comparing 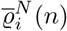 and 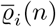

Recall 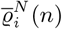 (approximations of annealed mean relative branch lengths associated with an ancestral process) and 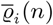 (approximations of annealed mean relative branch lengths associated with a coalescent) from (29), (30), (31). In Figures 2–4 we compare 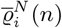 and 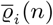 (black lines) when the population evolves as in Definition 2.8 and Definition 3.3 (type *A* random environment; Figures 2, 4) and Definition 3.4 (type *B* random environment; Figure 3) when {*ξ*^*n*^} is the *δ*_0_-Beta(*γ*, 2 − *α, α*)-coalescent (Figures 2–3; see also Figures C1 and C2 in Appendix C) or the *δ*_0_-Poisson-Dirichlet(*α*, 0)-coalescent (Figure 4; see also Figure E1 in Appendix E). When 1 ≤ *α* ≤ 3*/*2 (Figure 2) there is discrepancy between 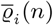 (black lines) and 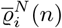 (coloured lines) regardless of the bound *ζ*(*N*). In Figure 2 the case *γ* = 1 (Figures 2a, C1a, C1b, C1c) is compared to the case *ζ*(*N*) = *N* log *N* (so that one would have *ζ*(*N*)*/N* → ∞ but lim sup _*N* →∞_ *m*_*N*_ *<* ∞ by Lemma 5.4 taking *ε*_*N*_ = *N* ^*α*−2^ log *N* since *κ* = 2, recall (53)), and the case *γ* = 1*/*(1 + m) (Figures C1d–C1f) to the case *ζ*(*N*) = *N*. The estimates 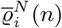 do not agree with 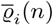 for the values of *α* considered when the population evolves according to Definition 3.4 (Figure 3); the agreement is somewhat better when the coalescent is the *δ*_0_-Poisson-Dirichlet(*α*, 0)-coalescent (Figure 4). In Figure 4 *ζ*(*N*) = *N* ^1*/α*^ log *N* so that *ζ*(*N*)*/N* ^1*/α*^ → ∞ as required for convergence to the *δ*_0_-Poisson-Dirichlet(*α*, 0)-coalescent. However, this means that as *α* approaches 0 the realised number of potential offspring any single individual may produce rapidly increases (see also Figure E2 in Appendix E for examples comparing estimates of E [*R*_*i*_(*n*)] as predicted by the *δ*_0_-Poisson-Dirichlet(*α*, 0)-coalescent for various parameter values). We have that *ε*_*N*_ is proportional to *N* ^*α*−2^ when Definition 3.3 holds as required by Lemma 5.7 (and 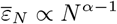 when Definition 3.4 holds as required by Lemma 5.16); thus in both cases as *α* decreases the probability of seeing large families in the ancestral process decreases; however, one expects the large families to be (on average) larger as *α* decreases (in Figure B2 in Appendix B we check that the algorithm for estimating E [*R*_*i*_(*n*)] correctly approximates *φ*_*i*_(*n*) (32)).

A *U*-shaped site-frequency spectra (excess of low-frequency and high-frequency derived alleles relative to predictions of the Kingman coalescent) is sometimes seen as a characteristic of multiple-merger coalescents, in particular Λ-coalescents [Freund et al., 2023]. Indeed, Λ-coalescents can predict *U*-shaped spectra [Birkner et al., 2013b]. Our results (see Figure B1 in Appendix B) do, however, reveal a broader range of shapes, including *L*-shaped spectra (displaying an excess of low-frequency alleles) similar to those predicted by a time-changed Kingman-coalescent as derived from a panmictic population evolving according to the Wright-Fisher model and continually increasing in size [Donnelly and Tavaré, 1995]. Thus, the statistical power of identifying multiple-merger coalescents from time-changed Kingman-coalescents using the site-frequency spectrum may have been somewhat diminished [Eldon et al., 2015, Koskela, 2018]. In particular, Figure B1 illustrates the effect of the cutoff parameter *γ* on the site-frequency spectrum, even for small *α* the site-frequency spectrum as predicted by the *δ*_0_-Beta(*γ*, 2 − *α, α*)-coalescent stays *L*-shaped if *γ* is small enough. In cases involving the Poisson-Dirichlet coalescents and reflecting high skewness of the offspring distribution (Figures A1, E2, and F2) the site-frequency spectrum exhibits a “plateau” (in particular Figures A1b, A1c, A1e, A1f, F2g, E2c). Recall that a Poisson-Dirichlet(*α*, 0)-coalescent is a Ξ-coalescent admitting simultaneous multiple mergers; when *n* blocks there can be up to ⌊*n/*2⌋ simultaneous mergers, thus as the skewness increases we can expect to see more simultaneous mergers. In contrast the Beta-coalescents admit only asynchronous mergers (at most one set of blocks merges at any given time). Moreover, in the cases where the “plateau” appears the relative length of external branches 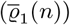 is increased.

#### 3.2.5 Comparing 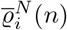 and 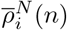

Recall 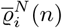 (approximations of annealed mean relative branch lengths) and 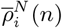 (approximations of quenched mean relative branch lengths) from (29), (30), (31). The approximations 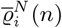 in Figures 2–4 result from ignoring the ancestral relations of the individuals in the population. An alternative way of approximating ‘branch lengths’ is to condition on the ancestral relations of the individuals in the population [Wakeley et al., 2012, Diamantidis et al., 2024]. This idea follows from the fact that as the population evolves ancestral relations are generated through inheritance. This means that the gene copies of a sample are related through a single complete gene genealogy. Given the *population ancestry*, the ancestral relations of the entire population of gene copies at the present and all previous times, and the identity of the sampled gene copies, the complete sample tree is fixed and known. Denote by 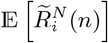 the *quenched* analogue of 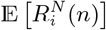 obtained by conditioning on the population ancestry, reading one sample tree from each ancestry, and averaging over independent ancestries to get an estimate of 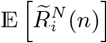. Our approach is conceptually similar to the approach of Wakeley et al. [2012] and Diamantidis et al. [2024], who consider gene genealogies within a fixed pedigree of diploid individuals. We consider a haploid population and work with population ancestries, read one sample tree from each ancestry, and average over population ancestries. We generate a complete sample tree forward-in-time before reading statistics off the tree. In our construction each individual occupies a level, and records the level occupied by its’ immediate ancestor. For example,

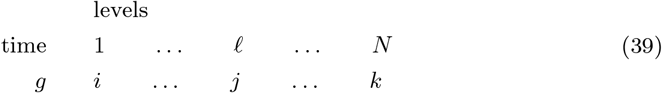

where the individual occupying level 1 resp. *ℓ* resp. *N* at time *g* derives from the individual occupying level *i* resp. *j* resp. *k* in the immediately previous generation. If the individual occupying level *ℓ* at time *g* contributes *x* surviving offspring, then *x* individuals at time *g* + 1 will be labelled (or “point to”) *ℓ*. In this way one can record the ancestry of the entire population. Given a complete sample tree we read statistics (branch lengths) off the tree, and repeat the procedure, starting from scratch with a new population ancestry.

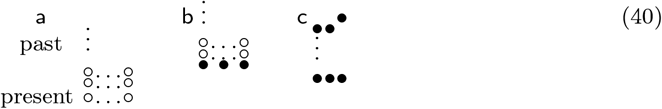

The process of sampling one quenched gene genealogy is illustrated in (40). We start by generating a population ancestry (40a; recalling (39) a population ancestry is a record of the ancestral relations of all the gene copies ◦ in the population), at some arbitrary time called “present” we draw a random sample (the sampled gene copies are shown as • in (40b)), once the identity of the sampled gene copies in the given population ancestry is known the sample tree (40c) is fixed, and one then simply reads the required statistic off the fixed tree; in (40c) the sampled gene copies and the gene copies ancestral to the sampled ones are shown as •. The quantity 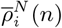 (31) is then obtained by averaging the statistics obtained by repeating the process illustrated in (40), each time starting with a new population ancestry generated independently of other population ancestries. Thus, 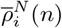 obtained in this way from a finite number of independent population ancestries is an approximation of 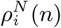. Here, we are concerned with a haploid panmictic population of constant size evolving according to a specific model of sweepstakes reproduction. The population could, however, include complex demography such as diploidy [Eldon, 2026], bottlenecks, population structure, selfing, dormancy (seedbanks), etc. Clearly, what the limit law is governing quenched gene genealogies as defined here, and what 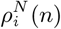 converges to as *N* → ∞, are open questions.

In the annealed (non-quenched) approach, we update the statistics generation by generation without ever generating a tree. With (*b*_1_, …, *b*_*m*_) denoting block sizes, starting with (*b*_1_, …, *b*_*n*_) = (1, …, 1), the annealed approach of sampling “branch lengths” 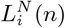 consists basically of two steps (recall we are working in discrete time),

1. 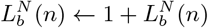 for *b* = *b*_1_, …, *b*_*m*_
2. sample *X*_1_, …, *X*_*N*_ according to a specific example of (18), merge blocks according to a multivariate hypergeometric using the sampled offspring numbers, and update block sizes

It is not *a priori* clear if 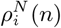 and 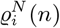 are different or not. Here we use simulations to compare 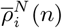 and 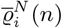 (recall (31)). We leave a mathematical investigation of gene genealogies conditional on the population ancestry in haploid populations evolving according to sweepstakes reproduction to future work. Section § 3.2.2 contains a brief description of the algorithm used to compute 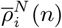.

In Figure 5 and Figure D1 in Appendix D we compare 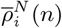 to 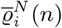 (blue lines) with *α* as shown and with the population evolving according to Definition 3.3 (Figure 5a, Figures D1a–D1d) and Definition 3.4 (Figure 5b, Figures D1e–D1f). Even though 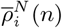 and 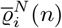 broadly agree, there is a visible difference. In Figure 5 and Figure D1 we take 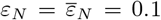 throughout since we are comparing predictions of the pre-limiting model and the focus is on seeing effects of conditioning on the population ancestry while keeping everything else the same between the two methods.

**Figure 5.**
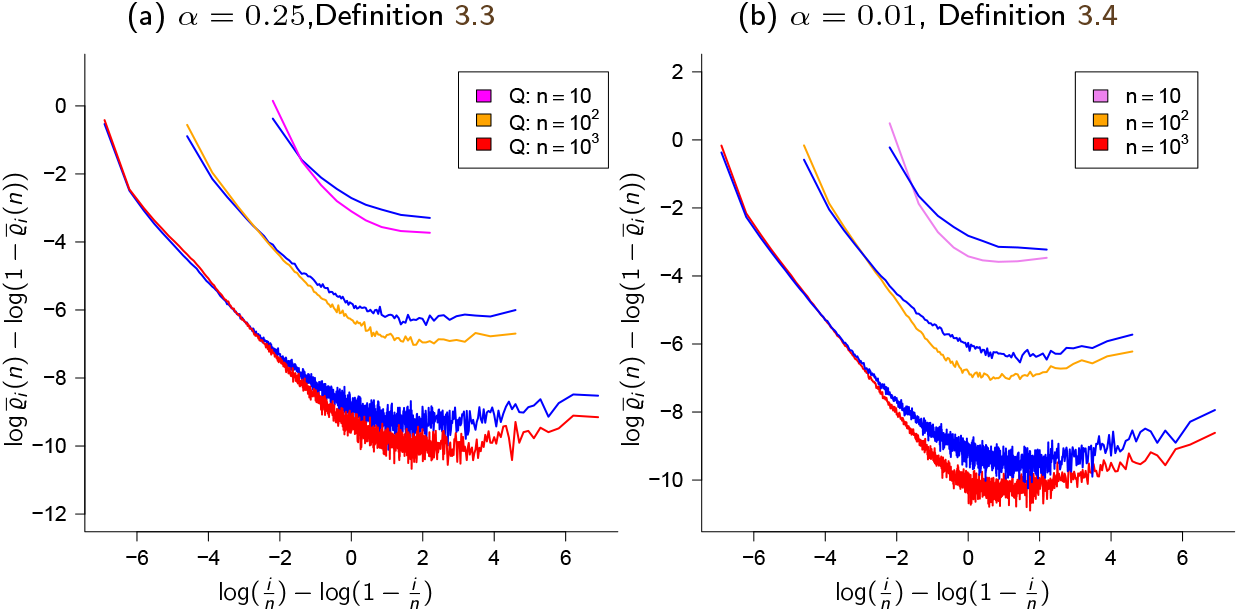
Quenched vs. annealed. Comparing 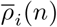 (estimates of mean relative branch lengths when conditioning on the population ancestry) and 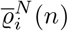 (blue lines; recall (31)) when the population evolves according to Definition 2.8 and Definition 3.3 (type *A* random environment) a; and Definition 3.4 (type *B* random environment) b; *N* = 10^3^, *α* as shown, *κ* = 2, *ζ*(*N*) = *N* ^1*/α*^ log *N* (a) and 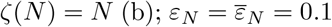; the approximations are from 10^4^ experiments. Sections 3.2.1 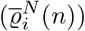 and 3.2.2 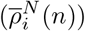 contain brief descriptions of the sampling algorithms. Further graphs are in Figure D1 in Appendix D. The scale of the ordinate (vertical axis) may vary between graphs.

It has been pointed out that increasing sample size may affect the site-frequency spectrum when the sample is from a finite population evolving according to the Wright-Fisher model (and so cause deviations from the one predicted by the Kingman coalescent), in particular when the sample size exceeds the effective size 1*/c*_*N*_ [Wakeley and Takahashi, 2003, Bhaskar et al., 2014, Melfi and Viswanath, 2018b,a]. We compare 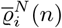 and 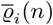 for different sample sizes to investigate if there is any noticeable effect of increasing sample size. Recalling that 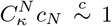 (recall (24)) we conjecture it is not surprising to see little effect of increasing sample size when 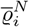 and 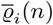 otherwise broadly agree (e.g. Figures C1c and 2b). Further graphs are in Figure C1 in Appendix C.

### 3.3 Time-changed Beta- and Poisson-Dirichlet-coalescents

The population size of natural populations may vary over time. Here we briefly consider fluctuations (primarily with population growth in mind) in the spirit of [Freund, 2020]. Freund studies Cannings models with large enough fluctuations in population size to lead to time-changed Λ-coalescents. See [Birkner et al., 2009] for an alternative approach with a focus on recurrent bottlenecks. Suppose 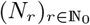 is a sequence of population sizes where *N*_*r*_ is the population size *r* generations into the past with *N*_0_ ≡ *N*, and such that 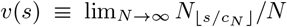 where *s >* 0 and *c*_*N*_ is the coalescence probability (Definition 2.2) for the fixed-population size case [Freund, 2020, Equation 4]. In particular, when the underlying population model is the one of [Schweinsberg, 2003] as giving rise to the Beta(2 −*α, α*)-coalescent with 1 ≤ *α <* 2, the time-change function *G*(*t*) is of the form 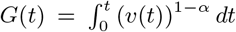 [Freund, 2020, Theorem 3]. It follows that gene genealogies will be essentially unaffected by variations in population size whenever the fluctuations satisfy [Freund, 2020, Equation 4] and *α* is close to 1.

A time-changed *δ*_0_-Beta(*γ*, 2 − *α, α*)-coalescent with 0 *< α <* 2 and whose time-change *G*(*t*) is independent of *α* follows almost immediately from Theorems 3.5 and 3.6 and [Freund, 2020, Lemma 4]. § 5.4 contains a proof of Theorem 3.8.

#### Theorem 3.8

(A time-changed *δ*_0_-Beta(*γ*, 2 − *α, α*)-coalescent). *Suppose a haploid population evolves according to Definitions 2.8 and Definition 3.3 when* 1 ≤ *α <* 2, *and Definition 3.4 when* 0 *< α <* 1. *Suppose* (*N*_*r*_)_*r*∈ℕ_ *is a sequence of population sizes with N*_*r*_ *the population size r generations into the past, and where*

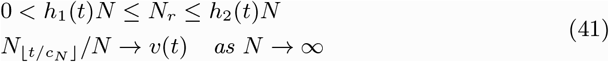

*for bounded positive functions h*_1_, *h*_2_, *v* : [0, ∞) → (0, ∞). *Then {ξ*^*n,N*^ (⌊*t/c*_*N*_ ⌋); *t* ≥ 0} *converges to* {*ξ*^*n*^(*G*(*t*)); *t* ≥ 0} *where* {*ξ*^*n*^(*t*);*t* ≥ 0} *is the δ*_0_*-Beta*(*γ*, 2 − *α, α*)*-coalescent with* 0 *< γ* ≤ 1 *and* 0 *< α <* 2, *and* 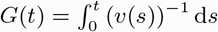.

A growing population is sometimes modelled as increasing exponentially in size [Donnelly and Tavaré, 1995]. Figure 6 (see also Figure F1 in Appendix F) holds examples of 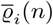 (recall (31)) for the time-changed *δ*_0_-Beta(*γ*, 2 − *α, α*)-coalescent when *N*_*r*_ = ⌊*N*_*r*−1_ (1 − *ρc*_*N*_)⌋, so that 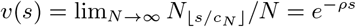 and *c*_*N*_ being the coalescent probability for the fixed-size case. Write *τ*_*m*_ for the time when {*ξ*^*n*^ (*G*(*t*)); *t ≥* 0} reached *m* blocks (with *τ*_*n*_ = 0). The distribution of the time during which there are *m* lineages is

**Figure 6.**
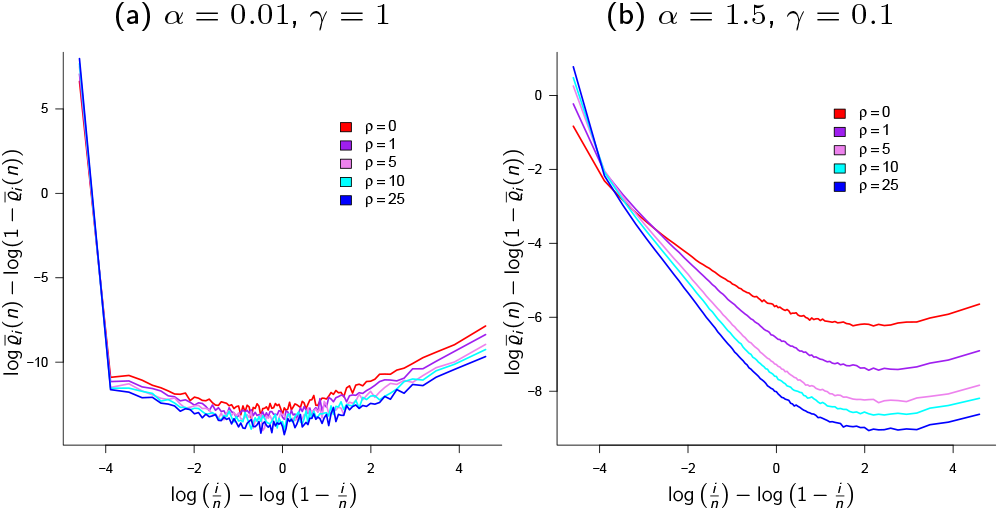
Approximations 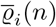 (estimates of mean relative branch lengths, recall (31)) predicted by the time-changed *δ*-Beta(*γ*, 2 − *α, α*)-coalescent with time-change 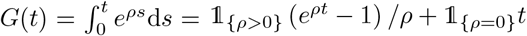 with *ρ* as shown, *κ* = 2, *c* = 1, *n* = 100. The scale of the ordinate (*y*-axis) may vary between the graphs; results from 10^5^ experiments. Figure F1 in Appendix F contains further graphs. The scale of the ordinate (vertical axis) may vary between graphs.

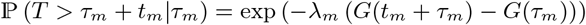

where *λ*_*m*_ is the total merging rate of {*ξ*^*n*^} when *m* blocks [Freund, 2020, Remark 7, Equation 23]. It follows that the time *t*_*m*_ during which {*ξ*^*n*^ (*G*(*t*)); *t ≥* 0} has *m* lineages is given by (for *ρ >* 0)

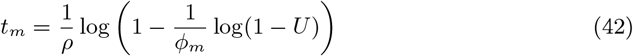

where *U* is a random uniform on (0, 1), *ϕ*_*m*_ = (1*/ρ*)*λ*_*m*_ exp(*ρτ*_*m*_), and *τ*_*n*_ = 0. As for the case when {*ξ*^*n*^} is the Kingman coalescent [Donnelly and Tavaré, 1995], the time-change acts to increases 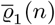 when {*ξ*^*n*^} is a *δ*_0_-Beta(*γ*, 2 ™ *α, α*)-coalescent.

A time-changed *δ*_0_-Poisson-Dirichlet(*α*, 0)-coalescent follows almost immediately from Case 3 of Theorem 3.5 and [Freund, 2020, Lemma 2]. § 5.5 contains a proof of Theorem 3.9.

#### Theorem 3.9

(Time-changed *δ*_0_-Poisson-Dirichlet(*α*, 0)-coalescent). *Suppose a haploid population evolves according to Definition 2.8 and type A random environment (Definition *3*.3) with* 0 *< α <* 1. *Suppose* (*N*_*r*_)_*r*∈ℕ_ *is as in Theorem 3.8 (recall* (41)*) and such that* 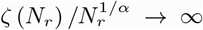 *as N*_*r*_ → ∞ *for all r* ∈ ℕ. *Then {ξ*^*n,N*^ (⌊*t/c*_*N*_ ⌋); *t* ≥ 0} *converges to* {*ξ*^*n*^ (*G*(*t*)); *t* ≥ 0} *where* {*ξ*^*n*^ (*t*); *t* ≥ 0} *is a δ*_0_*-Poisson-Dirichlet*(*α*, 0)*-coalescent, and* 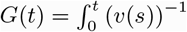 *with v as in* (41).

Figure 7 (see also Figure F2 in Appendix F) contains examples of 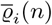 (recall (31)) for the time-changed *δ*_0_-Poisson-Dirichlet(*α*, 0)-coalescent. As for the time-changed *δ*_0_-Beta(*γ*, 2−*α, α*)-coalescent, the time-change applied to the *δ*_0_-Poisson-Dirichlet(*α*, 0)-coalescent, corresponding to exponential population growth with rate *ρ*, acts to extend 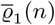.

**Figure 7.**
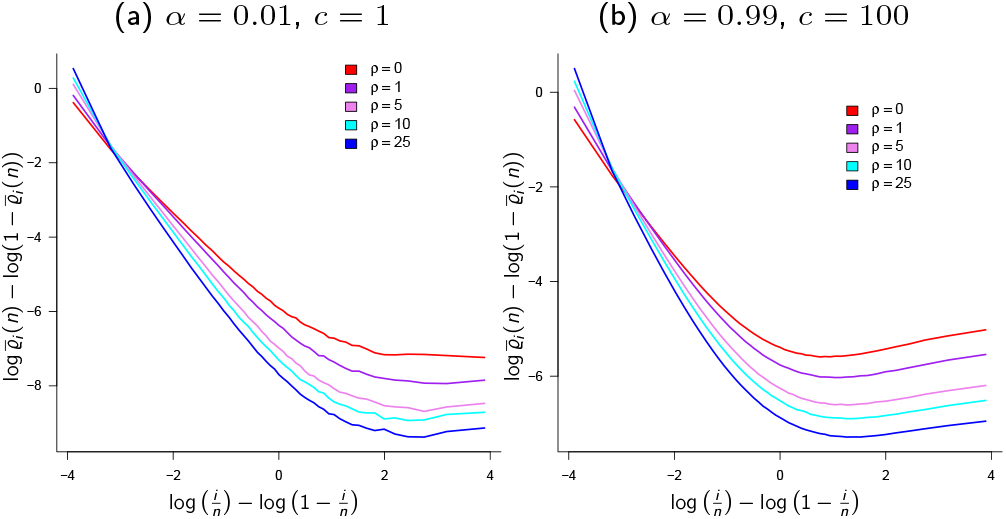
Approximations 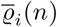 (estimates of mean relative branch lengths, recall (31)) predicted by the time-changed *δ*-Poisson-Dirichlet(*α*, 0)-coalescent with time-change 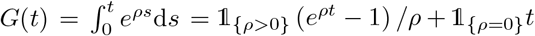 with *ρ* as shown, *n* = 50, *κ* = 2; results from 10^6^ experiments. The scale of the ordinate (vertical axis) may vary between graphs. Further graphs are in Figure F2 in Appendix F.

## Conclusion

Our main results are *(i)* continuous-time coalescents derived from population genetic models of sweepstakes reproduction; the coalescents are either the Kingman coalescent, or specific families of the *δ*_0_-Beta- or *δ*_0_-Poisson-Dirichlet-coalescents (see Appendix A for an example of a Ξ-coalescent without an atom at zero); *(ii)* in contrast to [Schweinsberg, 2003] (recall (15) in Theorem 2.9), time in (almost) all our coalescents is measured in units proportional to either *N/* log *N* or *N* generations (see Appendix A for an exception); *(iii)* time-changed *δ*_0_-Beta- or *δ*_0_-Poisson-Dirichlet-coalescents where the time-change is independent of *α*; *(iv)* when {*ξ*^*n,N*^} is in the domain of attraction of the *δ*_0_-Beta(*γ*, 2 − *α, α*)-coalescent the 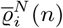 are not matched by 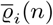 (recall (31)), in particular for small values of *α*; *(v)* even though 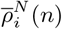 are broadly similar to 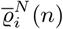 for the models considered here, the difference between them may indicate that 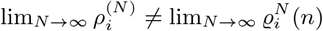.

We investigate multiple-merger coalescents where the driving measure Ξ is of the form

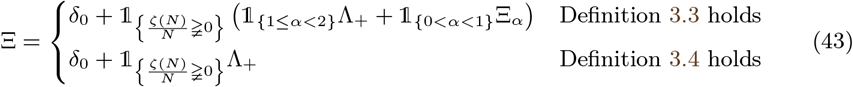

where Λ_+_ is a finite measure on (0, 1] without an atom at zero. In our formulation Ξ always retains an atom at zero. The coalescents derived here are based on a construction where most of the time small (relative to the population size) families occurs, but occasionally the environment favours the generation of (at least one) large family. Sweepstakes reproduction is incorporated as in Definition 3.3 (type *A* random environment) and Definition 3.4 (type *B* random environment) with an upper bound on the number of potential offspring any one individual may produce. Sweepstakes reproduction, for example in broadcast spawning marine organisms, may occur through chance matching of favorable environmental conditions and reproduction [Hedgecock and Pudovkin, 2011, Li and Hedgecock, 1998, Hedgecock et al., 1982, Beckenbach, 1994]. Random environments similar to the ones in Definitions 3.3 and 3.4 seem natural for modeling sweepstakes reproduction.

In our formulation (recall (18)) the upper bound *ζ*(*N*) on the number of potential offspring determines if the limiting coalescent admits multiple-mergers (recall (43)). For comparison, Λ = 1_{*α*≥2}_*δ*_0_ + 1_{1≤*α<*2}_Λ_+_ absent a bound on the number of potential offspring [Schweinsberg, 2003]. Moreover, we show that restricting *ζ*(*N*) to the order of the population size is sufficient to obtain a multiple-merger coalescent. In contrast, convergence to the original (complete) Beta(2 ࢤ *α, α*)-coalescent requires the (slightly) stronger assumption of *ζ*(*N*)*/N* → ∞. Applying an upper bound appears reasonable for at least some highly fecund populations. Considering broadcast spawners, we see potential offspring as representing fertilised eggs (or some later stage in the development of the organisms before maturity is reached). It is plausible that the number of fertilised eggs produced by an arbitrary individual for (at least some) broadcast spawners may be at most a fraction of the population size. Our numerical results show that an upper bound on the number of potential offspring markedly affects the predicted site-frequency spectrum (see Appendix B for examples comparing *φ*_*i*_(*n*) from (32) between *δ*_0_-Beta(*γ*, 2 − *α, α*)-coalescents). Thus, even though our extensions ((18), Theorems 3.5 and 3.6) of (14) are relatively straightforward, the implications for inference of our results are significant.

Theorems 3.5 and 3.6 combine to yield a continuous-time *δ*_0_-Beta(*γ*, 2 − *α, α*)-coalescent where 0 *< γ* ≤ 1 and 0 *< α <* 2, with the understanding that the population model when 1 ≤ *α <* 2 is different from the one when 0 *< α <* 1. When 1 ≤ *α <* 2 the population evolves as in Definition 3.3 (type *A* random environment), and according to Definition 3.4 (type *B* random environment) when 0 *< α <* 1. Thus, we have extended the Beta-coalescent [Schweinsberg, 2003] to include an atom at zero, a truncation point (*γ*; see also Chetwynd-Diggle and Eldon [2026]), and with the range of *α* extended from [1, 2) to (0, 2). However, the number of parameters involved has now increased from one (*α*) to four (*c, γ, α, κ*), with the mean *m*_∞_ being a function of *κ* (recall (35)). Moreover, when in the domain of attraction of the *δ*_0_-Beta(*γ*, 2 − *α, α*)-coalescent E [*X*_1_] *<* ∞ regardless of the value of *α*; under (14) E [*X*_1_] = ∞ when 0 *< α* ≤ 1. For the *δ*_0_-Beta(*γ*, 2 − *α, α*) coalescent the unit of time is proportional to *N/* log *N* generations when *κ* = 2, and to *N* generations when *κ >* 2. For comparison, time in the original complete Beta-coalescent is measured as shown in (15) [Schweinsberg, 2003, Lemma 16; Lemma 13]. Effective population size (*N*_*e*_) is often taken as 1*/c*_*N*_, and so would be proportional to at least *N/* log *N* for our models. One can ask what assumptions on the population size (*N*) and the mutation rate (*µ*) are required to recover the mutations in a sample of DNA sequences used to obtain an estimate of *α*. Using the *N* ^*α*−1^ timescale (1 *< α <* 2), it can be argued that recovering observed mutations requires strong assumptions on *N* (and/or *µ* if unknown) when the estimate of *α* is near 1 [Chetwynd-Diggle and Eldon, 2026]. Moreover, our results show that sweepstakes reproduction does not necessarily imply a small *N*_*e*_*/N* ratio [Waples, 2016, Árnason, 2004].

We obtain continuous-time *δ*_0_-Poisson-Dirichlet(*α*, 0)-coalescents with time measured in units proportional to at least *N/* log *N* generations. Sagitov [Sagitov, 2003, § 3] discusses an example of a Poisson-Dirichlet-coalescent deduced from a compound multinomial distribution. Combining Definition 2.8 and (14) when 0 *< α <* 1 yields a Poisson-Dirichlet(*α*, 0)-coalescent without rescaling time 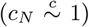 [Schweinsberg, 2003]. However, convergence to the *δ*_0_-Poisson-Dirichlet(*α*, 0)-coalescent requires that *ζ*(*N*)*/N* ^1*/α*^ → ∞, imposing a strong assumption on the distribution of potential offspring (recall 0 *< α <* 1). We leave it for later to investigate the coalescents that may arise when *ζ*(*N*)*/N* ^1*/α*^ converges as *N*→ ∞ to some constant *K >* 0; we conjecture (see Remark 5.12) that the resulting coalescent may be a form of an ‘incomplete Poisson-Dirichlet’ coalescent, similar to the case *ζ*(*N*)*/N* → *K* required for convergence to an incomplete Beta-coalescent. Other scalings of *ζ*(*N*) may also lead to new non-trivial coalescents.

The introduction of the coalescent, a sample-based approach for investigating evolutionary histories of natural populations moved population genetics forward. The coalescent enables efficient generation of gene genealogies, in particular when applying recent advances in algorithm design [Baumdicker et al., 2021, Kelleher et al., 2016]. However, a coalescent-based inference is only helpful when the coalescent correctly approximates the trees generated by the corresponding ancestral process. Here, we check this approximation by comparing functionals (recall (30) and (31)) of {*ξ*^*n*^} and {*ξ*^*n,N*^} when the population is haploid and panmictic of constant size and evolving under sweepstakes reproduction. Given the poor agreement between 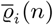 and 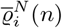 in some cases, in particular when the law for the number of potential offspring is strongly skewed, the usefulness of coalescent-based inference is in doubt in this case. Multiple-merger coalescents have been applied to various broadcast-spawning marine organisms [Árnason and Halldórsdóttir, 2015, Árnason et al., 2023, Vendrami et al., 2021, Niwa et al., 2016, Barfield et al., 2023], outbreaks of tuberculosis [Menardo et al., 2021], and even to the propagation of cancer cells[Kato et al., 2017]. Moreover, multiple-merger coalescents may be suitable for investigating plants and (crop-infesting) fungi that reproduce by distributing huge numbers of seeds or spores [Minadakis et al., 2025, Jigisha et al., 2025]. Thus, the models studied here, and the resulting multiple-merger coalescents may be relevant for quite a broad range of study systems. Moreover,

One might want to extend the models applied here to diploid populations (where gene copies always occur in pairs regardless of the reproduction mechanism), and to include complex demography such as recurrent bottlenecks [Birkner et al., 2009]. Multiple-merger coalescents derived from population models of sweepstakes reproduction in diploid populations tend to be Ξ-coalescents [Möhle and Sagitov, 2003, Birkner et al., 2018, 2013a]. Site-frequency spectra predicted by Ξ-coalescents can be different from the ones predicted by Λ-coalescents [Blath et al., 2016, Birkner et al., 2013b]. As we have seen (Theorem 3.6), coalescents derived for haploid panmictic populations of constant size and evolving according to sweepstakes may also be continuous-time Ξ-coalescents (the *δ*_0_-Poisson-Dirichlet(*α*, 0)-coalescent). In any extensions of the models considered here, one should compare 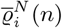 to 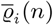, and to 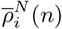. Should any observed difference between 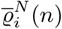 and 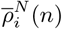 not be an artifact of the finite population size used in simulations such that 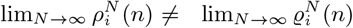 then we may have to rethink gene genealogies from scratch.

## 5 Proofs

This section contains proofs of Theorem 3.5 in § 5.2, of Theorem 3.6 in § 5.3, Theorem 3.8 in § 5.4, and of Theorem 3.9 in § 5.5. First, in § 5.1, we verify (12).

### 5.1 Verifying (12)

In this section we check that (12) holds. For *x, a* ∈ ℝ and *m* ∈ {0, 1, 2, …} define [*x*]_0;*a*_ = 1, and

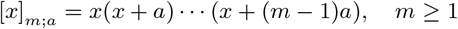

so that [*α*]_*r*™1;*α*_= *α*^*r*−1^(*r* − 1)! and [1]_*m*™1;1_ = (*m* − 1)!. Then, with *b* ≥ 2, 0 *< α <* 1, and *k*_1_ + *k*_*r*_ = *b* where {*k*_1_, …, *k*_*r*_} is a partition of *b*, the exchangeable probability function is

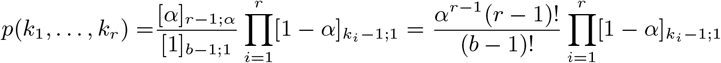

[Pitman, 1995, Proposition 9, Equation 16], where, recalling (1), 912

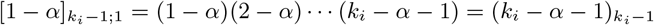

Following [Schweinsberg, 2003] write

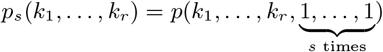

where

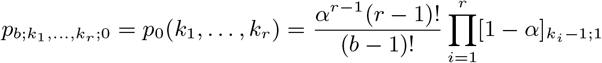

and

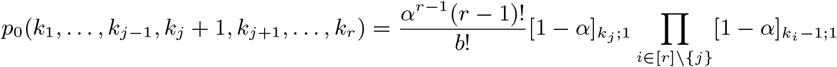

Then, by [Schweinsberg, 2003, page 137],

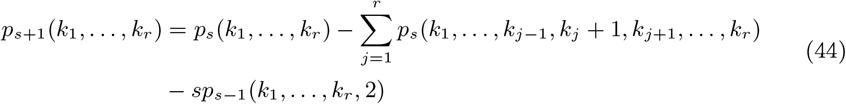

By (44) with *s* = 0, noting that [1 − *α*]_*k*;1_ = [1 − *α*]_*k*−1;1_(*k* − *α*),

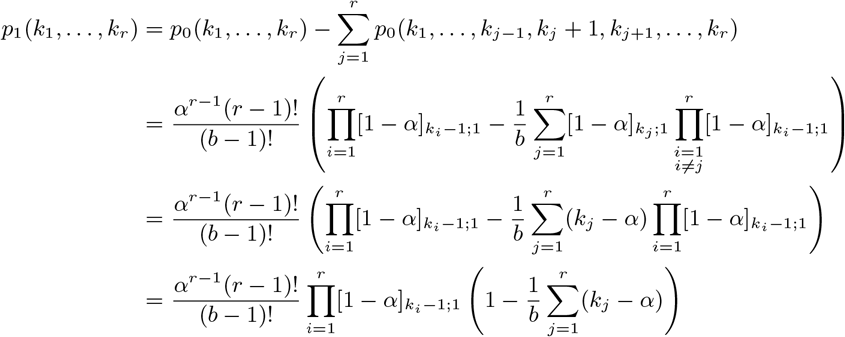

We then have, recalling *k*_1_ + · · · + *k*_*r*_ = *b*,

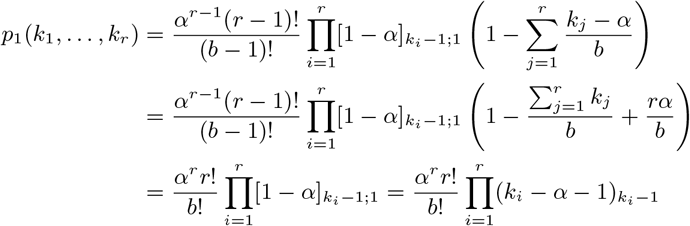

Similarly one uses (44) to check (12) for *s* = 1, hence to (12) by induction over *s*.

### 5.2 Proof of Theorem 3.5

In this section we give a proof of Theorem 3.5. Therefore, we consider a haploid population evolving according to Definitions 2.8 and 3.3 (type *A* random environment) with law (18) on the number of potential offspring produced by each individual. To briefly orient the reader (the proofs can always be skipped on first reading to get an overview of the lemmas leading to the theorem), Proposition 5.1 and Lemmas 5.2 and 5.3 collect useful results for deriving the limits of 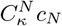 (recall 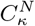 from (24)) in Lemma 5.7. Lemma 5.10 proves Case 1 of Theorem 3.5 by checking the condition given in (7). The proof of Case 2 of Theorem 3.5 involves checking the conditions of Proposition 2.3, Lemma 5.8 identifies the Λ_+_-measure (Condition (6c) of Proposition 2.3), and Lemma 5.9 checks that simultaneous mergers vanish as *N* → ∞ (Condition 6b of Proposition 2.3). The proof of Case 3 of Theorem 3.5 follows the proof of [Schweinsberg, 2003, Theorem 4d]. Lemmas 5.4 (*m*_∞_ *<* ∞) and 5.6 (*S*_*N*_ */*(*Nm*_*N*_) → 1 almost surely) are ancillary results required for checking the convergence of 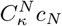. Note in particular that, since *m*_∞_ *<* ∞ in our framework, we can use the same method for *α* = 1 as for 1 *< α <* 2.

#### Proposition 5.1

([Chetwynd-Diggle and Eldon, 2026]; Lemma 7.2). *Suppose G and H are positive functions on* [1, ∞), *H be monotone increasing, G monotone decreasing, and ∫ HG*^′^ *exists. Then for ℓ, m* ∈ ℕ

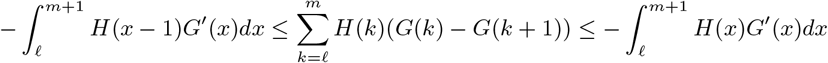

We will use Proposition 5.1 to obtain bounds on 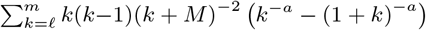; when we do so we intend to replace *M*, *ℓ*, and *m* as later specified to identify the limit of 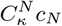 as *N* → ∞.

#### Lemma 5.2

(Bounds on ∑_*k*_ *k*(*k*−1)(*k* + *M*)^−2^ (*k*^−*a*^ −(*k* + 1)^−*a*^)). *For k* ∈ ℕ, *with M* ≫ 1, *a >* 0, 1 ≤ *ℓ* ≪ *m all fixed; write r*_*k*_(*a*) = *k*(*k* − 1)(*k* + *M*)^−2^(*k*^−*a*^ − (*k* + 1)^−*a*^).

1. *When* 0 *< a <* 1 *we have*

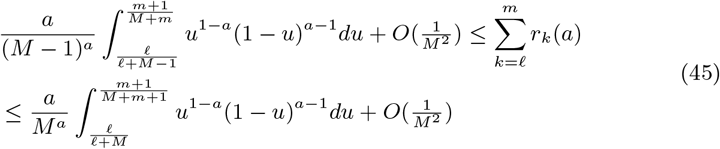
2. *When a* = 1 *we have*

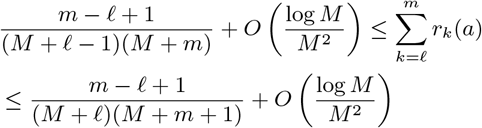
3. *When* 1 *< a <* 2 *the same bounds as in* (45) *for Case 1 hold*.
4. *When a* = 2

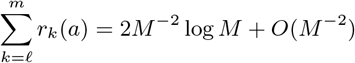
5. *When* 2 *< a*

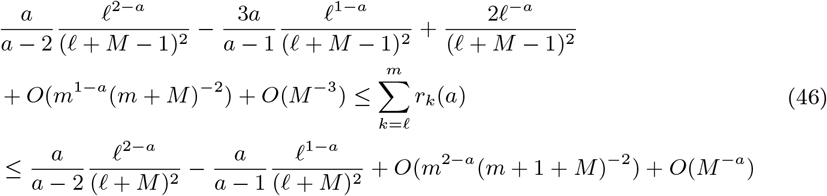

*Proof of Lemma 5.2*. Take *H*(*x*) = *x*(*x* − 1)(*x* + *M*)^−2^ and *G*(*x*) = *x*^−*a*^ in Proposition 5.1 to obtain

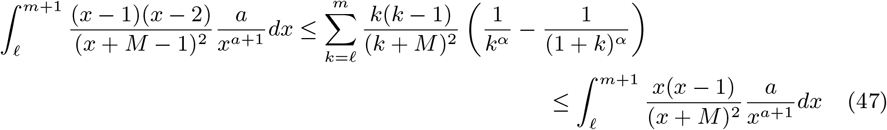

Now the task is to evaluate the integrals in (47). One can write

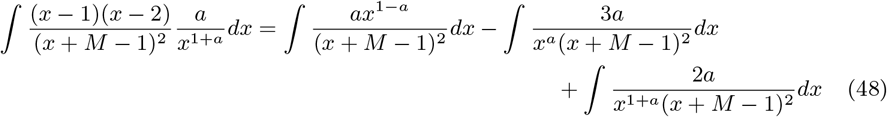

and

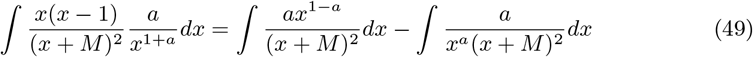

and evaluate each integral on the right in (48) and (49) separately using standard integration techniques.

When *a* = 1 using (49) and integration by partial fractions we see

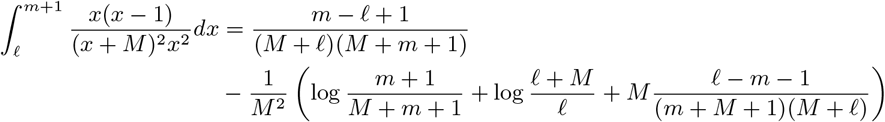

The lower bound in Case 2 is obtained in the same way using (48).

When 0 *< a <* 1 (Case 1) or 1 *< a <* 2 (Case 3) the substitution *y* = *M/*(*x* + *M*) (so that *x* = *My*^−1^ − *M* and *dx* = −*My*^−2^*dy*) gives

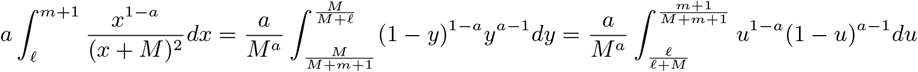

The same substitution and integration by parts gives

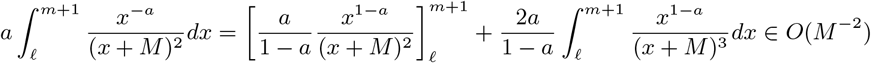

A similar calculation for the lower bound in (47) gives Case 1 and (45) for 1 *< a <* 2 (Case 3).

When *a* = 2 (Case 4) (49) and integration by partial fractions gives

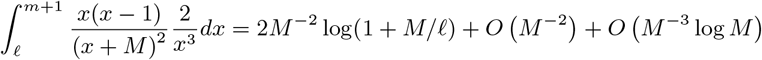

When *a >* 2 (Case 5) integration by parts and the substitution *y* = *M/*(*x* + *M*) as in the case 1 *< a <* 2 give the result. When 2 *< a <* 3 integration by parts gives

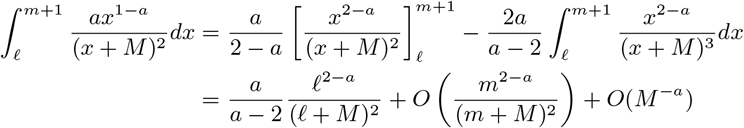

using the substitution *y* =*M/*(*x* + *M*) for the error term *O*(*M* ^−*a*^). When *a* ≥ 3 we simply repeat the calculation on ∫ *x*^2−*a*^(*x* + *M*)^−3^*dx* to see that we retain the same leading term. Considering ∫ *x*^−*a*^(*x* + *M*)^−2^*dx* we again apply integration by parts to see

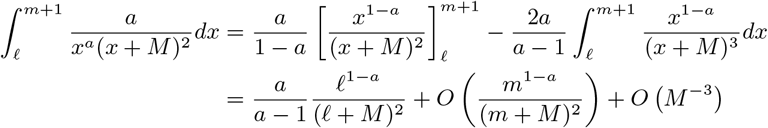

The remaining terms of Case 5 are identified in a similar way (here we omit the details).

#### Lemma 5.3

(Bounds on *k*^−*a*^ − (*k* + 1)^−*a*^). *For any* 0 *< a* ≤ 1 *and k* ∈ ℕ *we have*

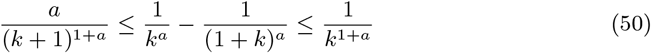

*When a* ≥ 1 *and k* ≥ 2 *we have*

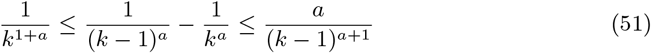

*Proof of Lemma 5.3*. Suppose 0 *< a* ≤ 1. Then

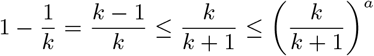

so that *k*^−*a*^ − *k*^−1−*a*^ ≤ (*k* + 1)^−*a*^. Rearranging terms gives the upper bound in (50).

Considering the lower bound in (51) we see, for any *a* ≥ 1,

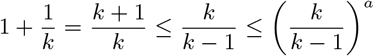

Then *k*^−*a*^ + *k*^−1−*a*^ ≤ (*k* − 1)^−*a*^ and rearranging terms gives the bound.

For the upper bound in (51) we recall the inequality

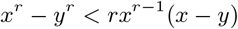

for any *x > y >* 0 and any rational *r >* 1 [Hardy, 2002, §74, Equation 9]. Suppose *x* = *k* + 1 and *y* = *k*. We see

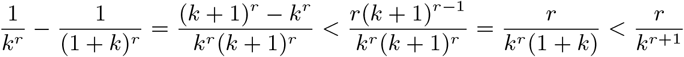

Since for any fixed *x >* 0 the function *r* ↦ *x*^−*r*^ is continuous in *r*, and the rationals are dense in ℝ, the upper bound in (51) follows.

The lower bound in (50) follows by similar calculations, using the inequality *sx*^*s*−1^(*x* − *y*) *< x*^*s*^ − *y*^*s*^ for 0 *< s <* 1 rational and any *x, y* ∈ ℝ with *x > y >* 0 [Hardy, 2002, §74, Equation 10, pp. 144].

The following lemma gives a condition on (*ε*_*N*_)_*N*_ (recall *ε*_*N*_ is the probability of event *E* in type *A* random environment defined in Definition 3.3) for the mean E [*X*_1_] = *m*_*N*_ from (23c) to be finite for all *N*, and for *m*_∞_ = lim_*N* →∞_ *m*_*N*_ to be finite.

#### Lemma 5.4

(Finite *m*_∞_). *Under the conditions of Theorem 3.5, if*

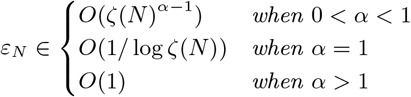

*then, with X the random number of potential offspring of an arbitrary individual*,

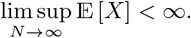

*Proof of Lemma 5.4*. Recalling event *E* from Definition 3.3 it is straightforward to check that lim sup_*N*→∞_ E [*X*|*E*] *<* ∞ whenever *α >* 1. Therefore, it is sufficient to consider E [*X*|*E*] for the case 0 *< α* ≤ 1. First with 0 *< α <* 1 and using the upper bound in (50) in Lemma 5.3 and recalling 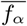 from (20a),

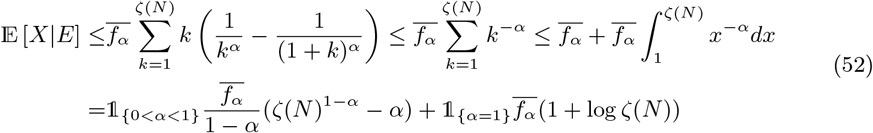

With event *E*^c^ as in Definition 3.3 and E [*X*] = E [*X*|*E*] *ε*_*N*_ + E [*X*|*E*^c^] (1 − *ε*_*N*_) the lemma follows.

#### Remark 5.5

(Lemma 5.4 using tail probability). *We see, with* 0 *< α <* 1

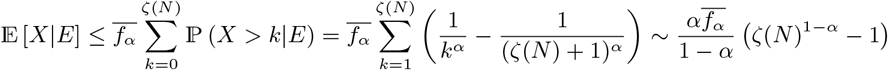

*in agreement with the choice of ε*_*N*_ *in Lemma 5.4*.

#### Lemma 5.6

(*S*_*N*_ */*(*Nm*_*N*_) → 1 almost surely). *Suppose X*_1_, …, *X*_*N*_ *are independent positive integer-valued random variables with X*_1_, …, *X*_*N*_ ▷ L(*α, ζ*(*N*)) *(recall* (19)*) with probability ε*_*N*_, *and X*_1_, …, *X*_*N*_ ▷ L(*κ, ζ*(*N*)) *with probability* 1 − *ε*_*N*_, *where* 1 ≤ *α <* 2 *and κ* ≥ 2 *fixed. Recall* 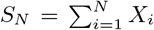 *from* (23a) *and* E [*X*_1_] = *m*_*N*_ *from* (23c) *in Definition 3.2, and choose ε*_*N*_ *as in Lemma 5.4 such that* lim sup_*N*→∞_ *m*_*N*_ *<* ∞. *Then S*_*N*_ */*(*Nm*_*N*_) → 1 *almost surely as N* → ∞.

*Proof of Lemma 5.6*. Let 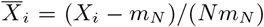 for *i* ∈ [*N*]. The 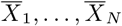 are i.i.d. and 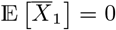. Then 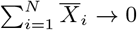 almost surely [Etemadi, 1981]. Hence, 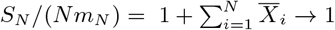 almost surely as *N* → ∞.

We verify (28) under the conditions of Case 2 of Theorem 3.5. Recall *c*_*N*_ from Definition 2.2 and (4).

#### Lemma 5.7

(*c*_*N*_ under Theorem 3.5). *Suppose the conditions of Theorem 3.5 hold with* 1 ≤ *α <* 2, *κ* ≥ 2, *and c >* 0 *all fixed, and that ζ*(*N*)*/N* ≩ 0. *Let L* ≡ *L*(*N*) *be a function of N such that L/N* → 0, *and*

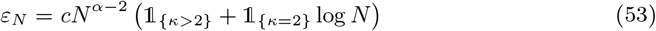

*Then, with κ*+2 *< c*_*κ*_ *< κ*^2^ *when κ >* 2 *and with* 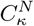 *as in* (24), *γ as in* (26a), *B*(*γ*, 2 −*α, α*) *as in Definition 2.4*, 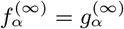 *and* 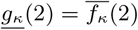, *we have* 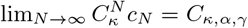 *where C*_*κ,α,γ*_ *as in* (26b) *and m*_∞_ *as in* (26d), *i.e*.

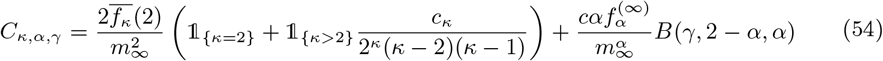

Choosing *ε*_*N*_ (recall Definition 3.3 of type *A* random environment) as in Lemma 5.7 gives *ε*_*N*_ */c*_*N*_ ≫ 0 in all cases and so we expect with high probability to see at least one event *E* (the event *X*_1_, …, *X*_*N*_ ▷ L(*α, ζ*(*N*))) over a period of time proportional to 1*/c*_*N*_ generations.

*Proof of Lemma 5.7*. Choosing *ε*_*N*_ as in (53) yields a finite *m*_∞_ by Lemma 5.4 and almost sure convergence of *S*_*N*_ */*(*Nm*_*N*_) to 1 by Lemma 5.6. Let, for 0 *< δ <* 1 fixed and such that (1 − *δ*)*m*_*N*_ *>* 1,

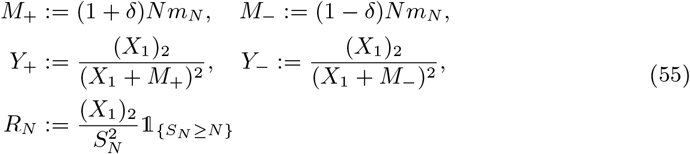

where E [*X*_1_] = *m*_*N*_ and *m*_∞_ = lim_*N* →∞_ *m*_*N*_ as in (23c) in Definition 3.2. By Theorem 2.9 (recall *c*_*N*_ from Definition 2.2)

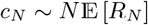

Adapting the arguments of [Schweinsberg, 2003, Lemma 13] and using Lemma 5.6 we have for any fixed *ϵ >* 0 (and assuming *N* is large enough that *N* − 1 ≈ *N*)

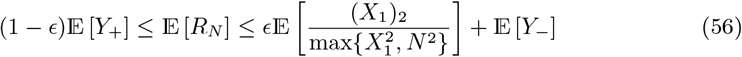

With 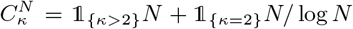 as in (24) we now see that it suffices to consider lower bounds on 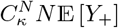 and upper bounds on 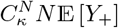 and to check that

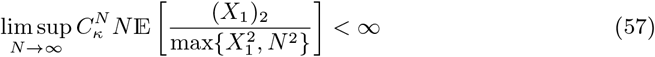

We check that (57) holds. Since (*X*_1_ + *N*)^2^ ≤ 4 max 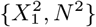 we see

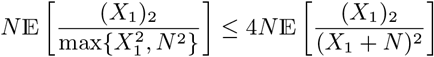

and so to (57) combining Lemma 5.2 and (53).

Write *r*_*k*_(*a, M*) ≡ (*k*)_2_(*k* + *M*)^−2^(*k*^−*a*^ − (*k* + 1)^−*a*^). Equation (54) will follow from Lemma 5.2 after substituting for *ℓ* and *m*. When *α* = 1

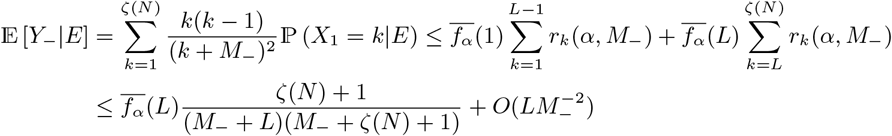

after substituting for *ℓ* and *m* in Case 2 of Lemma 5.2 so that by (53) when *α* = 1

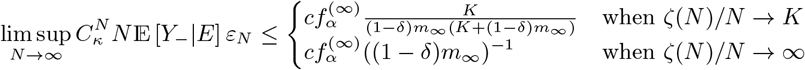

In the same way we get

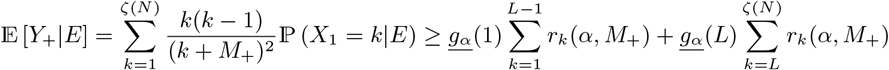

and when *α* = 1 we have the lower bound

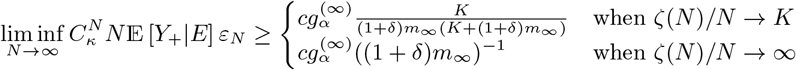

Write

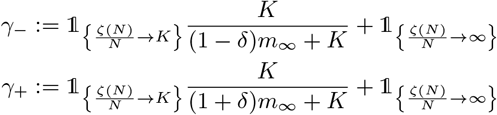

When 1 *< α <* 2 combining Case 3 of Lemma 5.2 with *ℓ* = 1 and *m* = *ζ*(*N*) and (53) we get the bounds

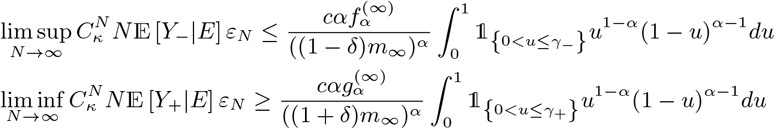

When *κ* = 2 combining Case 4 of Lemma 5.2 with *ℓ* = 2 and *m* = *ζ*(*N*) and (53) we get the bounds

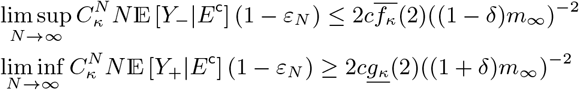

When *κ >* 2 combining Case 5 of Lemma 5.2 with *ℓ* = 2 and *m* = *ζ*(*N*) and (53) we get

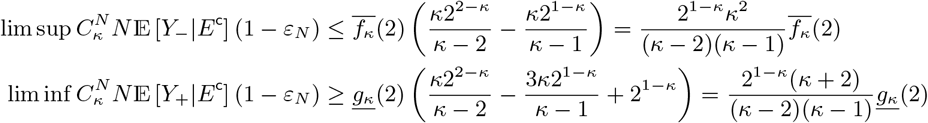

noting that, when *ℓ* = 2, the leading terms in the lower resp. upper bound in (46) simplify to

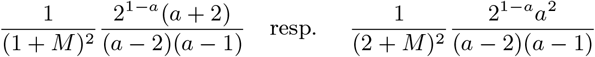

The lemma now follows from the assumption 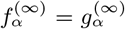 and 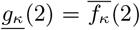 and taking *ϵ* and *δ* to 0. The proof of Lemma 5.7 is complete.

#### Lemma 5.8

(Identifying the Λ_+_-measure). *Under the conditions of part 2 of Theorem 3.5 with γ as in* (26a), *c from* (53), 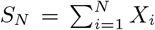 *as in* (23a), 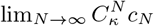 *as in* (54), 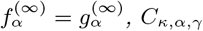 *as in* (26b), *m*_∞_ = lim_*N* →∞_ E [*X*_1_] *as in* (23c),

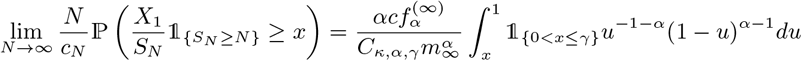

*Proof of Lemma 5.8*. The proof follows the proof of [Schweinsberg, 2003, Lemma 14]. Let *ϵ >* 0 and *δ >* 0 both be fixed with (1 − *δ*)*m*_*N*_ *>* 1. Then, by Lemma 5.6 we can take *N* large enough that

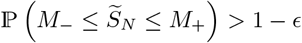

holds with *M*_−_ = (1 − *δ*)*Nm*_*N*_ and *M*_+_ = (1 + *δ*)*Nm*_*N*_ from (55) and 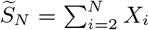 from (23b) in Definition 3.2. For any fixed 0 *< x <* 1

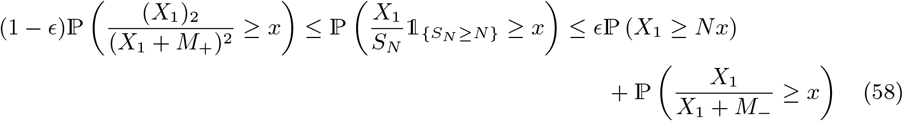

Recall event *E* (*X*_1_, …, *X*_*N*_ ▷ *L*(*α, ζ*(*N*))) from Definition 3.3 (type *A* random environment) and that by Lemma 5.4 and Lemma 5.7 we can choose (*ε*_*N*_) as in (53) so that *m*_∞_ *<* ∞. Using (21) we see (recall *m*_*N*_ = E [*X*_1_] as in (23c))

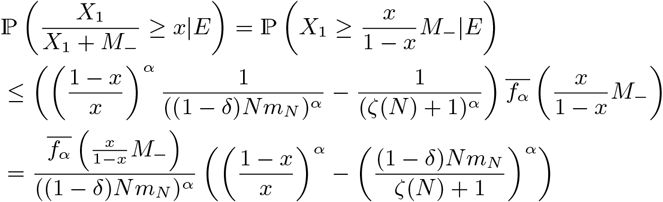

For 0 *< α <* 2, *w* ≥ 0, and 0 *< x* ≤ 1*/*(1 + *w*) we have [Chetwynd-Diggle and Eldon, 2026, Equation 10.2]

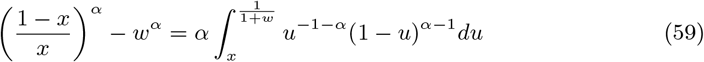

using the substitution *y* = (1 − *u*)*/u*. Write 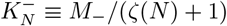 so that

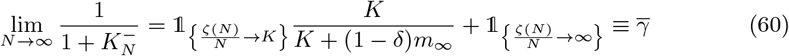

By Lemma 5.7, recalling *ε*_*N*_ from (53), with 1 ≤ *α <* 2,

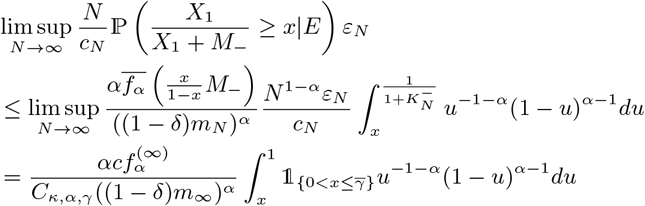

using that 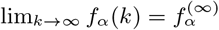 (recall (20a)), and *C*_*κ,α,γ*_ is as in (54) in Lemma 5.7.

We consider the lower bound in (58). Write 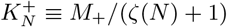 so that

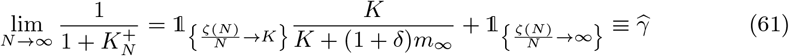

We see using (21) and that 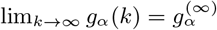 (recall (20a)) and with 1 ≤ *α <* 2

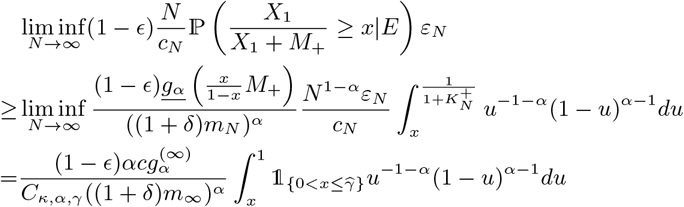

The lemma follows after taking *ϵ* and *δ* to zero and recalling that 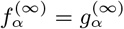 by assumption.

The proof of Lemma 5.8 is complete.

The following lemma shows that our choice (53) of *ε*_*N*_ guarantees that simultaneous mergers vanish in the limit when 1 ≤ *α <* 2.

#### Lemma 5.9

(Simultaneous mergers vanish). *Condition* (6b) *holds under the conditions of Case 2 of Theorem 3.5*.

*Proof of Lemma 5.9*. We see, following the proof of [Schweinsberg, 2003, Lemma 15], recalling 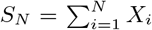 as in (23a), *c*_*N*_ from Definition 2.2, *ε*_*N*_ from Definition 3.3 (type *A* random environment)

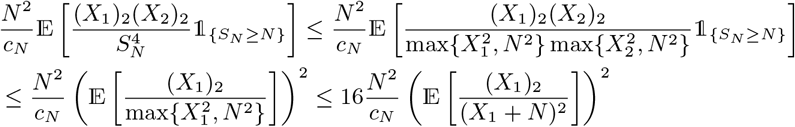

When *α* = 1 we use Case 2 of Lemma 5.2 to see that 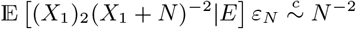 and Case 4 of the same lemma to see that 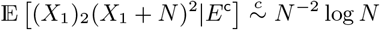 when *κ* = 2 as *N* → ∞ and with *ε*_*N*_ as in (53); hence lim sup_*N* →∞_ (*N*^2^/c_N_) (E [(*X*_1_)_2_(*X*_1_ + *N*)^−2^])_^2^_ = 0 using (28); the case for 1 *< α <* 2 follows similarly, hence the lemma using (16).

#### Lemma 5.10

(Convergence to the Kingman coalescent). *Suppose the conditions of Case 1 of Theorem 3.5 hold. Then* 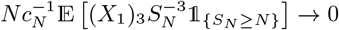.

*Proof of Lemma 5.10*. On *S*_*N*_ ≥ *N* we have 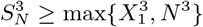 so that with *ζ*(*N*) *< N* we have

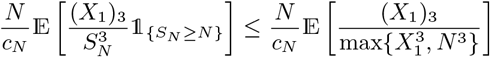

When *α >* 1 (51) gives

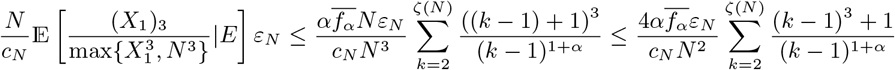

Combining (53) and Lemma 5.7 we see with 1 *< α <* 2

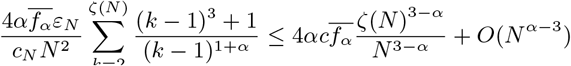

When *α* = 1 we have, by the upper bound in (50) and, since *ζ*(*N*)*/N* → 0 by assumption,

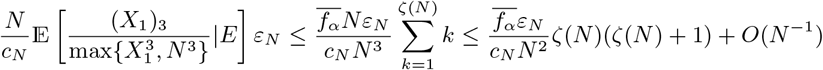

The lemma follows from Lemma 5.7 with *ε*_*N*_ as in (53) so that lim sup_*N*→∞_ *ε*_*N*_ */c*_*N*_ *<* ∞.

#### Remark 5.11

(Convergence to the Kingman coalescent when 0 *< α <* 1). *Suppose the conditions of Theorem 3.5 hold with ζ*(*N*)*/N* → 0 *and κ >* 2 *and* 0 *< α <* 1. *Then (assuming Nc*_*N*_ ~ *c and Nε*_*N*_ ~ *c*^′^ *for c, c*^′^ *>* 0 *fixed) similar calculations as in the proof of Lemma 5.10 give that*

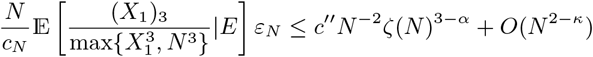

*for some fixed c*^′′^ *>* 0. *Hence, by [Schweinsberg, 2003, Proposition 4] convergence to the Kingman coalescent only holds if ζ*(*N*)^3−*α*^*/N* ^2^ → 0 *when* 0 *< α <* 1 *under our assumptions on c*_*N*_ *and ε*_*N*_. *Taking ζ*(*N*) = *N* ^*x*^ *shows that convergence to the Kingman coalescent follows if x <* 2*/*(3 − *α*) *<* 1, *compare with Case 1 of Theorem 3.5*.

*Proof of parts 1 and 2 of Theorem 3.5*. Part 1 of Theorem 3.5 is Lemma 5.10 when 1 ≤ *α <* 2 using (16) and [Schweinsberg, 2003, Proposition 2] (recall (7)).

We turn to part 2 of Theorem 3.5. We need to check the three conditions of [Schweinsberg, 2003, Proposition 3]; these are (6a), (6b), and (6c) in Proposition 2.3. Lemma 5.7 gives (6a); choosing *ε*_*N*_ as in (53) gives Lemma 5.4 so that Lemma 5.6 holds and with that Lemma 5.7. Condition (6b) is Lemma 5.9. We turn to (6c). We need to show that (recall *γ* from (26a), and *ν*_1_ is the random number of surviving offspring of (arbitrary) individual 1)

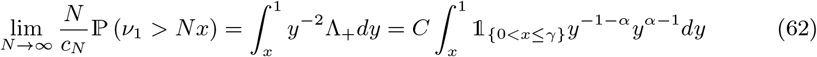

for 0 *< x <* 1. By the arguments of the proof of part (c) of Theorem 4 of [Schweinsberg, 2003] using bounds on the tail probability of a hypergeometric [Chvátal, 1979] we have

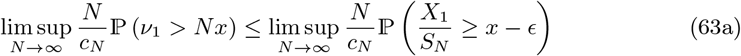

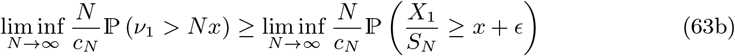

for 0 *< ϵ* ≤ *x*. Lemma 5.8 together with (63a) and (63b) and taking *ϵ* to 0 gives (62).

*Proof of part 3 of Theorem 3.5*. Finally, we check part 3 of Theorem 3.5, the case 0 *< α <* 1 and *ζ*(*N*)*/N* ^1*/α*^ → ∞.

Following [Schweinsberg, 2003, § 4] let *Y*_(1)_ ≥ *Y*_(2)_ ≥ … ≥ *Y*_(*N*)_ be the ranked values of *N* ^−1*/α*^*X*_1_, …, *N* ^−1*/α*^*X*_*N*_. With *p*_*i*_ = ℙ(*X*_1_ ≥ *N* ^1*/α*^*x*_*i*_|*E)* − ℙ (*X*_1_ ≥ *N* ^1*/α*^*x*_*i*−1_|*E)* using (21) we see, recalling *E* is the event *X*_1_, …, *X*_*N*_ ▷ L(*α, ζ*(*N*)) from Definition 3.3 (type *A* random environment), and (*x*_*j*_)_*j*∈ℕ_ is a decreasing sequence of positive numbers with *x*_0_ = ∞,

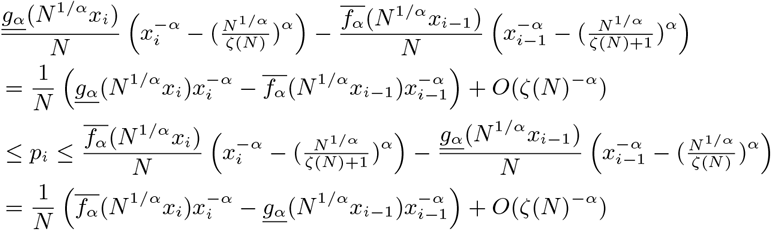

for 1 ≤ *i* ≤ *j* so that *p* = 1 − *p*_1_ −… −*p*_*j*_ = 1 − ℙ (*Y*_1_ ≥ *x*_*j* |_*E*). Moreover, for non-negative integers *n*_1_, …, *n*_*j*_ we have

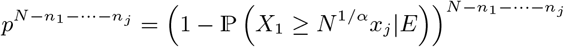

Using (21) again we have

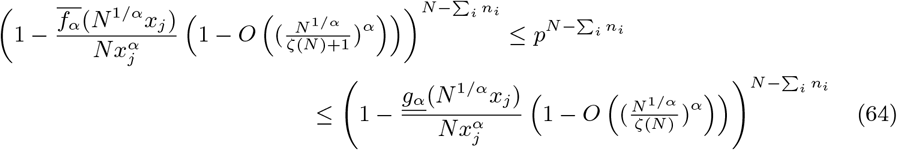

so that

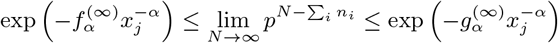

Then

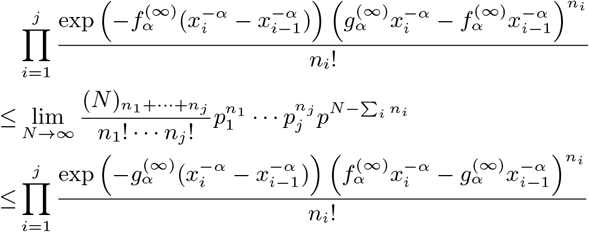

Choosing *g*_*α*_ and *f*_*α*_ in (18) such that 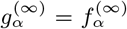 we have 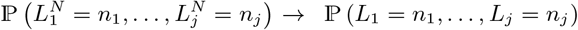 where 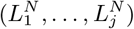 and (*L*_1_, …, *L*_*j*_) are as in the proof of [Schweinsberg, 2003, Lemma 20].

As briefly mentioned in § 2 Poisson-Dirichlet distributions can be constructed from the ranked points *Z*_1_ ≥ *Z*_2_ ≥ … of a Poisson point process with characteristic measure *ν*_*α*_((*x*, ∞)) = *1*_{*x>*0}_*Cx*^−*α*^. The law of (*Z*_*j*_*/* ∑*Z*_*i*_)_*j*∈ℕ_ is the Poisson-Dirichlet distribution with parameters (*α*, 0).

The upper bound in (50) gives (recalling *ζ*(*N*)*/N* ^1*/α*^ → ∞)

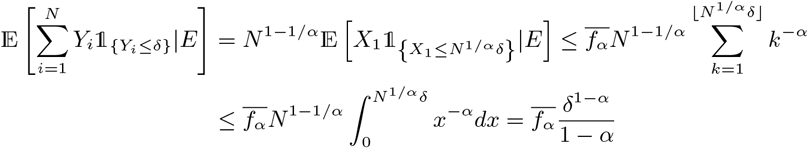

On *E* we then have that 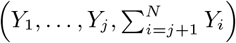 converges weakly to 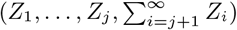 as *N* → ∞ by the arguments for [Schweinsberg, 2003, Lemma 21]. With (*W*_*j*_)_*j*∈ℕ_ = (*Z*_*j*_*/* ∑*Z*_*i*_)_*j*∈ℕ_ it follows that, provided 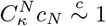 and with 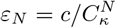 for some *c >* 0,

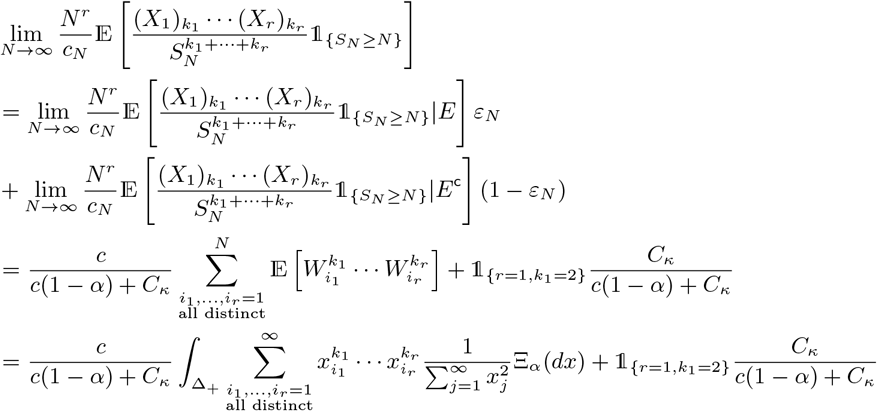

for all *k*_1_, …, *k*_*r*_ ≥ 2 and *r* ∈ ℕ where Ξ_*α*_ is as in Definition 2.6 with *C*_*κ*_ as in (26c) and 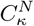 as in (24).

It remains to verify that 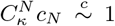. We have 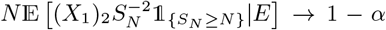 [Schweinsberg, 2003, pp. 137]. By Lemma 5.7 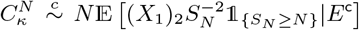. Then 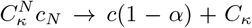. Part 3 of Theorem 3.5 now follows from our choice of *ε*_*N*_ and [Schweinsberg, 2003, Möhle and Sagitov, 2001, Proposition 1; Theorem 2.1] where

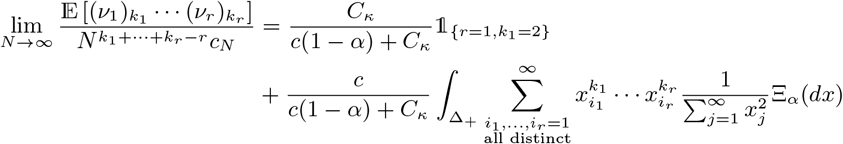

The proof of Theorem 3.5 is complete.

#### Remark 5.12

(The case 0 *< α <* 1 and *ζ*(*N*)*/N* ^1*/α*^ → *K*). *Suppose, under the conditions of Theorem 3.5, we have* 0 *< α <* 1 *and ζ*(*N*)*/N* ^1*/α*^ → *K (K >* 0 *fixed)*.

*Let K* = *x*_0_ ≥ *x*_1_ ≥ *x*_2_ ≥ … ≥ *x*_*j*_ *>* 0 *all be fixed. Then, with* 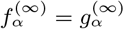 *and recalling event E from Definition 3.3 (type A random environment)*,

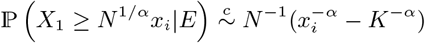

*Taking p* = ℙ (*X* ≥ *N* ^1*/α*^*x* |*E)* − ℙ (*X*_1_ ≥ *N* ^1*/α*^*x*_*i*−1_ |*E) for all* 1 ≤ *i* ≤ *j we have* 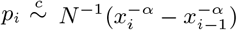. *Moreover, taking p* = 1 − *p*_1_ − · · · − *p*_*j*_ *we see*

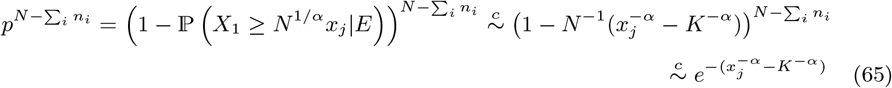

*for n*_1_, …, *n*_*j*_ ∈ ℕ ∪ {0}. *Then*

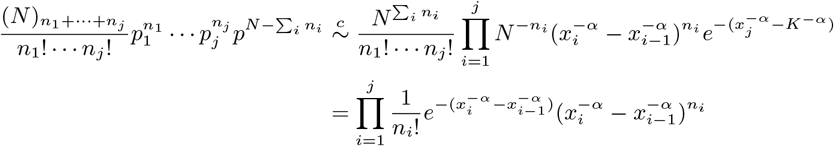

*We conjecture that the last expression corresponds to the probability* 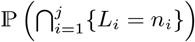 *where L*_*i*_ *is the number of points of a Poisson point process* {*N*_*α,K*_} *between x*_*i*_ *and x*_*i*−1_ *where* 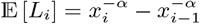 *with x*_0_ = *K and* {*N*_*α,K*_} *has intensity measure ν*_*α,K*_ ((*x*, ∞)) = *1*_{0*<x*≤*K*}_(*x*^−*α*^ − *K*^−*α*^). *Suppose Z*_1_ ≥ *Z*_2_ ≥ … *are the ranked points of* {*N*_*α,K*_}. *Then, in contrast to the case when ζ*(*N*)*/N* ^1*/α*^ → ∞, *it is not clear what the law of* (*Z*_*j*_ */* ∑_*i*_*Z*_*i*_)_*j*∈ℕ_ *converges to*.

### 5.3 Proof of Theorem 3.6

In this section we give a proof of Theorem 3.6 (recall the type *B* random environment from Definition 3.4). To briefly orient the reader through the proof of Theorem 3.6 (the proofs can always be skipped on first reading to get an overview of the lemmas leading to the theorem), we use the same approach as for Theorem 3.5, namely checking the conditions of Proposition 2.3 for Case 2 of Theorem 3.6, and Condition (7) for Case 1 of the theorem. Recall from Definition 3.4 that 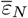 is the probability of event *E*_1_ (that one randomly picked *X*_*i*_ ▷ L(*α, ζ*(*N*)), and all other *X*_*j*_ ▷ L(*κ, ζ*(*N*))), Lemma 5.13 identifies conditions on 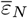 such that *m*_∞_ = lim_*N* →∞_ E [*X*] *<* ∞, and in Lemma 5.14 further conditions on 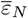 such that *S*_*N*_ */*(*Nm*_*N*_) converges almost surely to 1, in turn useful for deriving the limits of 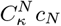 in Lemma 5.16. Lemma 5.18 identifies the Λ_+_-measure (recall Condition (6c) in Proposition 2.3), Lemma 5.19 checks that simultaneous mergers vanish in the limit (recall Condition 6b of Proposition 2.3), and Lemma 5.21 verifies Case 1 of Theorem 3.6 by checking Condition (7).

First we identify conditions for *m*_∞_ = lim_*N* →∞_ E [*X*] from (23c) to be finite, with *X* denoting the random number of potential offspring of an arbitrary individual.

#### Lemma 5.13

(Finite *m*_∞_). *Under the conditions of Theorem 3.6, with ζ*(*N*)*/N* ≩ 0 *and* 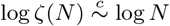. *Recalling* 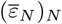 *from Definition 3.4 (* 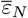 *is the probability of event E*_1_ *in type B random environment) suppose*

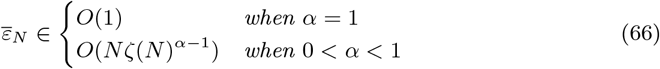

*Then we have* lim sup_*N* →∞_ E [*X*] *<* ∞.

*Proof of Lemma 5.13*. We see, recalling the events *E*_1_ and 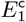 from Definition 3.4 (type *B* random environment; event *E*_1_ is when *X*_*i*_ ▷ L(*α, ζ*(*N*)), and *X*_*j*_ ▷ L(*κ, ζ*(*N*)) for all *j* ∈ [*N*] \ {*i*} with index *i* picked uniformly at random; event 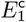 is when *X*_1_, …, *X*_*N*_ ▷ L(*κ, ζ*(*N*))),

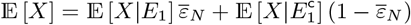

When *α* = 1 and *κ* ≥ 2 using the upper bound in (51) in Lemma 5.3

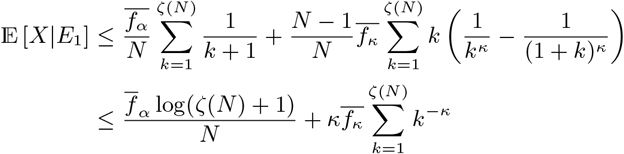

When *α* ∈ (0, 1), we have E [*X*|*E*_1_] ≤ *cζ*(*N*)^1−*α*^*/N* by (52) and so the lemma by (66).

We verify a convergence result on *S*_*N*_ */*(*Nm*_*N*_) similar to Lemma 5.6; the result will be useful for deriving limits of 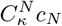.

#### Lemma 5.14

(*S*_*N*_ */*(*Nm*_*N*_) → 1 almost surely). *Suppose the conditions of Theorem 3.6 hold with m*_*N*_ = E [*X*_1_] *and α* ∈ (0, 1]. *Let* 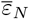 *from Definition 3.4 (type B random environment) fulfill the condition from Lemma 5.13 and, with* 1 *< r <* 2 *fixed*,

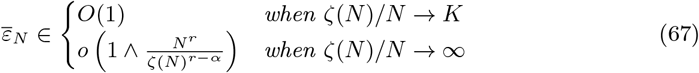

*Then S*_*N*_ */*(*Nm*_*N*_) *converges almost surely to 1 as N* → ∞ *with* 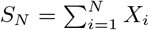 *as in* (23a).

*Proof of Lemma 5.14*. We follow the approach of [Panov, 2017]. Our choice of *ε*_*N*_ guarantees that lim sup_*N*→∞_ *m*_*N*_ *<* ∞ by Lemma 5.13. Write 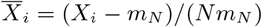, and

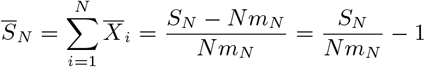

Fix 1 *< r <* 2. Provided 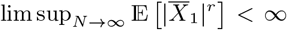 we have by [von Bahr and Esseen, 1965, Theorem 2]

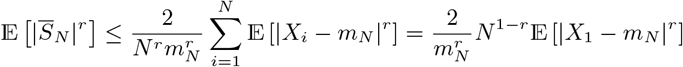

Then, using Lemma 5.3 with *α* ∈ (0, 1], and with *m*_*N*_ *<* ∞ for all *N* by Lemma 5.13

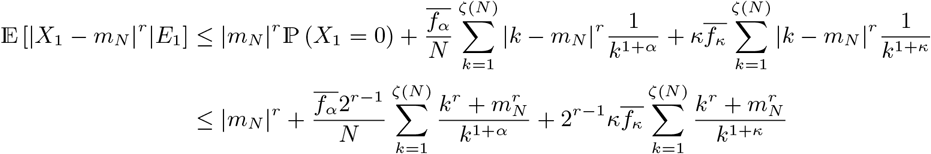

where we have used that |*x* + *y*| ^*r*^ ≤ 2^*r*−1^(|*x*|^*r*^ + |*y*|^*r*^) for any *x, y* real and *r* ≥ 1 [Athreya and Lahiri, 2006, Equation (1.12), Proposition 3.1.10*(iii)* pp. 87] (follows easily from Jensen’s inequality since ℝ ∋ *x* ↦→ |*x*|^*r*^ is convex for *r* ≥ 1). Recalling 0 *< α* ≤ 1 *< r <* 2 ≤ *κ* we see

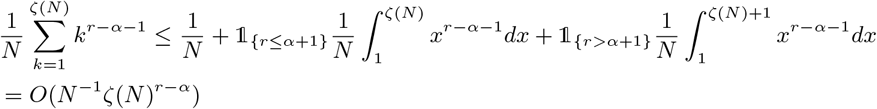

Then E [|*X*_1_ − *m*_*N*_ |^*r*^|*E*_1_] ∈ *O*(1 + *N* ^−1^*ζ*(*N*)^*r*−*α*^). With 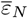 as in (67) we then have

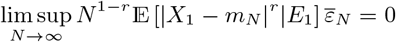

for any 1 *< r <* 2. By similar calculations 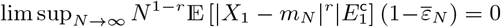 there is a *r* > 1 such that 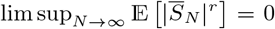. Hence 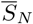 converges in probability to 0, and since the 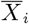 are independent, 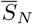 converges almost surely to 0 (see e.g. [Athreya and Lahiri, 2006, Theorem 8.3.3 pp 251]). Hence, 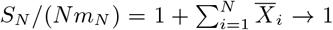 almost surely.

#### Remark 5.15

(Convergence of *S*_*N*_ */*(*Nm*_*N*_)). *Using the upper bounds in* (50) *and* (51) *in Lemma 5.3 it is straightforward to check that* 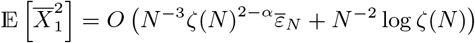.

*Then* 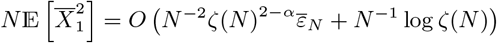. *Taking*

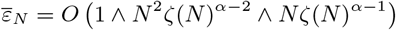

*then gives* lim sup_*N* →∞_ *m_N_ <* ∞ *by Lemma 5.13 and* 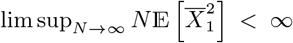. *By Lemma 5.14* 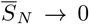 *in probability*, 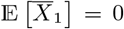, *so by Khinchine-Kolmogorov’s 1-series theorem* 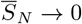 *almost surely. Then* 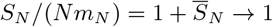 *almost surely*.

We verify the scaling of *c*_*N*_ (recall Definition 2.2).

#### Lemma 5.16

(Scaling of *c*_*N*_). *Suppose the conditions of Theorem 3.6 hold with* 0 *< α* ≤ 1, *κ* ≥ 2, *c >* 0 *and* 0 *< ε <* 1 *all fixed. Recall* 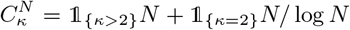 *as in* (24). *Let* 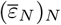 *from Definition 3.4 (type B random environment;* 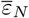 *is the probability of event E*_1_ *that a randomly picked X*_*i*_ ▷ L(*α, ζ*(*N*) *and all other X*_*j*_ ▷ L(*κ, ζ*(*N*))*) take the values*

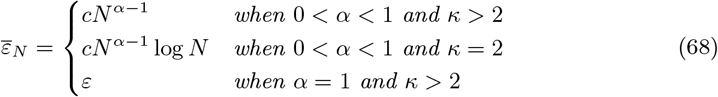

*and suppose* 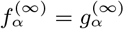. *Then* 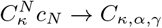 *with* 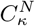 *as in* (24) *and C as in* (54) *except c*_*α*_ = *1*_{0*<α<*1}_*c* + *1*_{*α*=1}_*ε replaces c*.

#### Remark 5.17

(Condition on 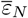 in Lemma 5.16). *When ζ*(*N*)*/N* → *K Lemma 5.14 imposes no new restrictions on* 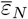, *as well as when* 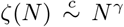 *with* 1 *< γ < r/*(*r* − *α*); *since r/*(*r* − *α*) ≤ *r* + 1 − *α so that N* ^*α*−1^ = *O*(*N* ^*r*^*ζ*(*N*)^*α*−*r*^), *and* lim sup_*N*→∞_ *m*_*N*_ *<* ∞ *by Lemma 5.13. Further*, _*N*_ ^*α*−1^ = *O*(*N* ^1−*γ*(1−*α*)^_)_ *for any γ >* 1. *Thus, a mild condition on ζ*(*N*) *when ζ*(*N*)*/N* → ∞ *allows to choose* 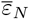 *as in* (68) *such that S*_*N*_ */*(*Nm*_*N*_) → 1 *almost surely by Lemma 5.14 and* lim sup_*N*→∞_ *m*_*N*_ *<* ∞ *by Lemma 5.13*.

*Proof of Lemma 5.16*. Recall *Y*_−_ and *Y*_+_ from (55). Similar to the proof of Lemma 5.7 we use Lemma 5.14 to get (56). It then suffices to consider lower bounds on 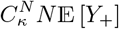 and upper bounds on 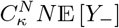 and to check that (57) holds. These all follow as in the proof of Lemma 5.7, noting that by Definition 3.4, writing *p*_*k*_(*a*) ≡ *k*^−*a*^ − (1 + *k*)^−*a*^ for *k* ∈ ℕ and *a >* 0,

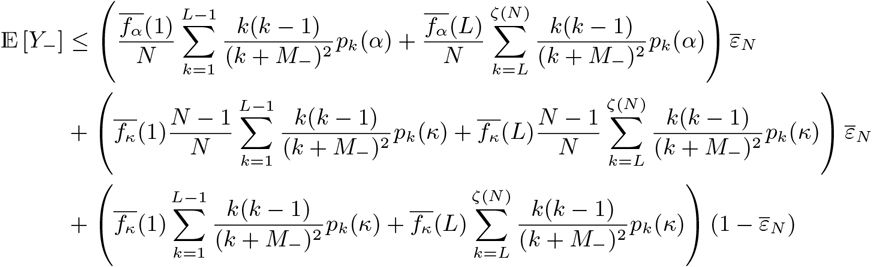

A lower bound on E [*Y*_+_] is of a similar form with 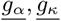 replacing 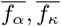, and the lemma follows as in the proof of Lemma 5.7 upon substituting for *ℓ* and *m* in Lemma 5.2 and taking limits.

#### Lemma 5.18

(Identifying the Λ_+_-measure). *Under the conditions of Theorem 3.6 for all* 0 *< x <* 1 *with m*_∞_ = lim_*N* →∞_ E [*X*_1_] *as in* (23c) *and* 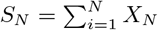 *as in* (23a) *we have*

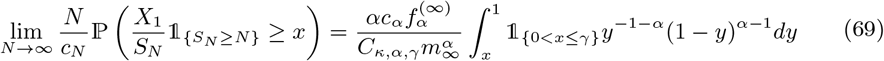

*with γ* = *1*_{*ζ*(*N*)*/N*→∞}_ + *1*_{*ζ*(*N*)*/N*→*K*}_*K/*(*m*_∞_ + *K*) *as in* (26a), *C*_*κ,α,γ*_ *as in* (26b) *with c*_*α*_ = *1* _0*<α<*1_ *c* + *1 ε replacing c*.

*Proof of Lemma 5.18*. The proof follows the one of Lemma 5.8. When *α* = 1 and *κ >* 2 with 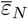 as in (68) so that *m*_∞_ *<* ∞ by Lemma 5.13 (recall *M*_+_ = (1 + *δ*)*Nm*_*N*_ from (55))

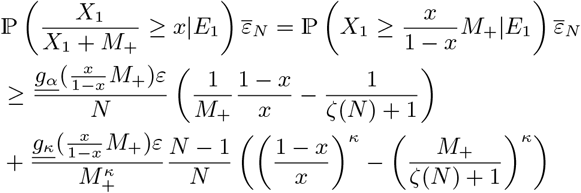

Using Lemma 5.16 (recall *γ* from (26a)) and since (with *α* = 1)

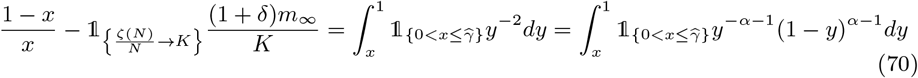

with 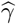 as in (61) we get (after checking that 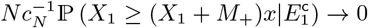)

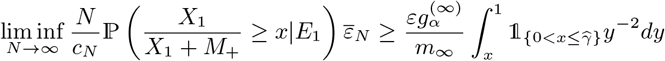

after taking *δ* to 0. For the upper bound we see (recall *M*_−_ = (1 − *δ*)*Nm*_*N*_ from (55), and *α* = −1)

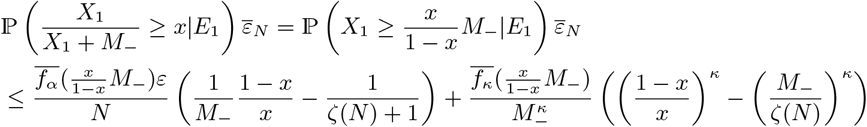

Analogously to the lower bound we get (when *α* = 1 and *κ >* 2) with 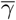 from (60)

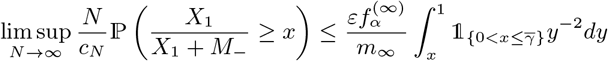

The result when *α* = 1 follows after taking *δ* to 0. The case 0 *< α <* 1 proceeds in the same way where we make use of (59).

#### Lemma 5.19

(Simultaneous mergers vanish). *Condition* (6b) *follows from the settings of Theorem 3.6*.

*Proof of Lemma 5.19*. The proof follows the arguments for Lemma 5.9. Choosing 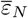 as in (68) then by Case 1 of Lemma 5.2 we have 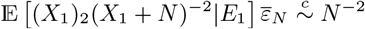 as *N* → ∞ so that by Lemma 5.16 lim sup_*N* →∞_ *N* ^2^(E [(*X*_1_)_2_(*X*_1_ + *N*)])^2^*/c*_*N*_ = 0. Hence to (6b) by the arguments for Lemma 5.9.

#### Remark 5.20

(Condition (6b) follows from *m*_∞_ *<* ∞). *Since m*_∞_ *<* ∞ *under Theorem 3.6 (by Lemma 5.13 choosing* 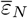 *as in* (68)*) and Case 2 of Theorem 3.5 (by Lemma 5.4 choosing ε*_*N*_ *as in* (53)*) condition* (6b) *follows from Lemma 15 of [Schweinsberg, 2003]*.

#### Lemma 5.21

(Convergence to the Kingman coalescent). *When ζ*(*N*)*/N* → 0 *Case 1 of Theorem 3.6 follows*.

*Proof of Lemma 5.21*. By [Schweinsberg, 2003, Proposition 4] (see also [Möhle, 2000, § 4]) it suffices to check that (7) holds. By (16) in Theorem 2.9 it is sufficient to show that 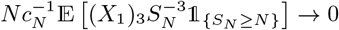. On *S*_*N*_ ≥ *N* so that 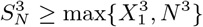.

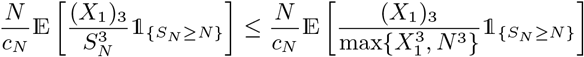

We see, for 0 *< α* ≤ 1 and *κ >* 2 using the upper bound in Eq (50) in Lemma 5.3 and (68) for some *C*^′′^ *>* 0 fixed

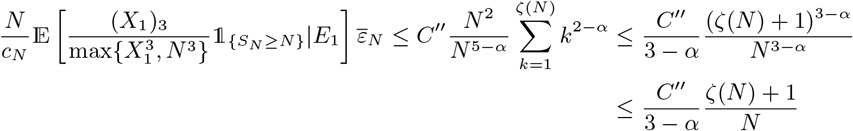

for all *N* ≥ *ζ*(*N*) + 1 (recall *ζ*(*N*)*/N* → 0 by assumption). Hence

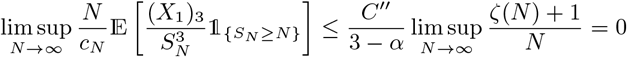

and the lemma follows using (16).

#### Remark 5.22

(The condition 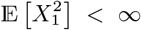). *In the same manner as in the proof of Lemma 5.21 one checks that* 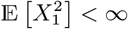 *when κ >* 2 *and ζ*(*N*)*/N* → 0; *then by [Schweinsberg, 2003, Proposition 7] Case 1 of Theorem 3.6 holds*.

*Proof of Case 1 of Theorem 3.6*. Case 1 of Theorem 3.6 is Lemma 5.21.

*Proof of Case 2 of Theorem 3.6*. Condition 6a is Lemma 5.16; condition 6b is Lemma 5.19; 6c follows from Lemma 5.18 in the same way as for Case 2 of Theorem 3.5. Case 2 of Theorem 3.6 then follows by Proposition 2.3.

### 5.4 Proof of Theorem 3.8

*Proof of Theorem 3.8*. Theorem 3.8 follows from Theorem 3 of [Freund, 2020]. Write *X*_*i*_(*r*) for the number of potential offspring of individual *i* (arbitrarily labelled) for all *i* ∈ [*N*_*r*_]. Suppose *ζ*(*N*_*r*_)*/N* ≩ 0 for all *r*, and that either *ζ*(*N*_*r*_)*/N*→ ∞ for all *r*, or *ζ*(*N*_*r*_)*/N* → *K* for all *r*. Choosing *f* and *g* in (18) such that E [*X*_*i*_(*r*)] *>* 1 for all *i* ∈ [*N*_*r*_] and *r* ∈ ℕ_0_ gives [Freund, 2020, Lemma 1],

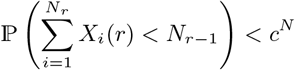

for some constant 0 *< c <* 1. By Theorems 3.5 and 3.6 (recall (28)) it holds that 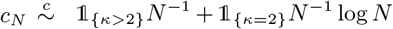. Then, by [Freund, 2020, Lemma 4], {*ξ*^*n,N*^ (⌊*t/c*_*N*_ ⌋); *t* ≥ 0} converges weakly to {*ξ*^*n*^ (*G*(*t*)); *t* ≥ 0}, and 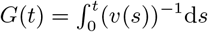.

Define 1_*A*_(*s*) ≡ 1 when *s* ∈ *A* for some set *A*, take 1_*A*_(*s*) ≡ 0 otherwise. We need to check that, for the case *κ* = 2, it holds that

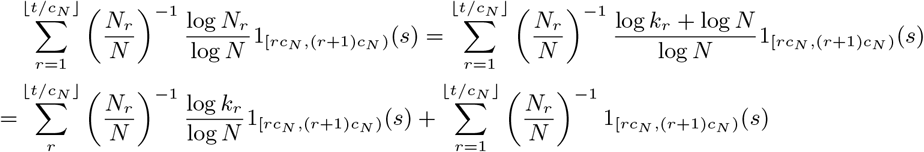

Using (41) it follows that, as *N* → ∞,

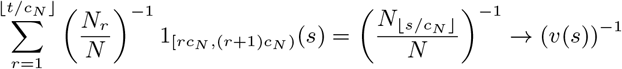

Recalling (41) again take *h*_1_(*t*) ≤ *k*_*r*_ ≤ *h*_2_(*t*) with *h*_2_ bounded. It remains to check that

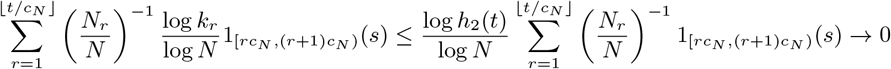

as *N* → ∞

#### Remark 5.23

(Exponential growth gives a well-defined population sequence). *When N*_*r*+1_ = ⌊*N*_*r*_ (1 + *ρc*_*N*_)⌋ *as in exponential growth it holds*

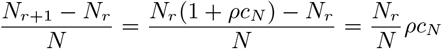

*Since* 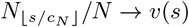 *and c*_*N*_ → 0 *it holds* ((*N*_*r*+1_ − *N*_*r*_) */N*)_*r* ∈ℕ_ *is a null-sequence such that [Freund, 2020, Lemma 1] holds for any fixed ρ >* 0.

### 5.5 Proof of Theorem 3.9

In this section we give a proof of Theorem 3.9. A version of Lemma 2 of [Freund, 2020, Lemma 2] holds for Ξ-coalescents (when started from a finite number *n* of blocks; Ξ-*n*-coalescents) provided *(i)* we have a sequence 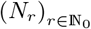 of population sizes where *N*_*r*_ is the population size *r* generations into the past such that (41) holds; *(ii)* since 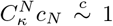 it holds that *c*_*N*_ is regularly varying so that [Freund, 2020, Equation 13]

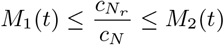

for all *r* ≤ ⌊*t/c*_*N*_ ⌋ for some bounded positive functions *M*_1_, *M*_2_ : [0, ∞) → (0, ∞); *(iii) c*_*N*_ → 0 as *N* → ∞ by (28); *(iv)* the condition given in [Freund, 2020, Equation 5] follows as in the proof of [Freund, 2020, Theorem 3]; *(v) {ξ*^*n,N*^ (⌊*t/c*_*N*_ ⌋); *t* ≥ 0} converges to a continuous-time Ξ-coalescent {*ξ*^*n*^} as *N* → ∞ by Case 3 of Theorem 3.5. Then, we have by [Freund, 2020, Lemma 2] that 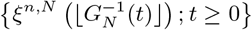 converges weakly to {*ξ*^*n*^} as *N* → ∞, where *G*_*N*_ is as in [Freund, 2020, Equation 1].

Since (41) and (28) hold, [Freund, 2020, Lemma 4] establishes that the convergence of 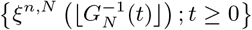; to {*ξ*^*n*^*}* is equivalent to the convergence of {*ξ*^*n,N*^ (*t/c*); *t* 0} to {*ξ*^*n*^ (*G*(*t*)); *t* ≥ 0}, where 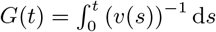.

## Declarations

### Ethical Statement

The author has no conflict of interest to declare that are relevant to the content of this article

## Data availability

The documented software (C/C++ code) developed for the numerical results is freely available at https://github.com/eldonb/gene_genealogies_haploid_populations_sweepstakes.

## Acknowledgements

The author would like to thank two anonymous reviewers for useful comments.

## APPENDIX A The Beta(2−*β, β*)-Poisson-Dirichlet(*α*, 0)-co

In this section we briefly study the Beta(2 − *α, α*)-Poisson-Dirichlet(*α*, 0)-coalescent. Fix 0 *< α <* 1 and 1 *< β <* 2. Consider a population evolving as in Definition 3.3 with *β* replacing *κ*, and such that *ζ*(*N*)*/N* ^1*/α*^ → ∞ as *N* → ∞. The calculations leading to Cases 2 and 3 of Theorem 3.5, assuming 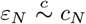 where 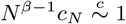 as *N* → ∞ lead to a coalescent with transition rates

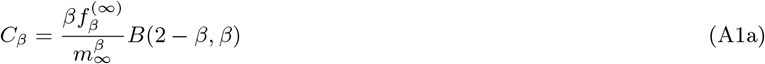

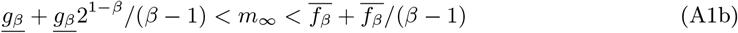

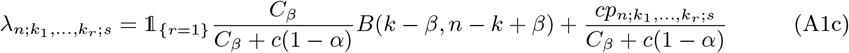

where *p*_*n*;*k*,…,*k*;*s*_ is as in (12), 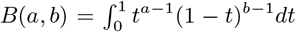 for *a, b >* 0, and *c >* 0 fixed (we take *ε*_*N*_ = *cN* ^1−*β*^). We will refer to the coalescent with transition rates as in (A1c) as the Beta(2 − *β, β*)-Poisson-Dirichlet(*α*, 0)-coalescent. It is a multiple-merger coalescent (a Ξ-coalescent) without an atom at zero. Moreover, one coalescent time unit is proportional to *N* ^*β*−1^ generations, in contrast to the time scaling in Theorems 3.5 and 3.6 (recall (28)). Figure A1 holds examples of 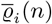 (31) predicted by the Beta(2 − *β, β*)-Poisson-Dirichlet(*α*, 0)-coalescent. The algorithm for sampling the coalescent is a straightforward adaption of the one for sampling the *δ*_0_-Poisson-Dirichlet(*α*, 0)-coalescent described in § 3.2.3.

## Appendix B Examples of *φ*_*i*_(*n*) for the *δ*_0_-Beta(*γ*, 2 − *α, α*)-coalescent

In this section we give in Figure B1 examples of *φ*_*i*_(*n*) as in (32), § 3.2, and predicted by the *δ*_0_-Beta(*γ*, 2 − *α, α*)-coalescent. In Figures B1a–B1c *γ* is fixed and we vary over *α* as shown, and in Figures B1d–B1i *α* is fixed and we vary over *γ* as shown; in each graph the black line represents 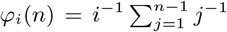 as predicted by the Kingman coalescent [Fu, 1995]. We see when both *α* and *γ* are small enough *φ*_*i*_(*n*) is decreasing (Figures B1a, B1b, B1d) and so will not predict a ‘U-shaped’ site-frequency spectrum seen as a characteristic of Λ-coalescents [Freund et al., 2023]. When *α* ≤ 1 the distribution for the number of potential offspring has a heavy right-tail (recall (18)). In Figure B2 we check that simulated values of *φ*_*i*_(*n*) predicted by the *δ*_0_-Beta(*γ*, 2 − *α, α*)-coalescent match the exact values of *φ*_*i*_(*n*) computed using a recursion for general Λ-coalescents [Birkner et al., 2013b].

**Figure A1.**
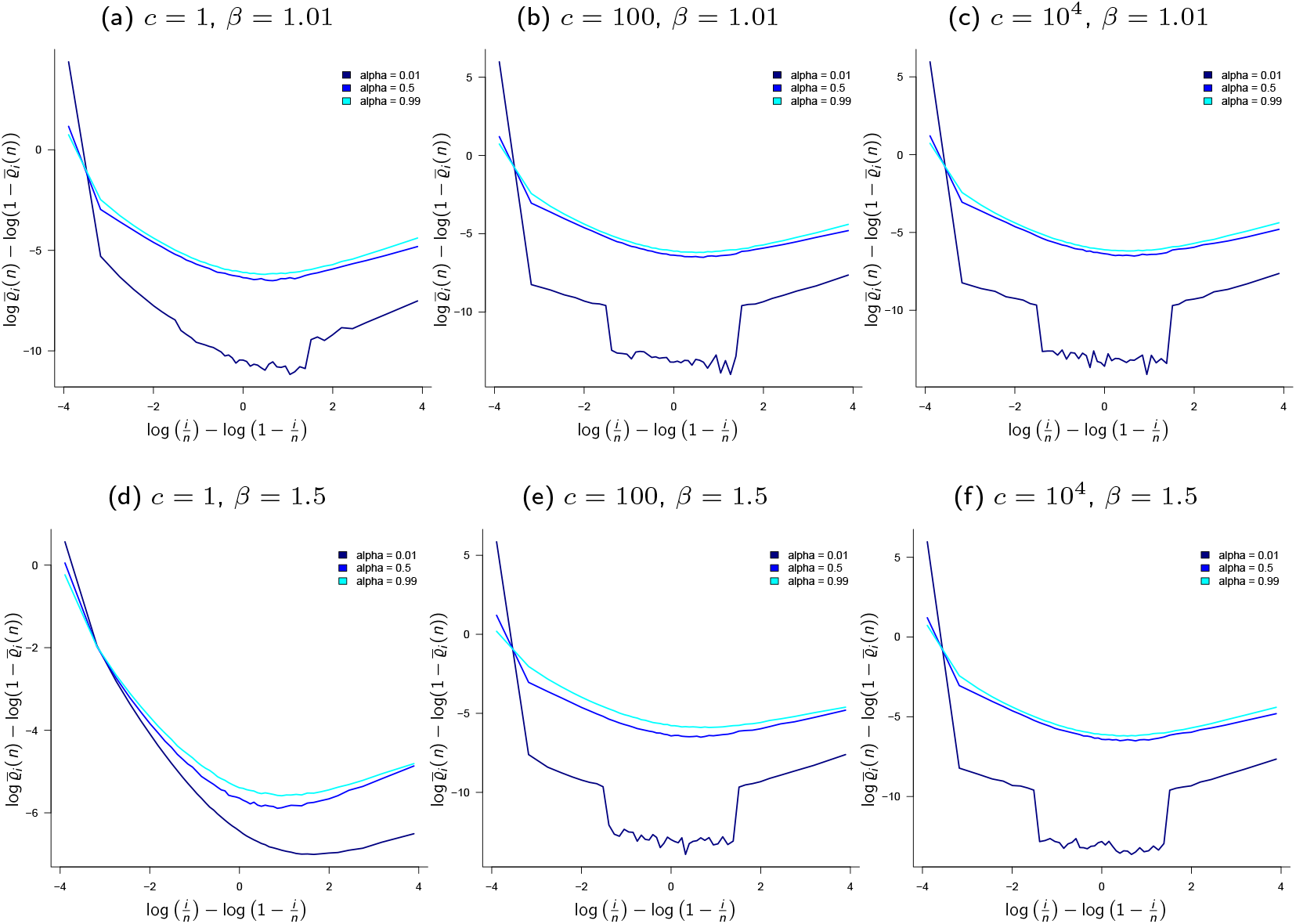
Examples of 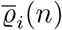 (estimates of mean relative branch lengths, recall (31)) predicted by the Beta(2 − *β, β*)-Poisson-Dirichlet(*α*, 0)-coalescent with *n* = 50, *β, α, c* as shown, and *m*_∞_ (A1b) approximated with (2+ (1 + 2^1+ *β*^) / (*β* − 1))/ 2. The scale of the ordinate (y-axis) may vary between graphs (results from 10^5^ experiments). The scale of the ordinate (vertical axis) may vary between graphs.

**Figure B1.**
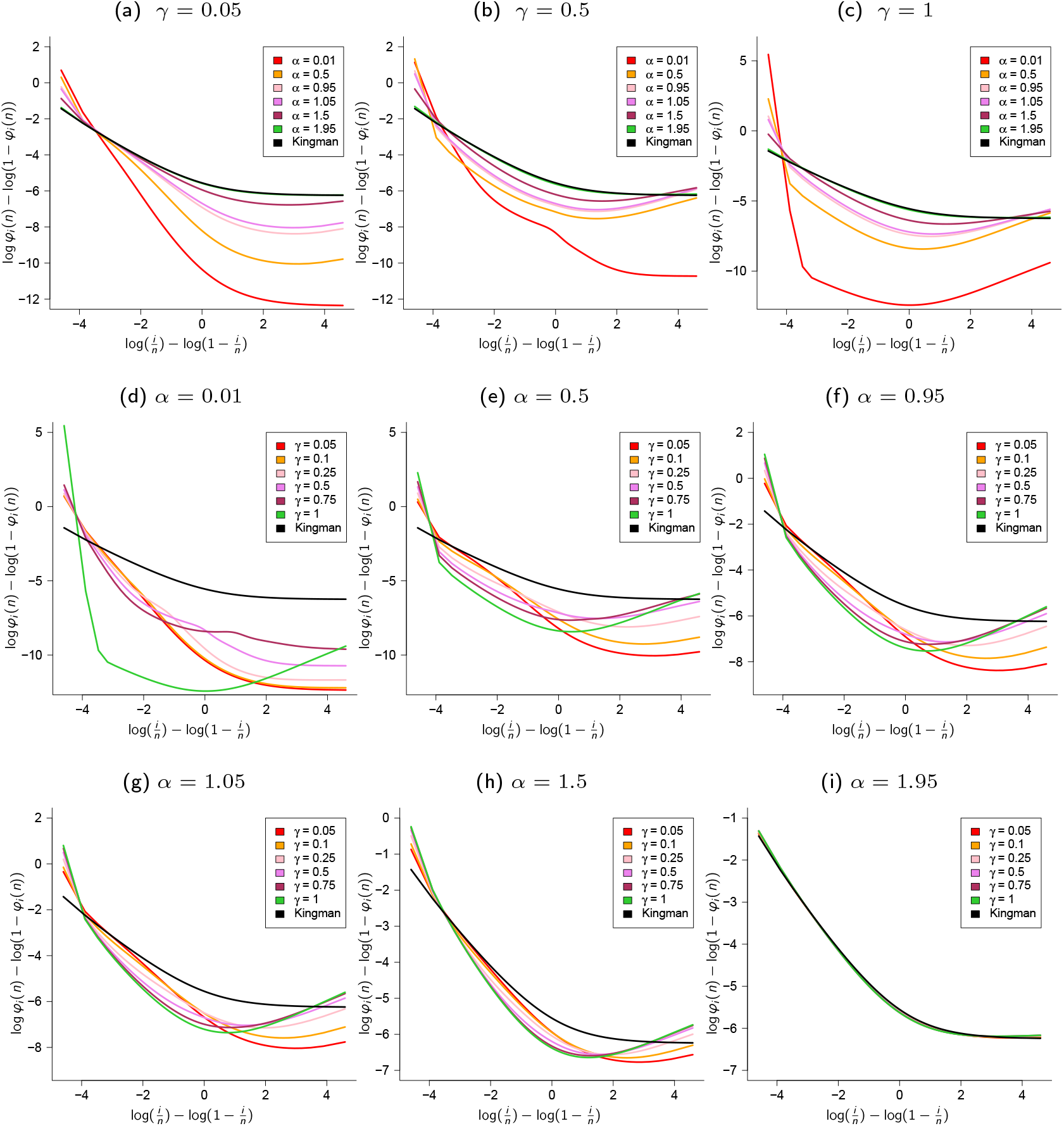
Relative expected branch length *φ*_*i*_(*n*) (32) predicted by the *δ*_0_-Beta(*γ*, 2 − *α, α*)-coalescent with transition rate as in (25) for sample size *n* = 100, *α* and *γ* as shown, and *c* = 1 and *κ* = 2. The expected values computed using a recursion for E [*L*_*i*_(*n*)] derived for Λ-coalescents [Birkner et al., 2013b]; for comparison we show the logits of 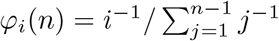 predicted by the Kingman coalescent [Fu, 1995]. The scale of the ordinate (vertical axis) may vary between graphs.

**Figure B2.**
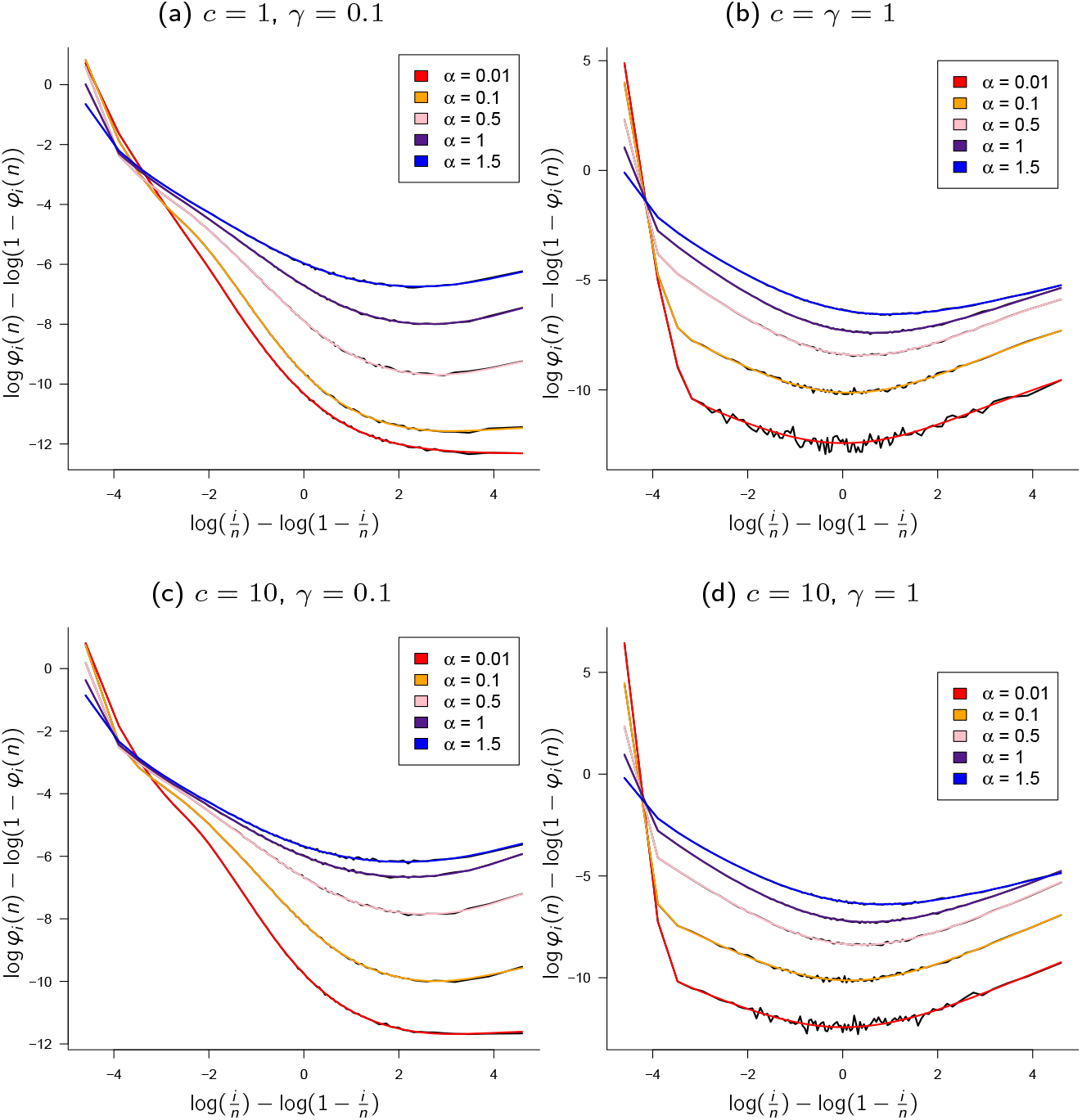
Comparing exact and approximated relative expected branch length *φ*_*i*_(*n*) (32) predicted by the *δ*_0_-Beta(*γ*, 2 − *α, α*) coalescent. Checking that approximations of *φ*_*i*_(*n*) (32) agree with exact values of *φ*_*i*_(*n*) computed using a recursion for general Λ-coalescents [Birkner et al., 2013b]; sample size *n* = 100 with *κ* = 2 and *α* and *c* and *γ* as shown; the coloured lines are exact logit values of *φ*_*i*_(*n*) and the black lines the corresponding estimates for each value of *α*. The approximations were obtained for 10^5^ experiments. The scale of the ordinate (vertical axis) may vary between graphs.

## Appendix C Further graphs comparing 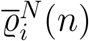 and 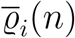

In this section we compare 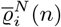 and *ϱ*_*i*_(*n*) (recall (31)) for the *δ*_0_-Beta(*γ*, 2 − *α, α*)-coalescent; in Figure C1 (see also Figure 2) the ancestral process {*ξ*^*n,N*^} is associated with the type *A* random environment (Definition 3.3) such that 1 ≤ *α <* 2, and in Figure C2 (see also Figure 3) the ancestral process is associated with type *B* random environment (Definition 3.4) such that 0 *< α <* 1.

**Figure C1.**
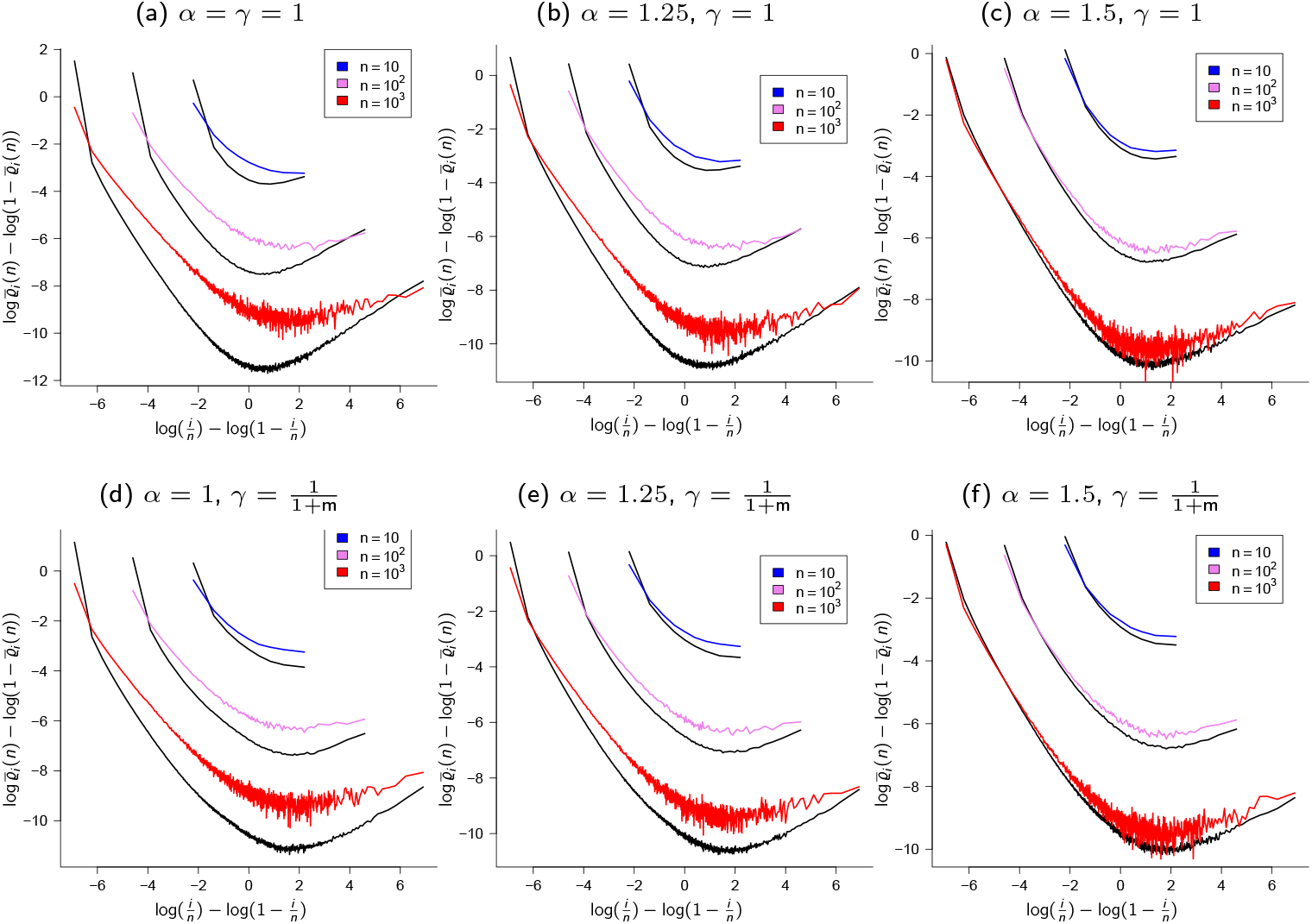
Type *A* random environment (Definition 3.3) and the *δ*_0_-Beta(*γ*, 2 −*α, α*)-coalescent. Comparing 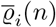 (estimates of mean relative branch lengths predicted by the *δ*_0_-Beta(*γ*, 2−*α, α*)-coalescent, black lines) and 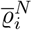 (estimates of mean relative branch lengths predicted by {*ξ*^*n,N*^} when in attraction of the *δ*_0_-Beta(*γ*, 2 − *α, α*)-coalescent, recall (31)) when *N* = 10^3^, *κ* = 2, *c* = 1, and *γ* as shown; black lines are 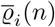 for sample size *n* as shown with rates as in (25) in Theorem 3.5, coloured lines are 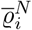 (*n*) for a sample from a population of finite size *N* evolving according to Definition 2.8 and Definition 3.3 with the potential offspring distributed as in (33) and with *ε*_*N*_ = *cN*^*α*−2^ log *N* as in (53) in Lemma 5.7; the case *γ* = 1 is compared to *ζ*(*N*) = *N* log *N*, and *γ* = 1*/*(1 + m) to *ζ*(*N*) = *N* with m as in (35) approximating 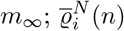 from 10^4^ experiments. The graphs are an addition to the graphs in Figure 2. The scale of the ordinate (vertical axis) may vary between graphs.

**Figure C2.**
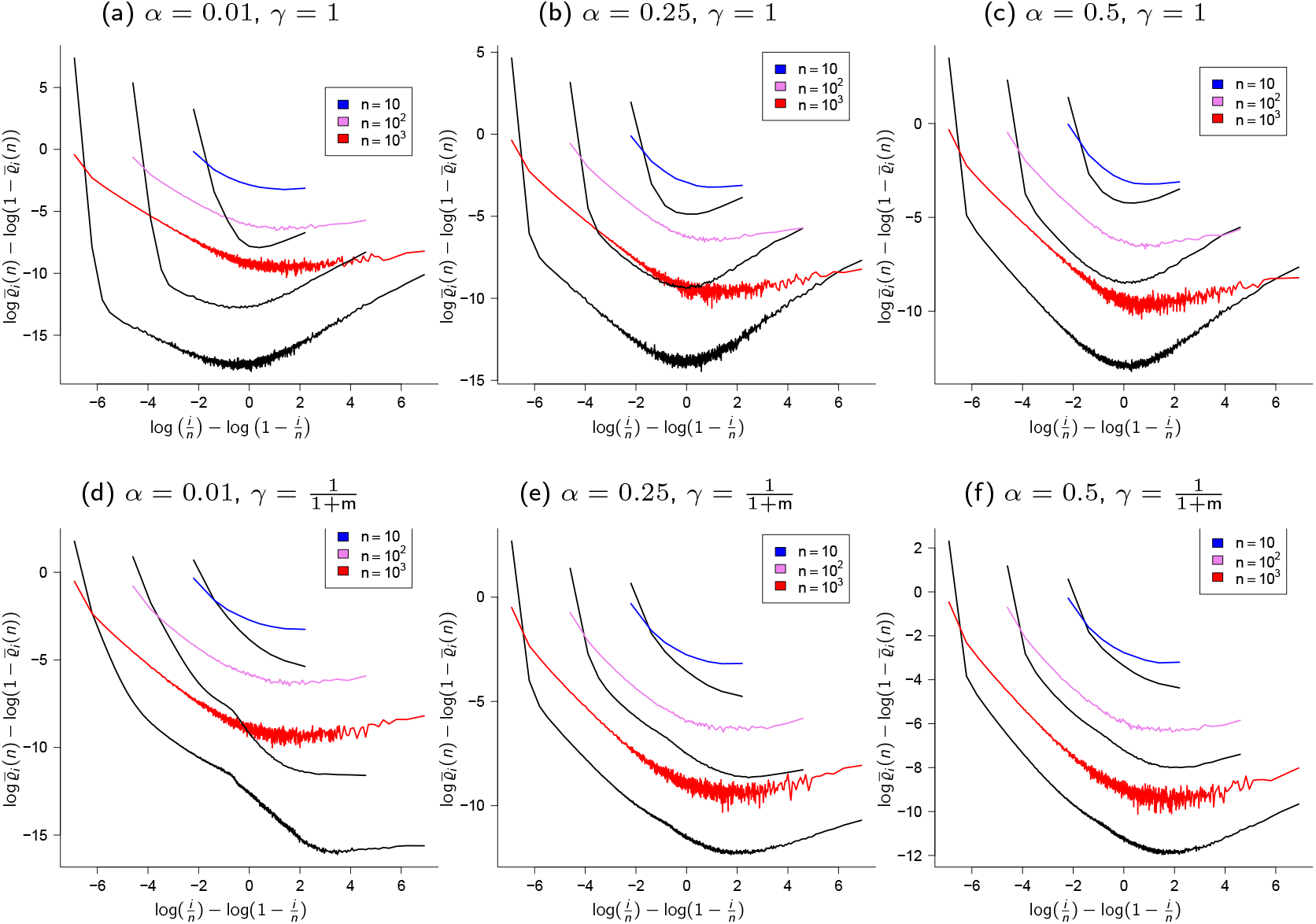
Definition 3.4 (type *B* random environment) and the *δ*_0_-Beta(*γ*, 2 − *α, α*)-coalescent. Comparing 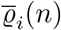 (estimates of mean relative branch lengths predicted by the *δ*_0_-Beta(*γ*, 2−*α, α*)-coalescent, black lines) and 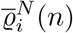 (estimates of mean relative branch lengths predicted by {*ξ*^*n,N*^} when in attraction of the *δ*_0_-Beta(*γ*, 2 − *α, α*)-coalescent, recall (31)) when *N* = 10^3^, *κ* = 2, *c* = 1, and with *α, γ*, and sampled size *n* as shown; black lines are 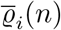 with rates as in (25), coloured lines are estimates of 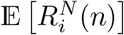 for a sample from a population evolving according to Definition 2.8 and Definition 3.4 with the potential offspring distributed as in (33) and with 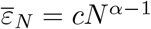 log *N* as in (68) in Lemma 5.16; the case *γ* = 1 is compared to *ζ*(*N*) = *N* log *N*, and the case *γ* = 1*/*(1 +m) compared to *ζ*(*N*) = *N* with m as in (35) approximating *m*_∞_; *ϱ*^*N*^ (*n*) from 10^4^ experiments. The graphs are an addition to Figure 3. The scale of the ordinate (vertical axis) may vary between graphs

## Appendix D Further graphs comparing 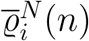 and 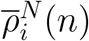

In this section we compare in Figure D1 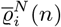 (the annealed mean relative branch lengths) and 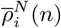 (the quenched mean relative branch lengths, recall (31)). Figure D1 is an addition to Figure 5.

## Appendix E Further graphs for the *δ*_0_-Poisson-Dirichlet(*α*, 0)-coalescent

In this section we consider in Figure E1 and Figure E2 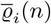 (annealed mean relative branch lengths, recall (31)) for the *δ*_0_-Poisson-Dirichlet(*α*, 0)-coalescent. In addition, in graphs E1c and E1d we compare 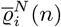 and 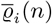. The graphs are an addition to Figure 4.

## Appendix F Further graphs for time-varying coalescents

This section contains further graphs showing the joint effects of population growth and sweepstakes reproduction on the site-frequency spectrum. Figure F1 contains graphs for the time-changed *δ*_0_-Beta(*γ*, 2 − *α, α*)-coalescent, and Figure F2 for the time-changed *δ*_0_-Poisson-Dirichlet(*α*, 0)-coalescent. Figure F1 is an addition to Figure 6, and Figure F2 to Figure 7.

**Figure D1.**
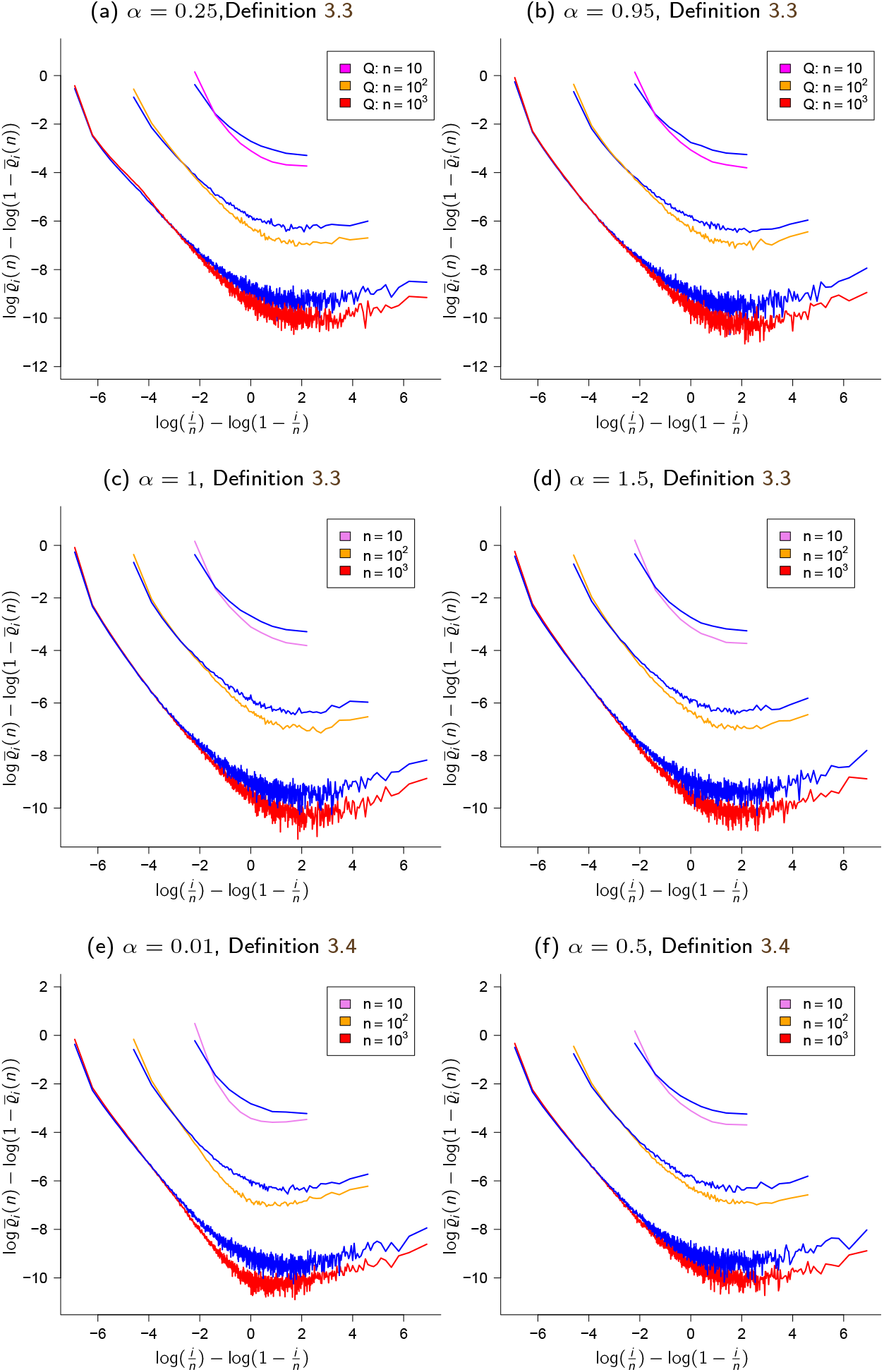
Quenched vs. annealed. Comparing 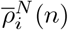 (estimates of mean relative branch lengths when conditioning on the population ancestry) and 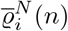 (estimates of mean relative branch lengths predicted by {*ξ*^*n,N*^}, blue lines; recall (31)) when the population evolves according to Definition 2.8 and Definition 3.3 (type *A* random environment; a,b,c,d) and Definition 3.4 (type *B* random environment; e,f); *N* = 10^3^, *α* as shown, *κ* = 2, *ζ*(*N*) = *N* ^1*/α*^ log *N* (a,b) and *ζ*(*N*) = *N* (c,d,e,f); 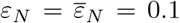 the approximations are from 10^4^ experiments. Sections § 3.2.1 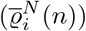 and § 3.2.2 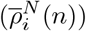 contain brief descriptions of the sampling algorithms. The scale of the ordinate (vertical axis) may vary between graphs. The graphs are an addition to Figure 5.

**Figure E1.**
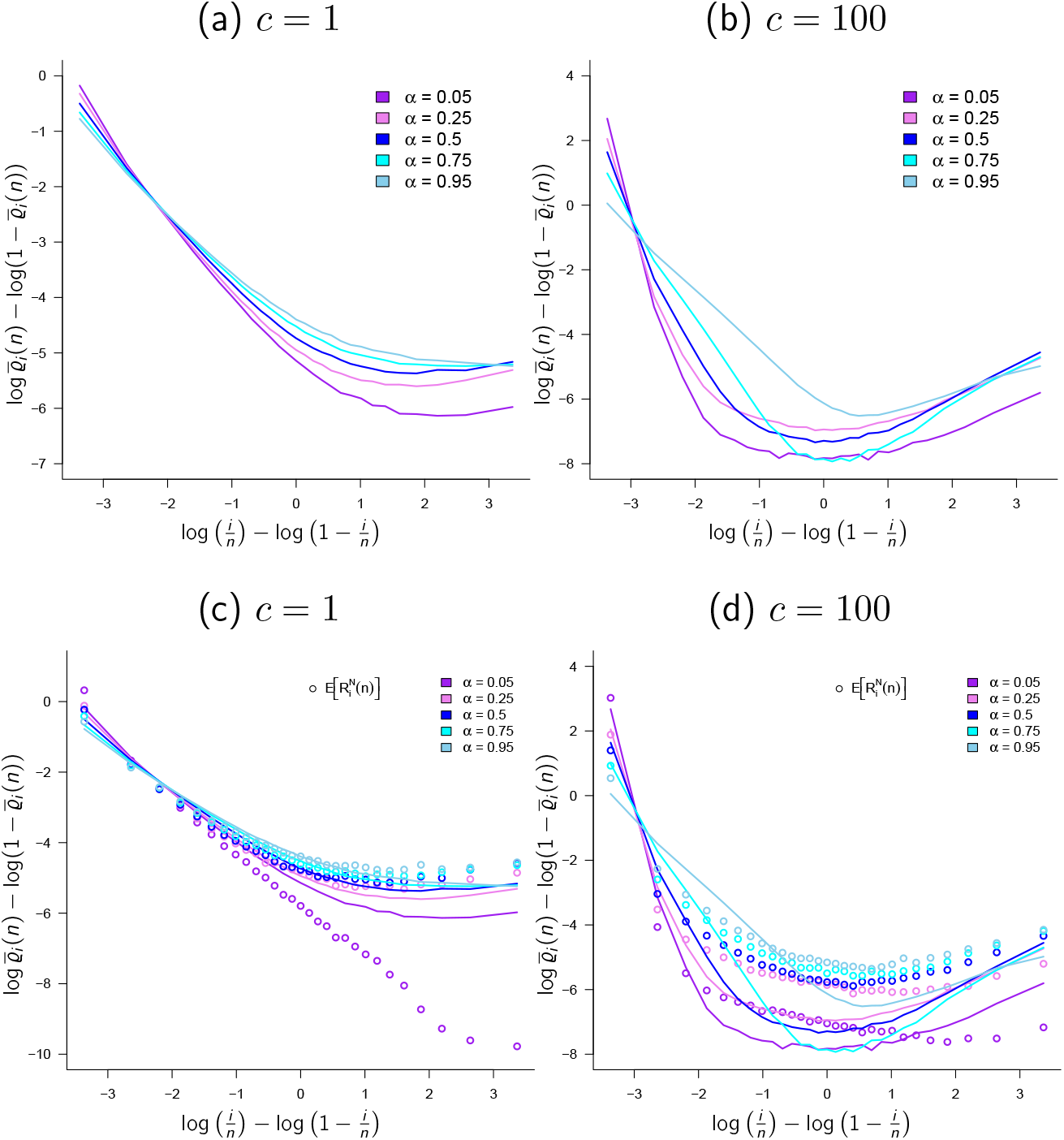
The *δ*_0_-Poisson-Dirichlet(*α*, 0)-coalescent. Approximations 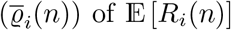 (approximations of mean relative branch lengths, recall (31); lines) predicted by the *δ*_0_-Poisson-Dirichlet(*α*, 0)-coalescent compared to 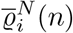 (estimates of mean relative branch lengths pre-ths predicted by {*ξ*^*n,N*^}? when in attraction of the *δ*_0_-Poisson-Dirichlet(*α*, 0)-coalescent; circles) when the population evolves according to Definition 3.3 (type A random environment) with *ζ*(*N*) = ∞, *ε* = *c*(log *N*)/*N* for *c* as shown, *N* = 3000, *α* as shown and *κ* = 2. The scale of the ordinate (*y*-axis) may vary between the graphs. See § 3.2.3 for an algorithm for sampling from the δ0-Poisson-Dirichlet(*α*, 0)-coalescent. The scale of the ordinate (vertical axis) may vary between graphs. The graphs are an addition to

**Figure E2.**
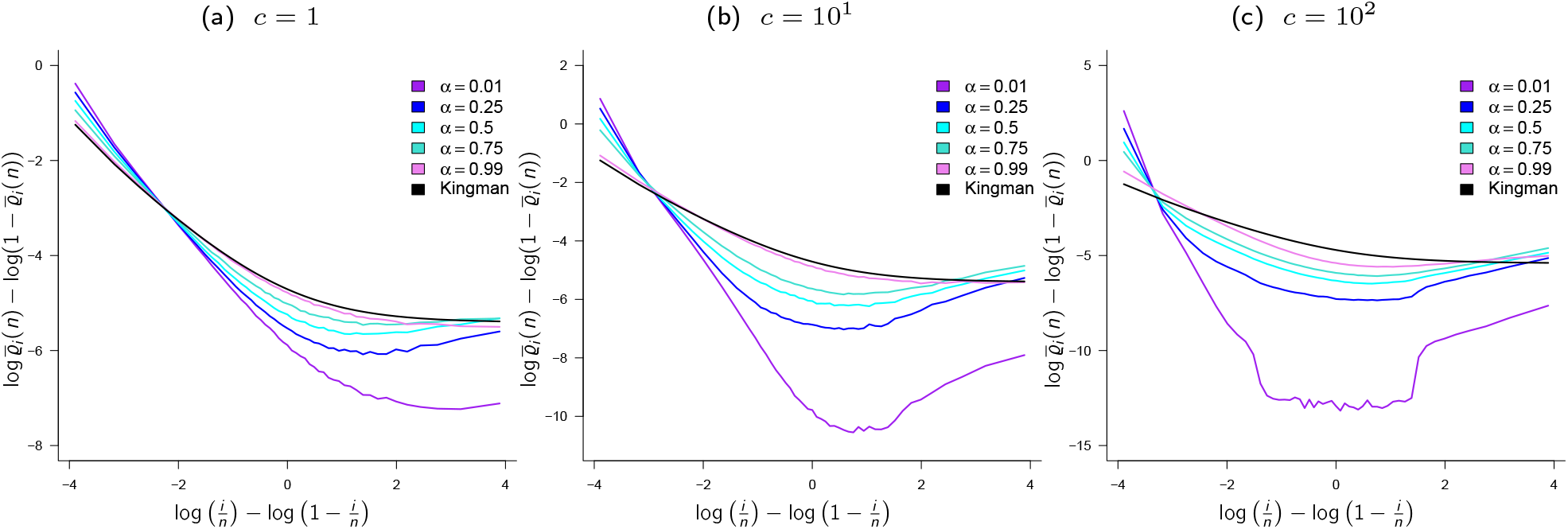
The-*δ*_0_-Poisson-Dirichlet(*α*, 0) coalescent. Examples of 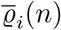 (approximations of mean relative branch lengths) (31) of E [*R*_*i*_(*n*)] predicted by the *δ*_0_-Poisson-Dirichlet(*α*, 0) coalescent (see § E) for *n* = 50, *κ* = 2, and *c* and *α* as shown. The black lines are the Kingman prediction 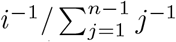 [Fu, 1995]. Results from 10^6^ experiments. The scale of the ordinate (vertical axis) may vary between graphs. The graphs are an addition to Figure 4.

**Figure F1.**
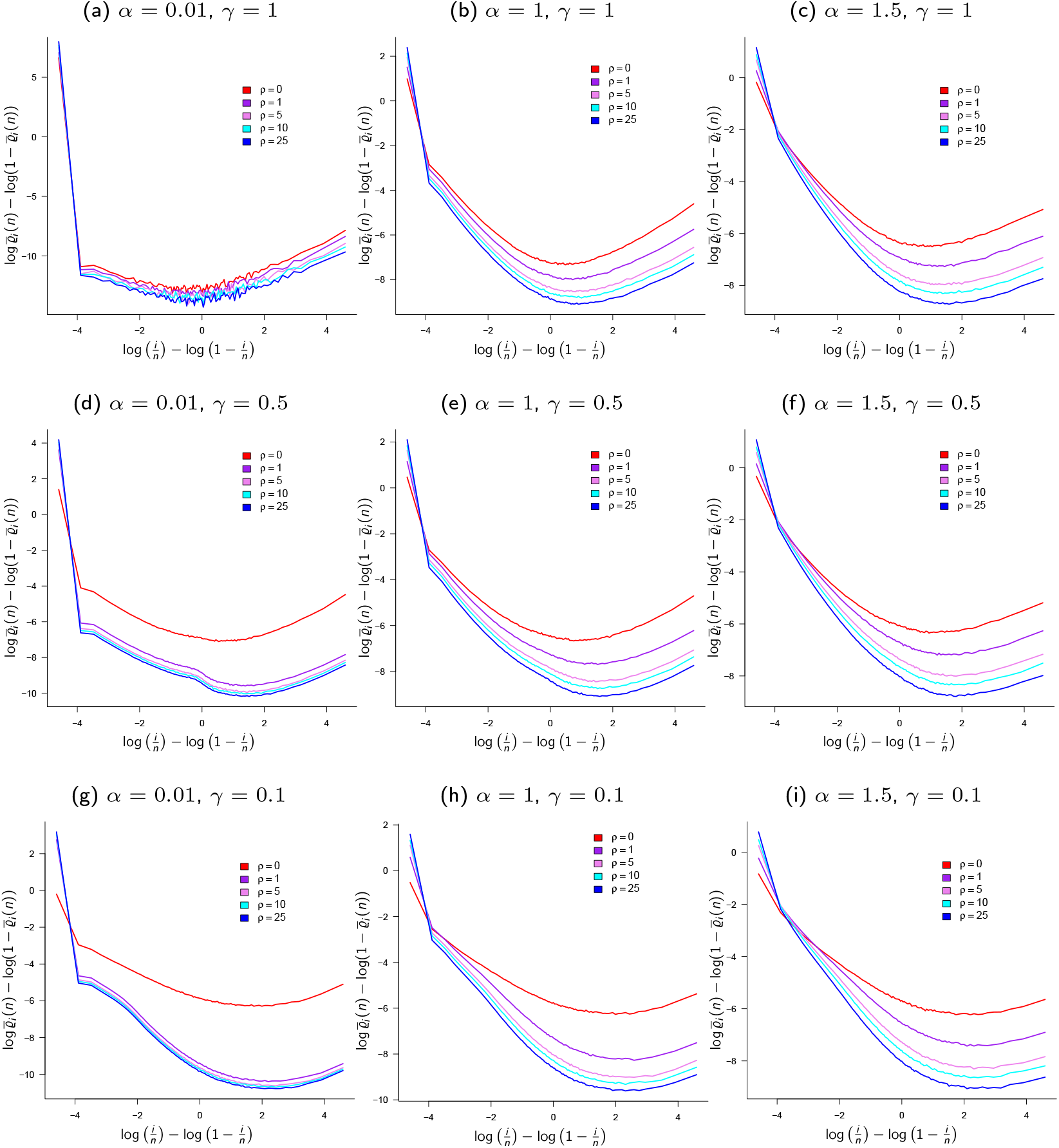
Approximations 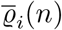 (estimates of mean relative branch lengths; recall (31)) predicted by the time-changed *δ*_0_-Beta(*γ*, 2 − *α, α*)-coalescent with time-change 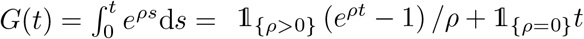 with *ρ* as shown, *κ* = 2, *c* = 1, *n* = 100. The scale of the ordinate (*y*-axis) may vary between the graphs; results from 10^5^ experiments. The scale of the ordinate (vertical axis) may vary between graphs. The graphs are an addition to Figure 6.

**Figure F2.**
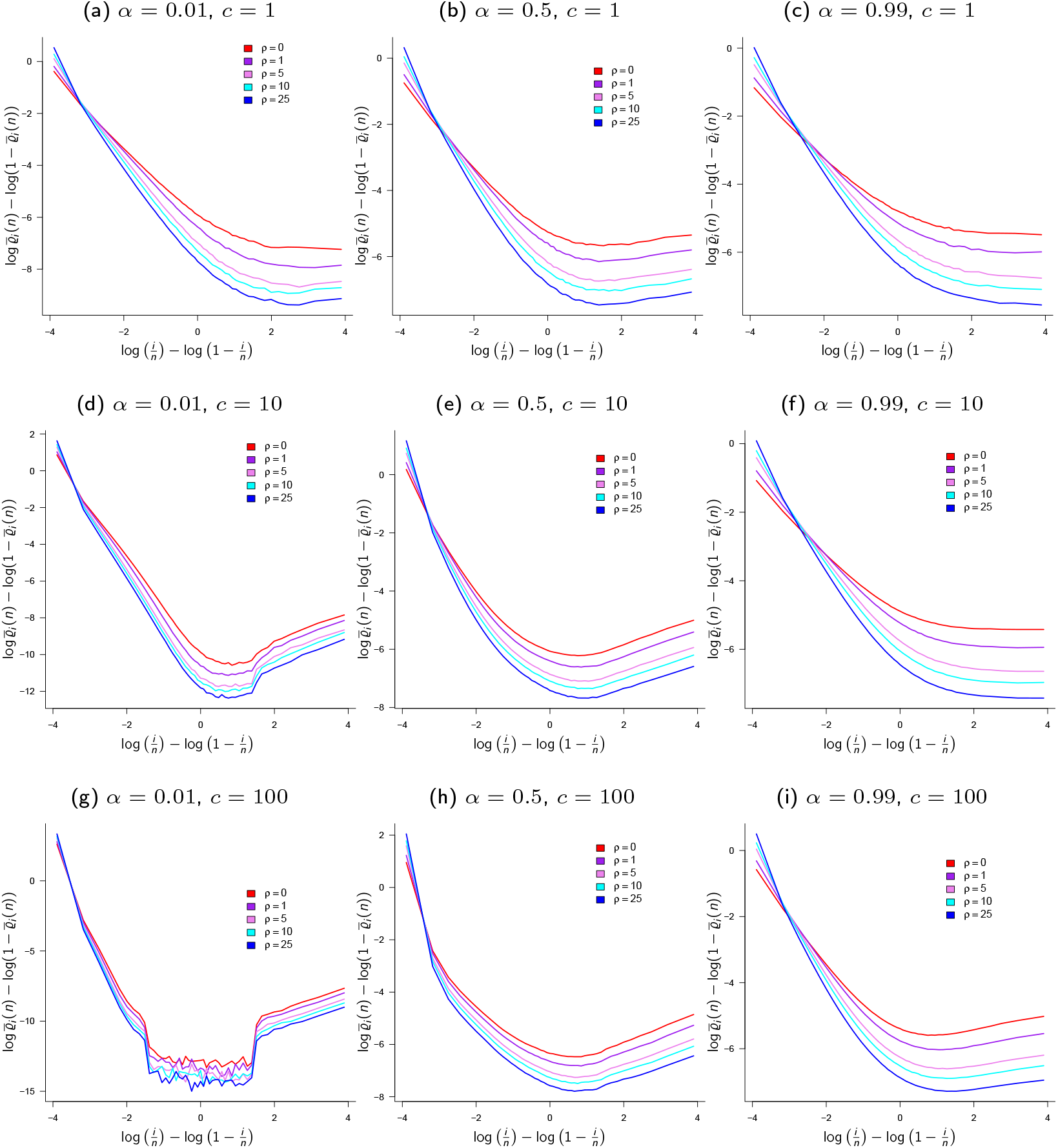
Approximations 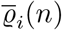 (estimates of mean relative branch lengths; recall (31)) predicted by the time-changed *δ*_0_-Poisson-Dirichlet(*α*, 0)-coalescent with time-change 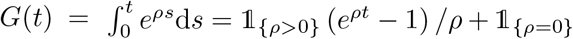 *t* with *ρ* as shown, *n* = 50, *κ* = 2; results from 10^6^ experiments. The scale of the ordinate (vertical axis) may vary between graphs. The graphs are an addition to Figure 7

## References

E Árnason. Mitochondrial cytochrome b variation in the high-fecundity Atlantic cod: trans-Atlantic clines and shallow gene genealogy. Genetics, 166:1871–1885, 2004. doi:10.1534/genetics.166.4.1871.

Einar Árnason and Katrín Halldórsdóttir. Nucleotide variation and balancing selection at the Ckma gene in Atlantic cod: analysis with multiple merger coalescent models. PeerJ, 3:e786, 2015. doi:10.7717/peerj.786.

Einar Árnason, Jere Koskela, Katrín Halldórsdóttir, and Bjarki Eldon. Sweepstakes reproductive success via pervasive and recurrent selective sweeps. eLife, 12:e80781, feb 2023. ISSN 2050-084X. doi:10.7554/eLife.80781. URL https://doi.org/10.7554/eLife.80781.

KB Athreya and SN Lahiri. Measure theory and probability theory. Springer, 2006. doi:10.1007/978-0-387-35434-7.

Sarah Barfield, Sarah W. Davies, and Mikhail V. Matz. Evidence of sweepstakes reproductive success in a broadcast-spawning coral and its implications for coral metapopulation persistence. Molecular Ecology, 32(3):696–702, 2023. doi:10.1111/mec.16774. URL https://onlinelibrary.wiley.com/doi/abs/10.1111/mec.16774.

Franz Baumdicker, Gertjan Bisschop, Daniel Goldstein, Graham Gower, Aaron P Ragsdale, Georgia Tsambos, Sha Zhu, Bjarki Eldon, E Castedo Ellerman, Jared G Galloway, Ariella L Gladstein, Gregor Gorjanc, Bing Guo, Ben Jeffery, Warren W Kretzschmar, Konrad Lohse, Michael Matschiner, Dominic Nelson, Nathaniel S Pope, Consuelo D Quinto-Cortés, Murillo F Rodrigues, Kumar Saunack, Thibaut Sellinger, Kevin Thornton, Hugo van Kemenade, Anthony W Wohns, Yan Wong, Simon Gravel, Andrew D Kern, Jere Koskela, Peter L Ralph, and Jerome Kelleher. Efficient ancestry and mutation simulation with msprime 1.0. Genetics, December 2021. doi:10.1093/genetics/iyab229. URL https://doi.org/10.1093/genetics/iyab229.

A T Beckenbach. Mitochondrial haplotype frequencies in oysters: neutral alternatives to selection models. In B Golding, editor, Non-neutral Evolution, pages 188–198. Chapman & Hall, New York, 1994. doi:10.1007/978-1-4615-2383-3.

J Berestycki, N Berestycki, and J Schweinsberg. Beta-coalescents and continuous stable random trees. Ann Probab, 35:1835–1887, 2007. doi:10.1214/009117906000001114.

J Berestycki, N Berestycki, and J Schweinsberg. Small-time behavior of beta coalescents. Ann Inst H Poincaré Probab Statist, 44:214–238, 2008. doi:10.1214/07-AIHP103.

N Berestycki. Recent progress in coalescent theory. Ensaios Mathématicos, 16:1–193, 2009. doi:10.21711/217504322009/em161.

Jean Bertoin. Random fragmentation and coagulation processes, volume 102. Cambridge University Press, 2006. doi:10.1017/CBO9780511617768.

A Bhaskar, AG Clark, and YS Song. Distortion of genealogical properties when the sample size is very large. PNAS, 111:2385–2390, 2014. doi:10.1073/pnas.1322709111.

Anand Bhaskar and Yun S. Song. Descartes’ rule of signs and the identifiability of population demographic models from genomic variation data. The Annals of Statistics, 42(6), December 2014. ISSN 0090-5364. doi:10.1214/14-aos1264. URL http://dx.doi.org/10.1214/14-AOS1264.

>M Birkner and J Blath. Measure-valued diffusions, general coalescents and population genetic inference. In J Blath, P Mörters, and M Scheutzow, editors, Trends in stochastic analysis, pages 329–363. Cambridge University Press, 2009. doi:10.1017/CBO9781139107020.

M Birkner, J Blath, M Capaldo, A M Etheridge, M Möhle, J Schweinsberg, and A Wakolbinger. Alpha-stable branching and Beta-coalescents. Electron. J. Probab, 10:303–325, 2005. doi:10.1214/EJP.v10-241.

M Birkner, J Blath, M Möhle, M Steinrücken, and J Tams. A modified lookdown construction for the Xi-Fleming-Viot process with mutation and populations with recurrent bottlenecks. ALEA Lat. Am. J. Probab. Math. Stat., 6:25–61, 2009. doi:10.34657/1856.

M Birkner, J Blath, and B Eldon. An ancestral recombination graph for diploid populations with skewed offspring distribution. Genetics, 193:255–290, 2013a. doi:10.1534/genetics.112.144329.

M Birkner, J Blath, and B Eldon. Statistical properties of the site-frequency spectrum associated with Λ-coalescents. Genetics, 195:1037–1053, 2013b. doi:10.1534/genetics.113.156612.

Matthias Birkner, Huili Liu, and Anja Sturm. Coalescent results for diploid exchangeable population models. Electronic Journal of Probability, 23(0), 2018. doi:10.1214/18-ejp175.

Matthias Birkner, Iulia Dahmer, Christina S. Diehl, and Götz Kersting. The joint fluctuations of the lengths of the Beta(2 - α, α)-coalescents. The Annals of Applied Probability, 34(1A), February 2024. ISSN 1050-5164. doi:10.1214/23-aap1964. URL http://dx.doi.org/10.1214/23-AAP1964.

Jochen Blath, Mathias Christensen Cronjäger, Bjarki Eldon, and Matthias Hammer. The site-frequency spectrum associated with Ξ-coalescents. Theoretical Population Biology, 110:36–50, aug 2016. doi:10.1016/j.tpb.2016.04.002. URL http://dx.doi.org/10.1016/j.tpb.2016.04.002.

Jonathan A Chetwynd-Diggle and Bjarki Eldon. Beta-coalescents when sample size is large. bioRxiv, January 2026. doi:10.64898/2025.12.30.697022. URL http://dx.doi.org/10.64898/2025.12.30.697022.

MARK R. Christie, DARREN W. Johnson, CHRISTOPHER D. Stallings, and MARK A. Hixon. Self-recruitment and sweepstakes reproduction amid extensive gene flow in a coral-reef fish. Molecular Ecology, 19(5):1042–1057, February 2010. ISSN 1365-294X. doi:10.1111/j.1365-294x.2010.04524.x. URL http://dx.doi.org/10.1111/j.1365-294X.2010.04524.x.

V. Chvátal. The tail of the hypergeometric distribution. Discrete Mathematics, 25(3): 285–287, 1979. doi:10.1016/0012-365x(79)90084-0. URL https://doi.org/10.1016/0012-365x(79)90084-0.

I Dahmer, G Kersting, and A Wakolbinger. The total external length of Beta-coalescents. Comb Prob Comp, 23:1010–1027, 2014. doi:10.1017/S0963548314000297.

Dimitrios Diamantidis, Wai-Tong (Louis) Fan, Matthias Birkner, and John Wakeley. Bursts of coalescence within population pedigrees whenever big families occur. GENETICS, 227 (1), February 2024. ISSN 1943-2631. doi:10.1093/genetics/iyae030. URL http://dx.doi.org/10.1093/genetics/iyae030.

P Donnelly and T G Kurtz. Particle representations for measure-valued population models. Ann Probab, 27:166–205, 1999. doi:10.1214/aop/1022677258.

Peter Donnelly and Simon Tavaré. Coalescents and genealogical structure under neutrality. Annual Review of Genetics, 29(1):401–421, December 1995. ISSN 1545-2948. doi:10.1146/annurev.ge.29.120195.002153. URL http://dx.doi.org/10.1146/annurev.ge.29.120195.002153.

Rick Durrett and Jason Schweinsberg. A coalescent model for the effect of advantageous mutations on the genealogy of a population. Stochastic Processes and their Applications, 115(10):1628–1657, October 2005. doi:10.1016/j.spa.2005.04.009.

B Eldon and J Wakeley. Coalescent processes when the distribution of offspring number among individuals is highly skewed. Genetics, 172:2621–2633, 2006. doi:10.1534/genetics.105.052175.

B Eldon, M Birkner, J Blath, and F Freund. Can the site-frequency spectrum distinguish exponential population growth from multiple-merger coalescents? Genetics, 199:841–856, 2015. doi:10.1534/genetics.114.173807.

Bjarki Eldon. Evolutionary genomics of high fecundity. Annual Review of Genetics, 54 (1):213–236, November 2020. doi:10.1146/annurev-genet-021920-095932. URL https://doi.org/10.1146/annurev-genet-021920-095932.

Bjarki Eldon. Gene genealogies in diploid populations evolving according to sweepstakes reproduction. 2026. doi:10.48550/arXiv.2601.10364. URL https://arxiv.org/abs/2601.10364.

Engen. Stochastic abundance models: with emphasis on biological communities and species diversity. Chapman-Hall, London, 1978. ISBN 94-009-5784-X. doi:10.1007/978-94-009-5784-8.

N. Etemadi. An elementary proof of the strong law of large numbers. Zeitschrift fuer Wahrscheinlichkeitstheorie und Verwandte Gebiete, 55(1):119–122, February 1981. doi:10.1007/bf01013465.

S Ethier and T Kurtz. Fleming-Viot processes in population genetics. SIAM J Control Optim, 31:345–86, 1993. doi:10.1137/0331019.

WJ Ewens. Mathematical population genetics. Springer-Verlag, Berlin, 1979. doi:10.1007/978-0-387-21822-9.

Shui Feng. The Poisson-Dirichlet distribution and related topics: models and asymptotic behaviors. Springer Science & Business Media, 2010. doi:10.1007/978-3-642-11194-5.

RA Fisher. On the dominance ratio. Proc Royal Society Edinburgh, 42:321–341, 1923. doi:10.1017/S0370164600023993.

WH Fleming and M Viot. Some measure-valued Markov processes in population genetics theory. Indiana University Mathematics Journal, 28:817–43, 1979. URL https://www.jstor.org/stable/24892583.

Fabian Freund. Cannings models, population size changes and multiple-merger coalescents. Journal of mathematical biology, 80(5):1497–1521, 2020. doi:10.1007/s00285-020-01470-5.

Fabian Freund and Arno Siri-Jégousse. The impact of genetic diversity statistics on model selection between coalescents. Computational Statistics & Data Analysis, 156:107055, 2021. doi:10.1016/j.csda.2020.107055.

Fabian Freund, Elise Kerdoncuff, Sebastian Matuszewski, Marguerite Lapierre, Marcel Hildebrandt, Jeffrey D. Jensen, Luca Ferretti, Amaury Lambert, Timothy B. Sackton, and Guillaume Achaz. Interpreting the pervasive observation of U-shaped site frequency spectra. PLOS Genetics, 19(3):e1010677, March 2023. doi:10.1371/journal.pgen.1010677. URL https://doi.org/10.1371/journal.pgen.1010677.

Y-X Fu. Statistical properties of segregating sites. Theor Popul Biol, 48:172–197, 1995. doi:10.1006/tpbi.1995.1025.

Alexander Gnedin, Alexander Iksanov, and Alexander Marynych. Λ-coalescents: a survey. Journal of Applied Probability, 51(A):23–40, December 2014a. doi:10.1239/jap/1417528464.

Alexander Gnedin, Alexander Iksanov, Alexander Marynych, and Martin Möhle. On asymptotics of the beta coalescents. Advances in Applied Probability, 46(2):496–515, June 2014b. ISSN 1475-6064. doi:10.1239/aap/1401369704. URL http://dx.doi.org/10.1239/aap/1401369704.

GH Hardy. A course of pure mathematics. Cambridge University Press, Cambridge, UK, tenth edition, 2002. doi:10.1017/CBO9780511989469.

D Hedgecock. Does variance in reproductive success limit effective population sizes of marine organisms? In A Beaumont, editor, Genetics and evolution of Aquatic Organisms, pages 1222–1344, London, 1994. Chapman and Hall. ISBN 978-0-412-49370-6.

D Hedgecock and A I Pudovkin. Sweepstakes reproductive success in highly fecund marine fish and shellfish: a review and commentary. Bull Marine Science, 87:971–1002, 2011. doi:10.5343/bms.2010.1051.

D Hedgecock, M Tracey, and K Nelson. Genetics. In L G Abele, editor, The Biology of Crustacea: Embryology, Morphology, and Genetics, volume 2, pages 297–403. Academic Press, New York, 1982. ISBN 0121064050.

D Hedgecock, S Launey, A I Pudovkin, Y Naciri, S Lapegue, and F Bonhomme. Small effective number of parents (Nb) inferred for a naturally spawned cohort of juvenile European flat oysters Ostrea edulis. Marine Biology, 150:1173–1182, 2007. doi:10.1007/s00227-006-0441-y. URL http://dx.doi.org/10.1007/s00227-006-0441-y.

R R Hudson. Properties of a neutral allele model with intragenic recombination. Theor Popul Biol, 23:183–201, 1983. doi:10.1016/0040-5809(83)90013-8.

T Huillet and M Möhle. Population genetics models with skewed fertilities: forward and backward analysis. Stoch Models, 27:521–554, 2011. doi:10.1080/15326349.2011.593411.

T Huillet and M Möhle. On the extended Moran model and its relation to coalescents with multiple collisions. Theor Popul Biol, 87:5–14, 2013. doi:10.1016/j.tpb.2011.09.004.

Jigisha Jigisha, Jeanine Ly, Nikolaos Minadakis, Fabian Freund, Lukas Kunz, Urszula Piechota, Beyhan Akin, Virgilio Balmas, Roi Ben-David, Szilvia Bencze, Salim Bourras, Matteo Bozzoli, Otilia Cotuna, Gilles Couleaud, Mónika Cséplő, Paweł Czembor Francesca Desiderio, Jost Dörnte, Antonín Dreiseitl, Angela Feechan, Agata Gadaleta, Kevin Gauthier, Angelica Giancaspro, Stefania L. Giove, Alain Handley-Cornillet, Amelia Hubbard, George Karaoglanidis, Steven Kildea, Emrah Koc, Žilvinas Liatukas, Marta S. Lopes, Fabio Mascher, Cathal McCabe, Thomas Miedaner, Fernando Martínez-Moreno, Charlotte F. Nellist, Sylwia Okoń, Coraline Praz, Javier Sánchez-Martín, Veronica Sărăţeanu, Philipp Schulz, Nathalie Schwartz, Daniele Seghetta, Ignacio Solís Martel, Agrita Švarta, Stefanos Testempasis, Dolors Villegas, Victoria Widrig, and Fabrizio Menardo. Population genomics and molecular epidemiology of wheat powdery mildew in europe. PLOS Biology, 23(5):e3003097, May 2025. ISSN 1545-7885. doi:10.1371/journal.pbio.3003097. URL http://dx.doi.org/10.1371/journal.pbio.3003097.

Mamoru Kato, Daniel A. Vasco, Ryuichi Sugino, Daichi Narushima, and Alexander Krasnitz. Sweepstake evolution revealed by population-genetic analysis of copy-number alterations in single genomes of breast cancer. Royal Society Open Science, 4(9):171060, 09 2017. ISSN 2054-5703. doi:10.1098/rsos.171060. URL https://doi.org/10.1098/rsos.171060.

Jerome Kelleher, Alison M Etheridge, and Gilean McVean. Efficient coalescent simulation and genealogical analysis for large sample sizes. PLOS Computational Biology, 12(5): e1004842, may 2016. doi:10.1371/journal.pcbi.1004842. URL http://dx.doi.org/10.1371/journal.pcbi.1004842.

M Kimura. The number of heterozygous nucleotide sites maintained in a finite population due to steady flux of mutations. Genetics, 61:893–903, 1969. doi:10.1093/genetics/61.4.893.

J. F. C. Kingman. Random discrete distributions. Journal of the Royal Statistical Society: Series B (Methodological), 37(1):1–15, September 1975. doi:10.1111/j.2517-6161.1975.tb01024.x. URL https://doi.org/10.1111/j.2517-6161.1975.tb01024.x.

J F C Kingman. On the genealogy of large populations. J App Probab, 19A:27–43, 1982a. doi:10.2307/3213548.

J.F.C. Kingman. The coalescent. Stochastic Processes and their Applications, 13(3):235–248, 1982b. ISSN 0304-4149. doi:10.1016/0304-4149(82)90011-4. URL https://www.sciencedirect.com/science/article/pii/0304414982900114.

Jere Koskela. Multi-locus data distinguishes between population growth and multiple merger coalescents. Statistical Applications in Genetics and Molecular Biology, 17(3), jun 2018. doi:10.1515/sagmb-2017-0011.

Gang Li and Dennis Hedgecock. Genetic heterogeneity, detected by PCR-SSCP, among samples of larval Pacific oysters (Crassostrea gigas) supports the hypothesis of large variance in reproductive success. Can. J. Fish. Aquat. Sci., 55(4):1025–1033, apr 1998. doi:10.1139/f97-312.

JW McCloskey. A model for the distribution of individuals by species in an environment. PhD thesis, Michigan State University, 1965. URL 10.25335/3syn-6j67.

Andrew Melfi and Divakar Viswanath. Single and simultaneous binary mergers in Wright-Fisher genealogies. Theoretical Population Biology, 121:60–71, May 2018a. doi:10.1016/j.tpb.2018.04.001.

Andrew Melfi and Divakar Viswanath. The Wright–Fisher site frequency spectrum as a perturbation of the coalescent’s. Theoretical Population Biology, 124:81–92, December 2018b. doi:10.1016/j.tpb.2018.09.005.

Fabrizio Menardo, Sébastien Gagneux, and Fabian Freund. Multiple merger genealogies in outbreaks of mycobacterium tuberculosis. Molecular Biology and Evolution, 38(1):290–306, 01 2021. ISSN 1537-1719. doi:10.1093/molbev/msaa179. URL https://doi.org/10.1093/molbev/msaa179.

Nikolaos Minadakis, Jigisha Jigisha, Luca Cornetti, Lukas Kunz, Marion C. Müller, Stefano F. F. Torriani, and Fabrizio Menardo. Genomic surveillance and molecular evolution of fungicide resistance in european populations of wheat powdery mildew. Molecular Plant Pathology, 26(3), March 2025. ISSN 1364-3703. doi:10.1111/mpp.70071. URL http://dx.doi.org/10.1111/mpp.70071.

M Möhle and S Sagitov. A classification of coalescent processes for haploid exchangeable population models. Ann Probab, 29:1547–1562, 2001. doi:10.1214/aop/1015345761.

M Möhle and S Sagitov. Coalescent patterns in diploid exchangeable population models. J Math Biol, 47:337–352, 2003. doi:10.1007/s00285-003-0218-6.

Martin Möhle. A convergence theorem for Markov chains arising in population genetics and the coalescent with selfing. Adv Appl Prob, 30:493–512, 1998. doi:10.1239/aap/1035228080.

Martin Möhle. Robustness results for the coalescent. Journal of Applied Probability, 35(2): 438–447, 1998. doi:10.1017/S0021900200015060.

Martin Möhle. Total variation distances and rates of convergence for ancestral coalescent processes in exchangeable population models. Advances in Applied Probability, pages 983–993, 2000. doi:10.1017/S0001867800010417.

Simon Myers, Charles Fefferman, and Nick Patterson. Can one learn history from the allelic spectrum? Theoretical Population Biology, 73(3):342–348, May 2008. ISSN 0040-5809. doi:10.1016/j.tpb.2008.01.001. URL http://dx.doi.org/10.1016/j.tpb.2008.01.001.

Hiro-Sato Niwa, Kazuya Nashida, and Takashi Yanagimoto. Reproductive skew in japanese sardine inferred from DNA sequences. ICES Journal of Marine Science: Journal du Conseil, 73(9):2181–2189, may 2016. doi:10.1093/icesjms/fsw070. URL http://dx.doi.org/10.1093/icesjms/fsw070.

Vladimir Panov. Limit theorems for sums of random variables with mixture distribution. Statistics & Probability Letters, 129:379–386, 2017. ISSN 0167-7152. doi:10.1016/j.spl.2017.06.017. URL https://www.sciencedirect.com/science/article/pii/S0167715217302213.

J Pitman. Coalescents with multiple collisions. Ann Probab, 27:1870–1902, 1999. doi:10.1214/aop/1022874819.

Jim Pitman. Exchangeable and partially exchangeable random partitions. Probability theory and related fields, 102(2):145–158, 1995. doi:10.1007/BF01213386.

R Core Team. R: A Language and Environment for Statistical Computing. R Foundation for Statistical Computing, Vienna, Austria, 2026. URL https://www.R-project.org/.

Zvi Rosen, Anand Bhaskar, Sebastien Roch, and Yun S Song. Geometry of the sample frequency spectrum and the perils of demographic inference. Genetics, 210(2):665–682, July 2018. ISSN 1943-2631. doi:10.1534/genetics.118.300733. URL http://dx.doi.org/10.1534/genetics.118.300733.

S Sagitov. The general coalescent with asynchronous mergers of ancestral lines. J Appl Probab, 36:1116–1125, 1999. doi:10.1239/jap/1032374759.

S Sagitov. Convergence to the coalescent with simultaneous multiple mergers. J Appl Probab, 40:839–854, 2003. doi:10.1239/jap/1067436085.

O Sargsyan.and J Wakeley. A coalescent process with simultaneous multiple mergers for approximating the gene genealogies of many marine organisms. Theor Pop Biol, 74: 104–114, 2008. doi:10.1016/j.tpb.2008.04.009.

J Schweinsberg. Coalescents with simultaneous multiple collisions. Electron J Probab, 5: 1–50, 2000. doi:10.1214/EJP.v5-68.

J Schweinsberg. Coalescent processes obtained from supercritical Galton-Watson processes. Stoch Proc Appl, 106:107–139, 2003. doi:10.1016/S0304-4149(03)00028-0.

J Schweinsberg.and R Durrett. Random partitions approximating the coalescence of lineages during a selective sweep. Ann Appl Probab, 1591–1651, 2005. doi:10.1214/105051605000000430.

F Tajima. Evolutionary relationships of DNA sequences in finite populations. Genetics, 105:437–460, 1983. doi:10.1093/genetics/105.2.437.

O Tange. GNU parallel – the command-line power tool. The USENIX Magazine, 2011.

David L. J. Vendrami, Lloyd S. Peck, Melody S. Clark, Bjarki Eldon, Michael Meredith, and Joseph I. Hoffman. Sweepstake reproductive success and collective dispersal produce chaotic genetic patchiness in a broadcast spawner. Science Advances, 7(37), September 2021. ISSN 2375-2548. doi:10.1126/sciadv.abj4713. URL http://dx.doi.org/10.1126/sciadv.abj4713.

Bengt von Bahr and Carl-Gustav Esseen. Inequalities for the rth Absolute Moment of a Sum of Random Variables, 1 ≦ r ≦ 2. The Annals of Mathematical Statistics, 36(1):299 –303, 1965. doi:10.1214/aoms/1177700291. URL https://doi.org/10.1214/aoms/1177700291.

J Wakeley.and T Takahashi. Gene genealogies when the sample size exceeds the effective size of the population. Mol Biol Evol, 20:208–2013, 2003. doi:10.1093/molbev/msg024.

John Wakeley. Coalescent Theory: An Introduction. Roberts & Company Publishers, Greenwood Village, 2009. ISBN 0-9747077-5-9.

John Wakeley, Léandra King, Bobbi S Low, and Sohini Ramachandran. Gene genealogies within a fixed pedigree, and the robustness of Kingman’s coalescent. Genetics, 190(4): 1433–1445, 2012. doi:10.1534/genetics.111.135574.

R. S. Waples. Tiny estimates of the Ne/N ratio in marine fishes: Are they real? Journal of Fish Biology, 89(6):2479–2504, 2016. ISSN 1095-8649. doi:10.1111/jfb.13143.

G A Watterson. On the number of segregating sites in genetical models without recombination. Theor Pop Biol, 7:256–276, 1975. doi:10.1016/0040-5809(75)90020-9.

Sewall Wright. Evolution in mendelian populations. Genetics, 16(2):97–159, 03 1931. ISSN 1943-2631. doi:10.1093/genetics/16.2.97. URL https://doi.org/10.1093/genetics/16.2.97.

